# A heterogeneous pool of tumor precursor cells with self-replenishing and tumor-originating properties in early pathogenesis of Chronic Lymphocytic Leukemia

**DOI:** 10.1101/2022.05.04.490383

**Authors:** Bettina Budeus, Maria Dampmann, Artur Kibler, Bianca Faßbender, Martina Brauser, Kevin Bronischewski, Ekaterina Homp, Sarah Taudien, Patricia Johansson, Johannes Bloehdorn, Stephan Stilgenbauer, Julia von Tresckow, Jan Dürig, Hans Christian Reinhardt, Ralf Küppers, Sascha Dietrich, Marc Seifert

**Affiliations:** Institute of Cell Biology (Cancer Research), Medical Faculty, University of Duisburg-Essen, Essen, Germany; Department of Hematology and Stem Cell Transplantation, Medical Faculty, University of Duisburg-Essen, Essen, Germany; Department of Haematology, Oncology and Clinical Immunology, University Hospital Düsseldorf, Düsseldorf, Germany; Department of Internal Medicine III, University of Ulm, Ulm, Germany; German Cancer Consortium (DKTK); Center for Integrated Oncology Aachen-Bonn-Cologne-Düsseldorf (CIO ABCD), Aachen, Bonn, Cologne, Düsseldorf, Germany

## Abstract

Intratumor heterogeneity (ITH) refers to the coexistence of distinct cancer cell subpopulations within a single tumor, each with unique molecular and functional properties. Understanding the dynamics and evolution of ITH is crucial for predicting tumor progression and the development of therapy resistance.

Here, we conducted a comprehensive investigation of ITH in Chronic Lymphocytic Leukemia (CLL) and identified a subpopulation that carried the CLL B-cell receptor rearrangement but contrasted from the CLL main population by low CD5 and high CD20 expression and highly mutated Ig-genes. This CLL-related memory B-cell population shared somatic driver mutations with the main CLL population but also exhibited exclusive somatic mutations. Phylogenetic analysis suggested a pathogenically early generation of these CLL-related B cells before monoclonal B lymphocytosis or CLL manifestation. Our data indicated that CLL-related B cells have self-replenishing potential, as they diminish upon treatment but recover indistinguishably on relapse. This contrasts with the main CLL population, which mostly represents a selective and novel expansion of the CLL-related B cells. This differentiation capacity into conventional CLL cells, and the expression of leukemic stem cell signatures further supported their tumorigenic capacity.

We propose that these CLL-related B cells represent a pool of highly diversified, early-stage CLL precursor cells, which persist in the shape of “malignant memory B cells”. Longitudinal analyses of these CLL precursor cells suggested that they form a reservoir of malignant, leukemia-originating cells which contribute during disease progression to CLL outgrowth and clonal evolution.

## Introduction

Despite increasing knowledge on intra-tumor heterogeneity (ITH) and tumor-originating cells with stem cell properties in the evolution of leukemias and solid tumors,^1–3^ the existence of lymphoma-originating cells remain controversial.^4–6^ This is of particular interest in clinically heterogenous malignancies such as chronic lymphocytic leukemia (CLL), where the prediction of clonal evolution dynamics is difficult.^7,8^ Multiple factors affect CLL outcome, including the tumor microenvironment,^9,10^ and the recirculation of tumor cells between periphery and lymphoid tissue.^11^ Events impacting the CLL course can occur early in pathogenesis, including CLL mutations in hematopoietic precursors, the equipment with B cell receptor (BCR) specificities,^12^ chromosomal aberrations,^13,14^ the cellular origins,^15^ B cell maturation intensity,^16,17^ and mutations present already in the asymptomatic precursor state monoclonal B cell lymphocytosis (MBL).^18–20^

It is widely accepted that Ig-unmutated CLL (U-CLL) cases derive from T cell-independently activated B cells, whereas IgV-mutated CLL (M-CLL) cases derive from germinal center (GC)-experienced B cells. These different origins affect clinical outcome.^15,17,21,22^ CLL is known to be composed of tumor subpopulations associated with proliferation centers in the tissue,^10,23^ and tumor states associated with tissue egress or recirculation.^24^ However, there is little information whether distinct lymphoma-originating subpopulations exist and contribute to CLL evolution and resistance devlopment.^11,24,25^ The evolution of genomic diversity in CLL has been deeply investigated.^13,18,26^ Somatic CLL mutations occur already in hematopoietic precursor cells,^19^ and in monoclonal B cell lymphocytosis (MBL).^20^ It Is debated whether MBL progression occurs through ongoing diversification and outgrowth of selected clones, or rather selective expansion of subclones of an a-priori heterogeneous population.^26,27^ Both ideas are supported by high-throughput analyses of genetic lesions^7,28,29^ and mitochondrial mutations in CLL.^30^ Notably, CLL outcome is often shaped by genetic lesions that were already present in earlier, slow-growing stages,^31^ hence, there is a clinical need to understand early tumor stage diversity and evolution.

Our study identified a novel CLL subpopulation, hidden within the CD20^high^ and CD5^low^ B cell pool and distinct from the commonly observed CD20^low^ and CD5^high^CLL population. These CLL-related B cells share the CLL-VDJ rearrangement and CLL drive mutations but also differ in their genomic landscape, gene expression, and functional capacity. They represent an intermediate stage between antigen-experienced non-malignant B cells and fully transformed CLL cells, bearing signatures of tumor stem cells. These CLL precursor cells develop and diversify early in CLL pathogenesis, persist, and expand during disease progression or relapse. According to IgV-mutation phylogenetics, they contribute to the evolution of previously absent or underrepresented CLL subpopulations. We provide evidence that these CLL precursor cells possess tumor-originating capacity and hold crucial information for understanding the pathogenesis and clonal evolution of CLL.

## Results

### Discovery of a transitional CD20^high^/CD5^low^ CLL-related B cell subpopulation

The predominant population of CLL cells in patient peripheral blood (PB) is typically marked by CD20^low^ and CD5^high^ expression, in contrast to the minority of CD20^high^ and CD5^low^ non-malignant residual B-cells (NRB). We measured the frequency of NRB cells in 70 CLL cases, which ranged from 0.05% to 4.75% (Tables 1 and 2), (see Figures 1A-C, S1A-F).

**Figure 1:**
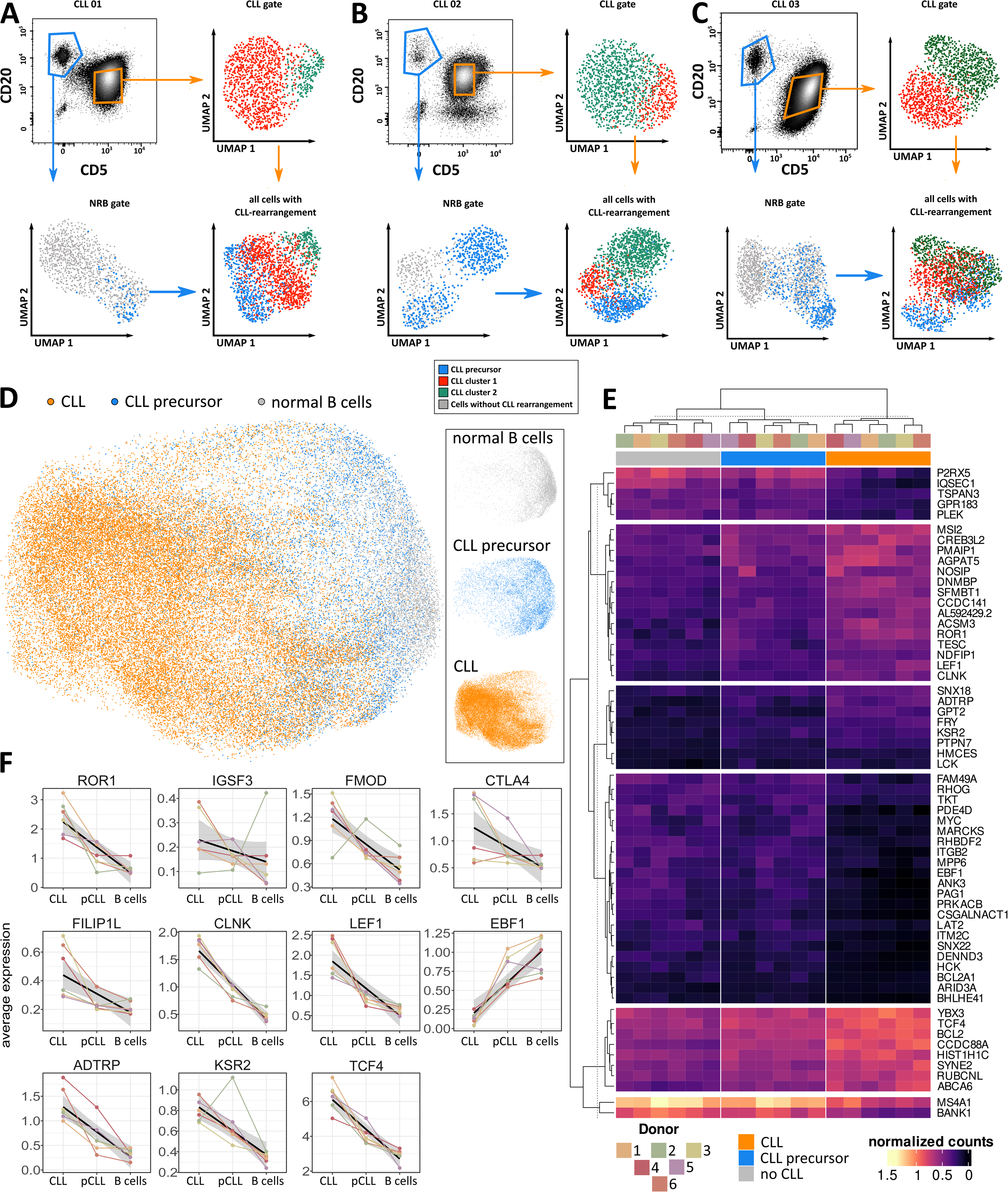
Single-cell RNA/VDJseq analysis of circulating CLL cells. **(A-C)** The cell sorting strategy for CLL cells (orange) and normal B cells (blue) from three out of six cases (from left to right CLL 01 - 03) is shown in the top left plots. Top right plots: UMAP of scRNA profiles from the CLL gate (orange), showing two CLL clusters (red and green), all of which express the known CLL-IgH/IgL rearrangement. Lower left plots: UMAP of scRNA profiles from the normal B cell gate (blue), showing one cluster of normal B cells with polyclonal IgH/IgL rearrangements (grey) and one cluster of normal-associated CLL cells expressing the known CLL-IgH/IgL rearrangement (blue). Lower right plots: integrated UMAP and clustering from all cells expressing the known CLL-IgH/IgL rearrangement (red, green, and blue). Blue events colocalizing with CLL clusters (red/green) likely represent cell-sorting contaminations. **D)** Integrated UMAP and clustering analysis of scRNA/VDJseq data from six CLL cases (CLL 01-06). CLL cells are shown in orange, CLL precursor cells in blue, and polyclonal B cells in grey. **E)** Heatmap of the top differentially expressed genes between CLL, CLL precursor and normal B cells. **F)** Average expression values of typical CLL genes in CLL cells, CLL precursor cells and normal B cells from six donors, paired samples are connected by line.

**Table 1:** CLL patient information.

|  |  |  |
| --- | --- | --- |
| **Number of CLL patients** |  | 70 |
| **Age** | *Mean ± SD* | 68.1 ± 14.5 |
|  | *Range* | 37 - 89 |
| **Gender** | *Male* | 48 |
|  | *Female* | 22 |
| **IGHV mutation status** | *Mutated* | 27 |
|  | *Unmutated* | 19 |
|  | *n/a* | 24 |
| **Binet stage (at sampling)** | *A* | 50 |
|  | *B* | 12 |
|  | *C* | 8 |
| **High risk cytogenetics** | *Del17p or del 11q* | 13 |
|  | *No del17p/del11q* | 39 |
|  | *n/a* | 18 |
| **Lymphocyte doubling time** | *< 12 months* | 18 |
|  | *> 12 months* | 41 |
|  | *n/a* | 11 |
| **Time to first treatment** | *Patients with treatment* | 26 |
|  | *Median (months)* | 34.7 |
|  | *Range (months)* | 0 - 131 |
| **Follow up** | *Median (months)* | 67 |
|  | *Range (months)* | 0 - 278 |

n/a, not available

**Table 2:**
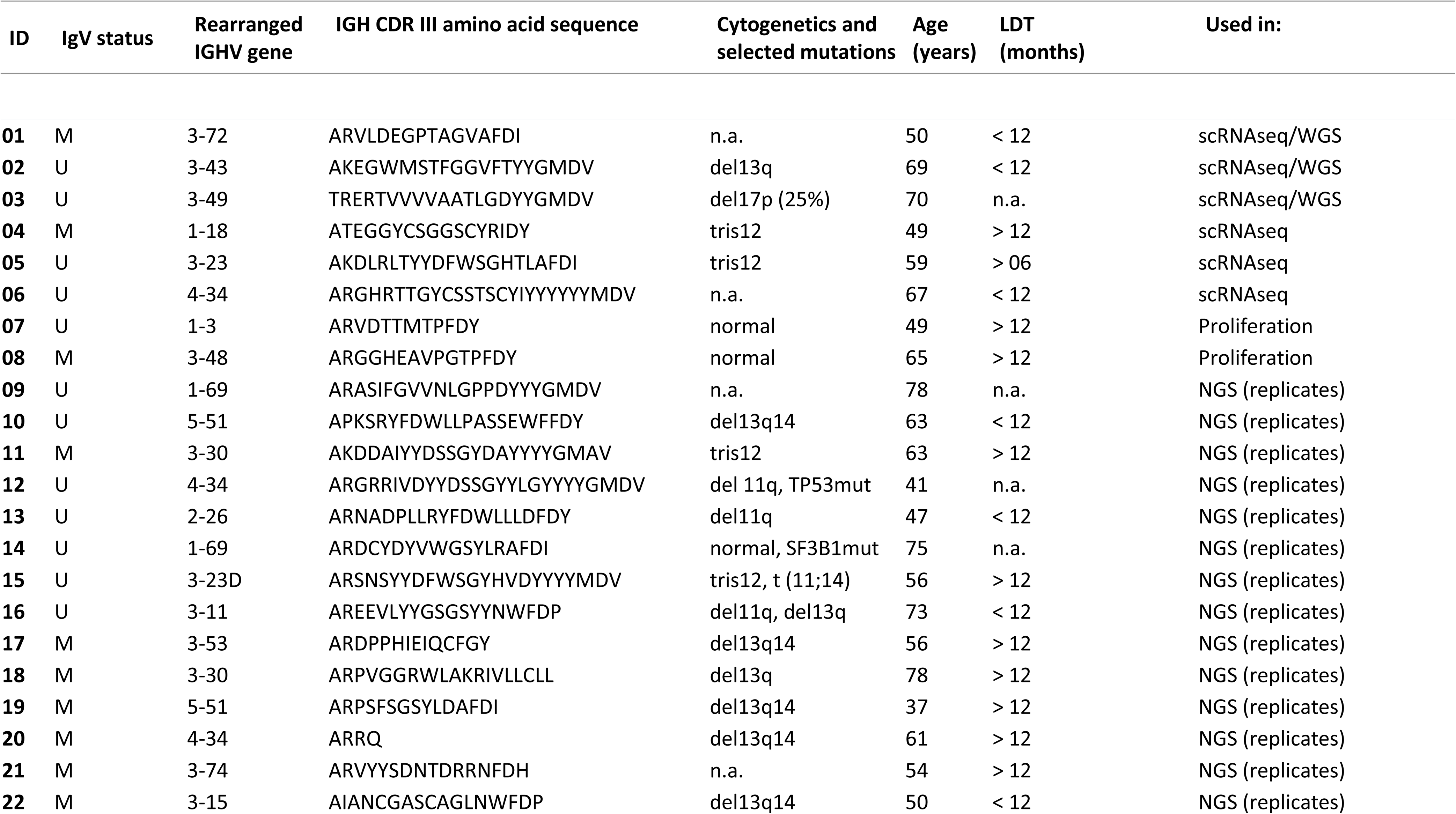
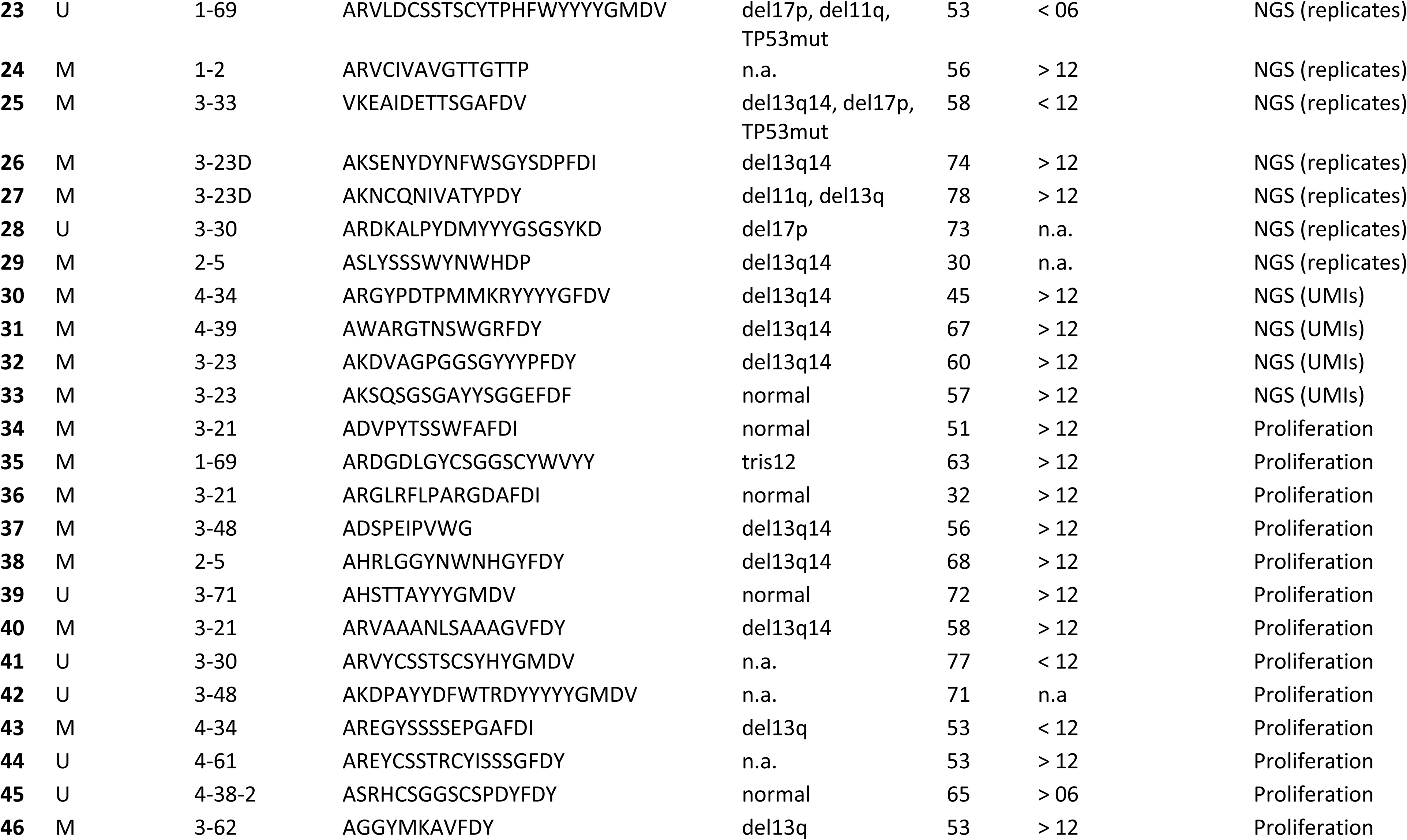
Characteristics of the CLL cohort with BCR repertoire sequencing. IGHV mutation status was determined by IGHV gene sequencing at diagnosis and confirmed by IGHV repertoire deep sequencing. Cases with an average IGHV mutation frequency < 2% were regarded as uCLL. Cytogenetic abnormalities, age in years at diagnosis (age), and lymphocyte doubling time (LDT) determined during the first 12 months after diagnosis are given. Unavailable information is denoted as n.a., not detected as n.d..

We isolated these NRB cells and their corresponding CLL cells from six selected CLL cases (3 M-CLL, 3 U-CLL, Table 2) and investigated their intra-tumor heterogeneity (ITH) by single-cell RNA- and VDJ-sequencing (scRNA/VDJseq, Figures 1A-C, S1A-C and Table 3). Equal numbers of cells from both subpopulations were isolated by cell sorting, resulting in an enrichment of the NRB cells by a factor of 25 to 100. Individual barcodes were assigned to each subpopulation during scRNA/VDJseq library generation.

**Table 3:**
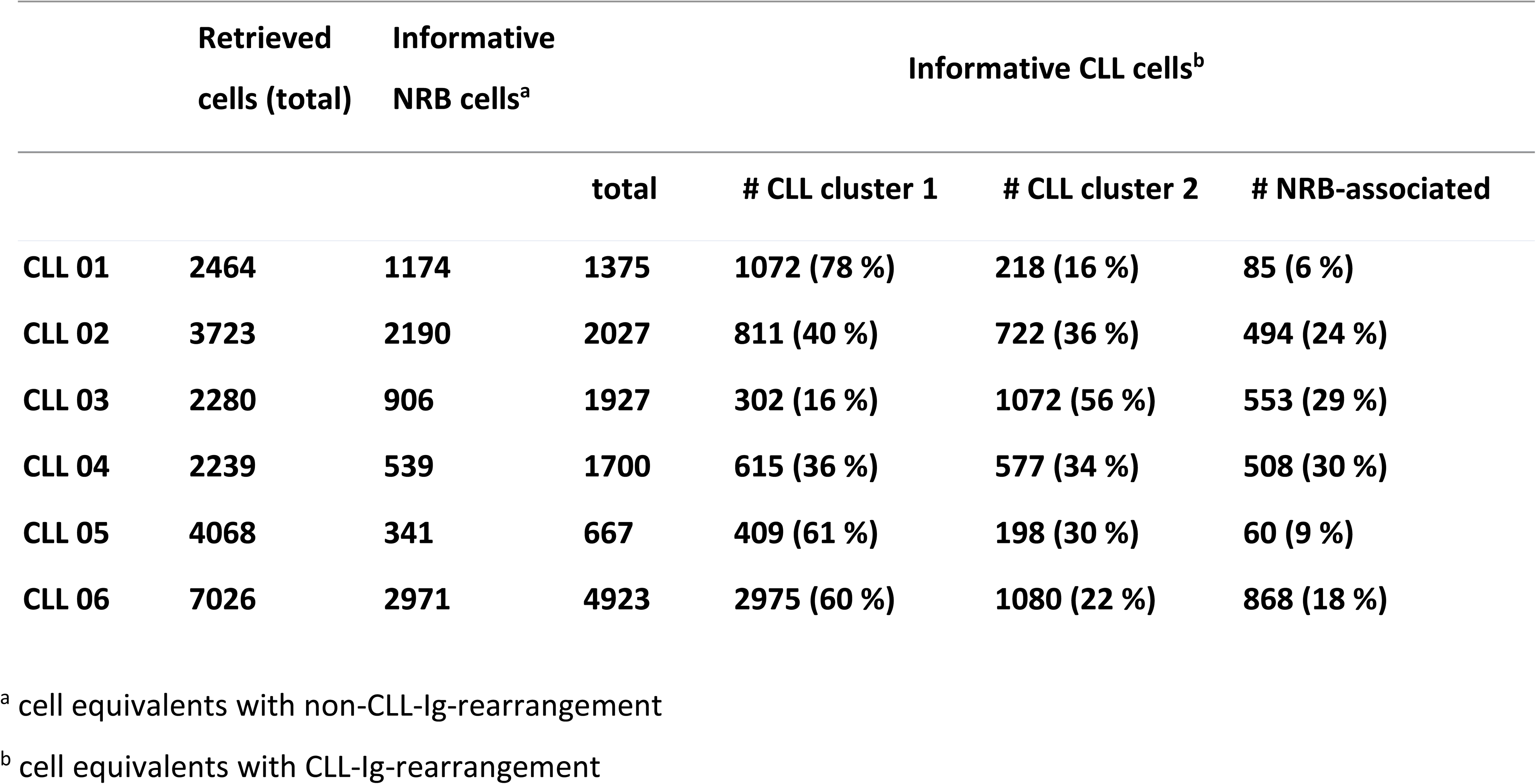
Cell equivalents retrieved from scRNA-seq/VDJ-seq profiles.

|  | Retrieved<br>cells (total) | Informative<br>NRB cells<sup>a</sup> | Informative CLL cells<sup>b</sup> |  |  |  |
| --- | --- | --- | --- | --- | --- | --- |
|  |  |  | total | # CLL cluster 1 | # CLL cluster 2 | # NRB-associated |
| CLL 01 | 2464 | 1174 | 1375 | 1072 (78 %) | 218 (16 %) | 85 (6 %) |
| CLL 02 | 3723 | 2190 | 2027 | 811 (40 %) | 722 (36 %) | 494 (24 %) |
| CLL 03 | 2280 | 906 | 1927 | 302 (16 %) | 1072 (56 %) | 553 (29 %) |
| CLL 04 | 2239 | 539 | 1700 | 615 (36 %) | 577 (34 %) | 508 (30 %) |
| CLL 05 | 4068 | 341 | 667 | 409 (61 %) | 198 (30 %) | 60 (9 %) |
| CLL 06 | 7026 | 2971 | 4923 | 2975 (60 %) | 1080 (22 %) | 868 (18 %) |

<sup>a</sup> cell equivalents with non-CLL-Ig-rearrangement
<sup>b</sup> cell equivalents with CLL-Ig-rearrangement

**Table 4:** Treatment of patients included in Figure S5 and S4.

| Patient | Time | Therapy |
| --- | --- | --- |
| 10 | 2008 | Chemo-Immunotherapy |
| 10 | 2013 | Chemo-Immunotherapy |
| 12 | 2007-2008 | Chemo-Immunotherapy |
| 12 | 2011-2013 | Immunomodulatory therapy |
| 12 | 2014 | Chemo-Immunotherapy + BTK-Inhibitor |
| 12 | 2014-2018 | BTK-Inhibitor |
| 13 | 2010 | Chemo-Immunotherapy |
| 13 | 2012-2013 | Chemo-Immunotherapy |
| 13 | 2016-2017 | Chemo-Immunotherapy |
| 25 | 2006 | Chemotherapy |
| 25 | 2017 | Chemo-Immunotherapy |
| 25 | 2017-2018 | BTK-Inhibitor |
| 28 | 2013-2014 | Chemo-Immunotherapy |
| 28 | 2014 | Radiotherapy |
| 28 | 2014-2015 | BCL2 Inhibitor |
| 28 | 2016-2018 | BTK-Inhibitor + anti-CD20 antibody |
| 28 | 2018 | PI3K-Inhibitor |
| **28** | 2018 | Immunomodulatory therapy |
| **29** | 2010 | Chemo-Immunotherapy |

To uncover additional substructures within each population, we conducted separate analyses on the CD20^low^/CD5^high^ CLL cells and the CD20^high^/CD5^low^ NRB cells using Uniform Manifold Approximation and Projection (UMAP) and clustering. Within CD20^low^/CD5^high^ CLL cells, we discerned two distinct subpopulations, consistent with findings from earlier studies (Figure S2A).^24^ The CD20^high^/CD5^low^ NRB population consisted of both naïve and memory B cells, as confirmed by flow cytometry (Figure S1E), and contained polyclonal *IGH*- and/or *IGL*-rearrangements (Figure S2A). Surprisingly, among CD20^high^/CD5^low^ NRB cells, a fraction of 2-20% carried the known CLL-IgV-rearrangement (Table 2,3 Figure 1). Despite sharing the same CLL-IgV-rearrangement, these CD20^high^/CD5^low^ CLL-related B cells were characterized by a unique scRNAseq signature, which was distinct from NRB cells and the CD20^low^/CD5^high^ main CLL population (Figure 1A-C lower panels). The CLL-related CD20^high^/CD5^low^ population was identified in all three CLL patients, exhibiting a consistent expression profile across each case. (Figure 1D, and Table 3). CLL-related CD20^high^/CD5^low^ B cells represent a transcriptional “intermediate” state between NRB cells and the main CLL population. These cells exhibited normal B cell signatures alongside elevated levels of genes highly expressed in both CLL cells and the premalignant precursor monoclonal B cell lymphocytosis (MBL) (Figure 1 D-F).

We discovered a previously unrecognized CD20^high^/CD5^low^ CLL-related subpopulation, which is characterized by the same CLL-IgV-rearrangement as the CD20^low^/CD5^high^ main CLL population and transcriptionally resembles a transitional state between non-malignant and MBL/CLL B cells.

### Genetic relationship and divergence of CD20^high^/CD5**^l^**^ow^ CLL-related and CD20^low^/CD5^high^ CLL cells

We compared the mutational landscape of CD20^high^/CD5^low^ CLL-related B cells and the main CLL population to ascertain their genetic relationship using whole genome sequencing of three cases (1 M-CLL, 2 U-CLL) and germline controls (T-cells). We detected on average 196 non-synonymous mutations per case (variant allele frequency, VAF > 15%). An average of 13% (range 10% - 16%) of somatic variants were shared between the CD20^high^/CD5^low^ CLL-related cells and the main CLL population (Figure 2A-C), confirming their clonal relationship. However, most mutations (average 87%) were exclusively detected in either CLL population which included recurrent somatic CLL mutations (e.g. SF3B1, SETD2, USP8, Figure 2A-C, red).^32^ The frequent presence of CLL driver mutations, and point mutations in *hTERT*, *ATM*, and *ATRX* in CD20^high^/CD5^low^ CLL-related cells in 3/3 cases indicated a definitive malignant determination of CLL-related B cells.

**Figure 2:**
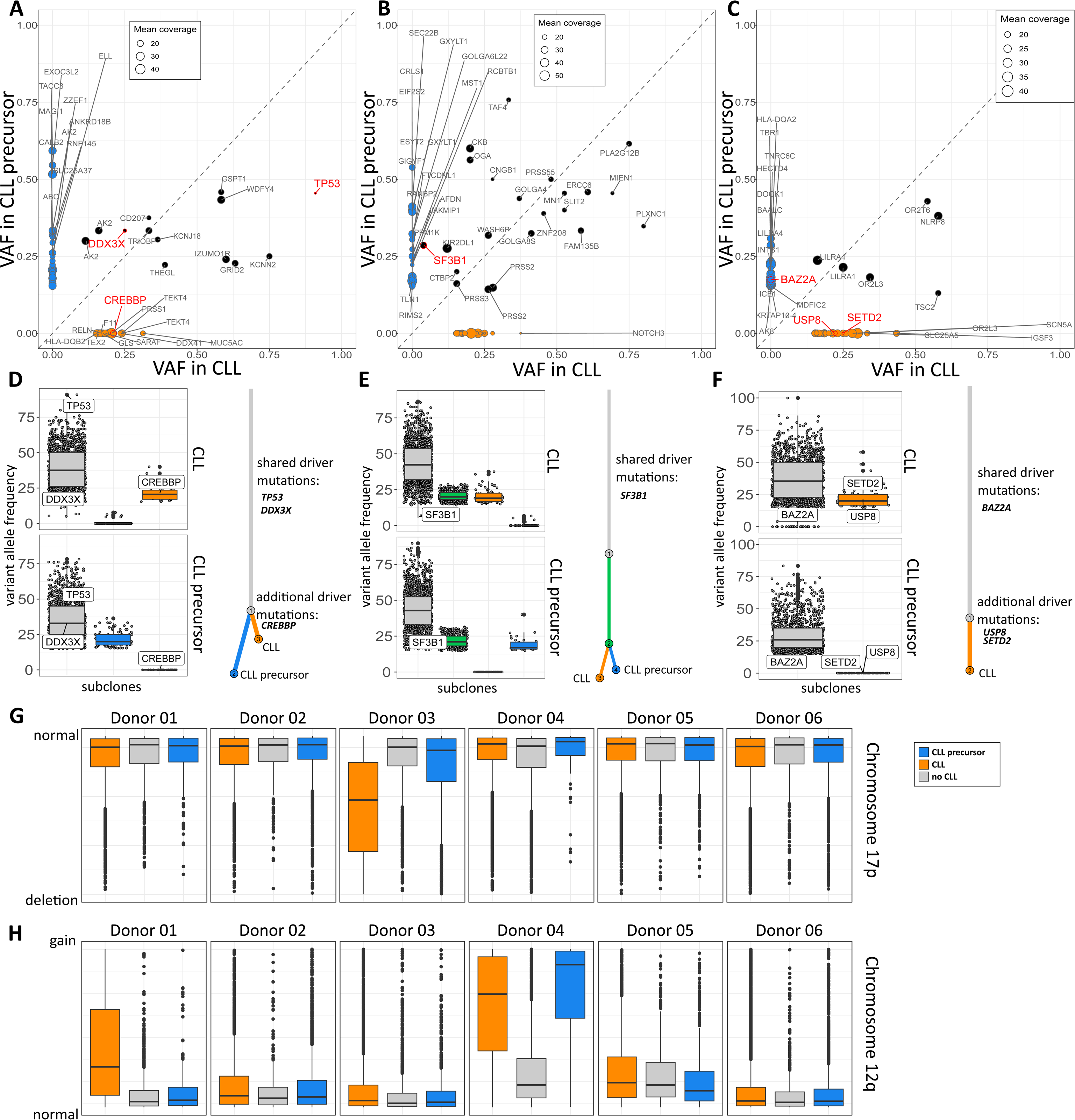
Whole genome variant analysis and CNV analysis of CLL subpopulations. **A-C)** Variant allele frequencies (VAF) of mutated genes in either CLL (orange) or CLL precursor cells (blue), or both (black) from three CLL patients. Dot sizes reflect the mean coverage. CLL driver mutations are marked in red. **D-F)** Sub clones (left side) and deduced clonal evolution trees (right side) with shared and distinct driver mutations according to the analysis with the R-packages sciClone and ClonEvol from three patients. **G-H)** Estimated copy numbers of selected gene loci located on chromosome 17p **(G)** and chromosome 12q **(H)** among all cells from each subpopulation, from scRNAseq data of six CLL patients, as determined by the R-package CONICS.

Although clonally related, both populations accumulated additional distinct sets of genomic variants. We used sciClone and ClonEvol^33,34^ to infer clonal hierarchies and reconstructed the clonal evolution tree of both subpopulations for each donor. Known CLL driver events^32^ were then mapped to the corresponding section of the tree (Figure 2D-F). This analysis suggested that after the shared accumulation of mutations, CD20^high^/CD5^low^ and CD20^low^/CD5^high^ CLL cells diverged in their mutational landscape, either in both subpopulations (Figure 2D and E), or exclusively in the main CLL population alone (Figure 2F).

To infer copy number variants (CNVs), we quantified expression levels of genes which would be affected by CNVs recurrently found in CLL in the six scRNAseq profiles using the software CONICS^35^. This analysis was separately performed for non-malignant B cells, CD20^high^/CD5^low^ CLL-related and CD20^low^/CD5^high^ CLL cells. We identified a trisomy 12 in both malignant populations but not in the polyclonal non-malignant B-cells in case CLL 04 (Figure 2G). In another example (CLL 01) we identified a trisomy 12 in the CLL main population but not in CD20^high^/CD5^low^ CLL-related cells, supporting their development at a timepoint before the occurrence of trisomy 12.

Taken together we show that CD20^high^/CD5^low^ CLL-related cells were genetically related to the CLL tumor, but also harbored distinct genomic features.

### BCR gene diversity and evolution in CLL precursor cells and the main CLL population

IgV-mutations in BCR genes are characteristic of memory B cells and M-CLL cells ^36–38^, and the presence and extent of IgV-mutations are predictive of clinical outcomes in CLL.^16,39^ CLL cells can actively diversify their BCR genes through somatic hypermutation, but this process is limited in MBL^26,27^ and CLL^40–46^, in contrast to the extensive diversity observed in normal memory B cell clones from healthy adults.^15,37,47,48^

We employed scVDJseq sequencing to delineate the intratumoral BCR diversity in all identified subpopulations, uncovering numerous subclonal variants of the CLL-BCR rearrangement, each distinguished by a range of one to fifteen IgV-mutations. Such IgV-mutations were observed in both CD20^high^/CD5^low^ CLL-related cells and the CD20^low^/CD5^high^ CLL main population. However, detailed IgV-mutation phylogenetics of the scVDJseq data revealed that BCR sequences from CD20^high^/CD5^low^ CLL-related cells showed significantly greater BCR diversity than the main CLL population, which always represented a skewed subset of this BCR pool (Figure 3A and S3A).

**Figure 3:**
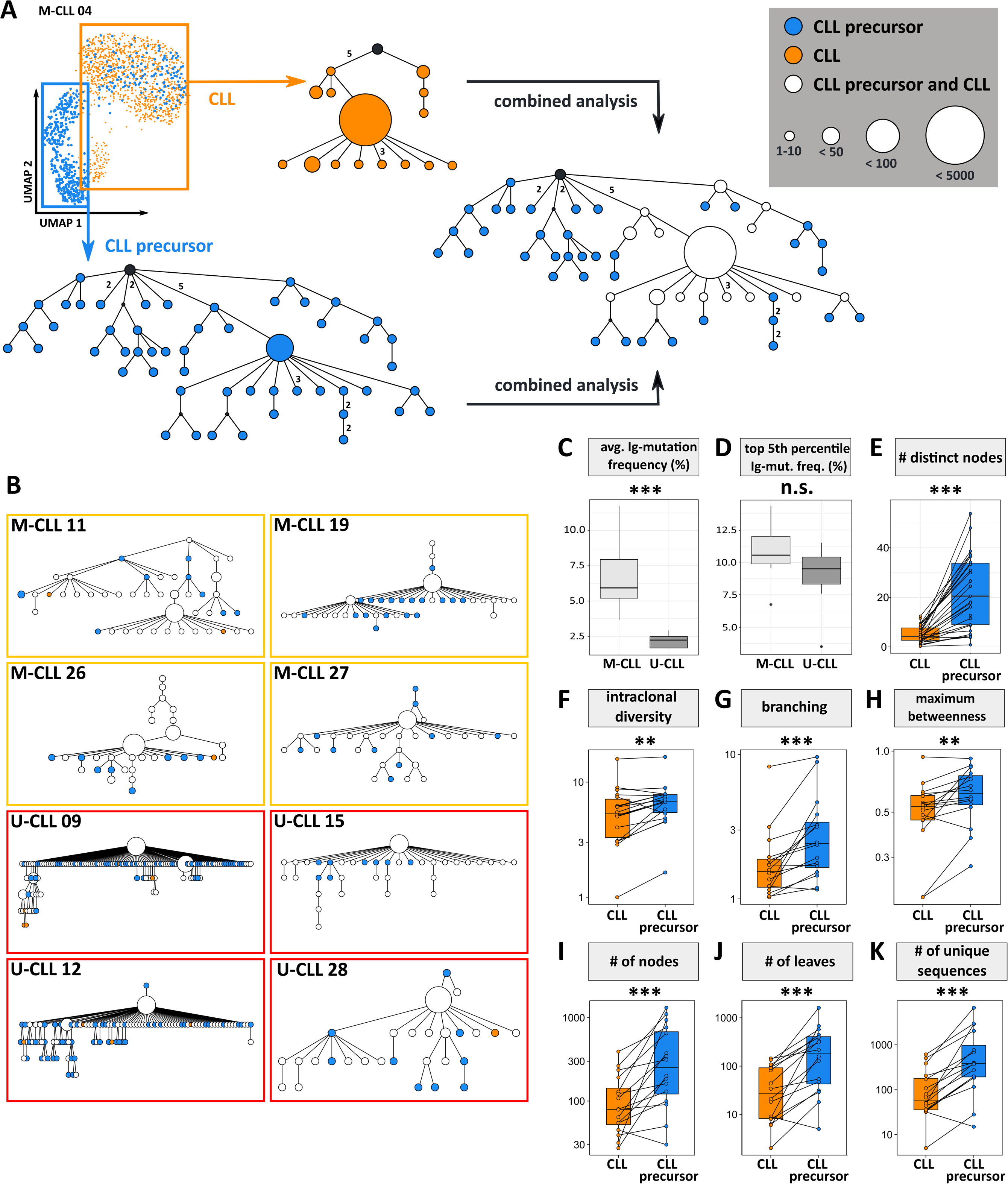
Phylogenetic analysis of IGHV-mutation patterns among CLL subsets. **A)** IgV-mutation dendrograms of CLL cells alone (orange), CLL precursor cells alone (blue), or in combination (nodes with CLL and CLL precursor cell sequences shown in white), based on scVDJseq data. UMAP and clustering of cells is shown to visualize subpopulation gating. **B)** IgV-mutation dendrograms of four M-CLL and four U-CLL independent cases analyzed by bulk VDJ-deep-sequencing. **C)** Median IGHV mutation frequency among M-CLL and U-CLL cases (n=21). **D)** IGHV mutation frequency of the top 5th percentile of mutated IGHV sequences in M-CLL and U-CLL (= the highest IGHV gene mutation frequency value of the 5% IGHV sequences with highest mutation load from each clone, n=21). **E-K)** IGHV mutation dendrogram parameters from 21 CLL cases analyzed with bulk VDJ-sequencing and replicate controls. ***p < 0.001, n.s. not significant, Wilcoxon test.

In a subsequent step, we aimed to validate our findings by examining IGHV-mutation diversity between bulk sorted CD20^high^/CD5^low^ CLL-related cells and the CD20^low^/CD5^high^ CLL main population (Figure 3B and S3B), across a cohort of 25 CLL cases (16 M-CLL and 9 U-CLL, as detailed in Table 2). This was conducted through two separate IGHV deep-sequencing approaches, employing replicate analysis to verify subclonal expansions (Figures 3B and S3B), and utilizing unique molecular identifiers (UMIs) to eliminate sequencing artifacts (Figure S3C-F). In all 25 cases analyzed, the IGHV-mutation patterns of CLL cells were consistently derived from those observed in the CD20^high^/CD5^low^ CLL-related cells, again supporting their role as a highly diversified pool of progenitor cells, from which the main CLL population inherited a selection of IgV mutation variants (Figures 3A, B and S3A-F).

As anticipated, M-CLL cases demonstrated greater BCR diversity, yet the maturation from CD20^high^/CD5^low^ precursors to CD20^low^/CD5^high^ CLL cells could be observed regardless of IGHV mutation status (Figures 3A, B and S3A-F). Even U-CLL cases predominantly exhibiting 100% germline identity in IGHV sequences we found small subset of CD20^low^/CD5^high^ cells with IgV mutation diversity inherited from their CD20^high^/CD5^low^ precursor pool which underscores their relatedness also in U-CLL. It is important to note, that this observation does not violate the concept of U-versus M-CLL, but it suggests that the U-versus M-CLL commitment has already been fixed at the level of CD20^high^/CD5^low^ precursors. CLL-IgV-mutation patterns were significantly enriched in AID-hotspot-motifs (p < 0.01, Figure S3G) and nonsense mutations were counter-selected (p < 0.001, Figure S3H). According to phylogenetic complexity parameters, IgV-mutation diversity is significantly enriched in CD20^high^/CD5^low^ CLL precursors versus main CLL population (Figure 3G-M), although the former subpopulation abundance in PB is minimal.

According to the WGS data (Figure 2) and the Ig-mutation hierarchies, the small subset of CD20^high^/CD5^low^ CLL-related cells develops early in pathogenesis and exhibits highly diversified Ig-genes, akin to the multiple cycles of mutation and selection known from normal memory B cells.^48,49^ This Ig-diversity seen in CD20^high^/CD5^low^ CLL-related cells is always found partly and skewed in the main CLL population. We deduce that both M-CLL and U-CLL cases originate from such CD20^high^/CD5^low^ CLL precursor cell pools, which is why we refer to CD20^high^/CD5^low^ CLL-related subpopulation from here on as “CLL precursor cells”.

### Longitudinal analysis of the evolution of CLL precursor cells

To determine whether CLL precursor cells represent a homeostatic population or exhibit growth dynamics, we measured the abundance and IgV-mutation patterns in CLL precursor cells versus the main CLL cell population in longitudinal samples from seven patients with variable clinical courses, spanning 5 to 16 years (average 10 years).

For indolent cases, characterized by a stable PB lymphocyte count and absence of treatment indication, the frequencies of CLL precursor cells remained stable across different timepoints. Correspondingly, the IgV-mutation patterns exhibited only minor structural alterations as shown in Figure 4A. Given that the sampling timepoints during the indolent course can span up to 10 years, this suggests a homeostatic expansion of CLL precursor cells.

**Figure 4:**
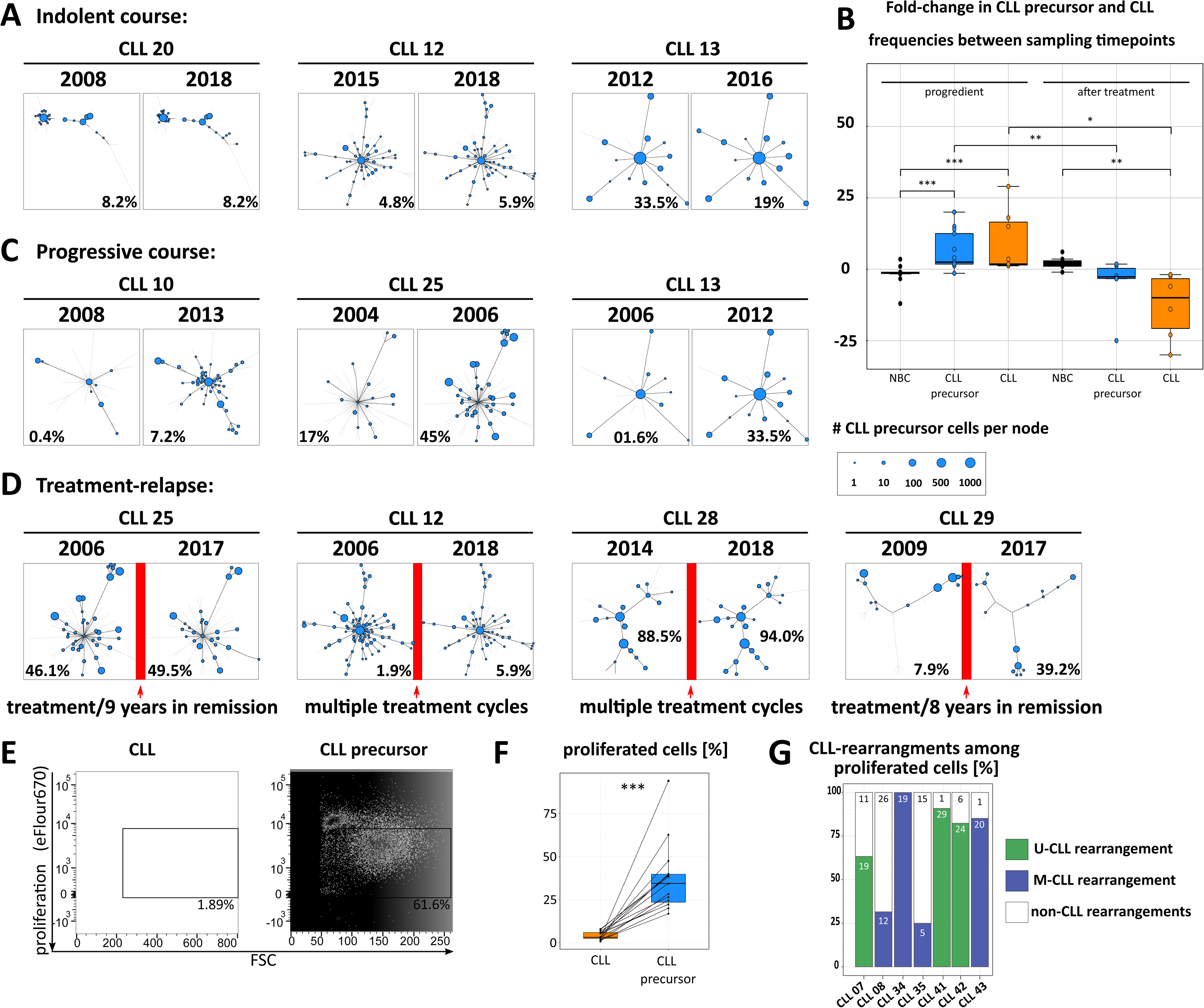
Longitudinal analysis of IGHV-mutation phylogenetics and proliferation potential among CLL and CLL precursor cells. A, C, D) Representative dendrogram structures from seven CLL patients with (A) indolent, (C) progressive course and (D) during treatment-relapse. Numbers give the percentage of CLL precursor cells among non-CLL B cells. B) Fold-change of CLL and CLL precursor cell frequencies between sampling timepoints in progredient cases or after treatment. E) Exemplary FACS-plot of proliferating CLL or NRB cells from stimulated samples. Proliferated cells are identified by dilution of the cell tracking dye eFlour670 and isolated by cell-sorting (black gates). F) Percentage of proliferated cells among the main CLL population and NRB cells (p < 0.01, Wilcoxon test, n=15). G) Percentage of the CLL-IGHV-rearrangement (U-CLL: green, M-CLL: blue) and non-CLL-rearrangements (white) among the total number of proliferated cells. Numbers give the amount of CLL- or non-CLL-IGHV-rearrangements identified per sample.

In patients showing progression, characterized by a rise in lymphocyte counts signaling the need for treatment, and in patients experiencing relapse, marked by an increase in lymphocyte counts post-treatment, there was a corresponding increase in the frequencies of CLL precursor cells along with CLL cells. Conversely, the count of non-malignant B cells decreased during these stages (as illustrated in Figure 4B). Crucially, in cases of CLL relapse, the IgV-mutation dendrograms of CLL precursor cells showed conservative reconstitution, with the arrangement and sequence of branches/nodes closely mirroring those from earlier timepoints. This pattern persisted even after extended periods of deep remission (as depicted in Figure 4D). These observations strongly suggested, that CD20^high^CD5^low^ CLL precursor cells show self-replenishing potential. Rarely, novel or previously undetected IgV-mutation variants emerged (Figure 4D). Across all tested timepoints, the dendrograms derived from CLL consistently included a biased assortment of IgV-mutation variants from the CD20^high^CD5^low^ CLL precursor cells (Figure S4).

To measure the expanding potential of CLL precursor cells, we performed *in vitro* proliferation assays. Up to 10^5^ NRB cells (CD20^high^CD5^low^) and CLL cells (CD20^low^CD5^high^) were isolated, marked with cell tracking dye and stimulated with CpG and CD40L *in vitro*. After 5 days the proliferated cells were isolated by cell sorting and the fraction of proliferating tumor cells was determined by IGHV gene PCR, cloning and sequencing, using the CLL-IGHV-rearrangement as tumor marker. Unstimulated normal B cells or CLL cells did not proliferate, suggesting that neither population included actively dividing cells in steady state. Upon stimulation, CLL cells mostly died within five days (Figure 4E). In contrast, stimulation induced proliferation among NRB cells of 15/15 CLL patients (Figure 4F). From 7 patients sufficient proliferating cells were recovered, and CLL IGHV rearrangements (55/147 total, range 5 to 29 per patient) were detectable among the polyclonal IGHV rearrangements of non-CLL origin, showing that the CLL precursor cells were responsive to activation (Figure 4G).

In summary, CLL precursor cells exhibit a wide range of Ig-diversity and represent a progenitor stage of CLL cells. Remarkably, even after achieving a substantial and prolonged remission, the pool of relapsing CLL precursor cells can restore their initial diversity as observed before treatment. This may suggest that these cells have the capacity for self-renewal, enabling them to persist over time.

### CLL cells develop from CLL precursor cells throughout the course of the disease

Here we have shown that CLL cells originate from the CD20^high^/CD5^low^ precursor population. To investigate whether CLL precursor cells can seed subclonal development in progressive disease, we determined their temporal and hierarchical relationship by IgV-phylogenetics on longitudinal samples from seven cases. To allow statistical comparability, equal sequence numbers were randomly drawn from each timepoint, pooled and analyzed. The resulting dendrogram was used as a scaffold, on which the individual sequence patterns per timepoint were displayed (Figure 5A).

**Figure 5:**
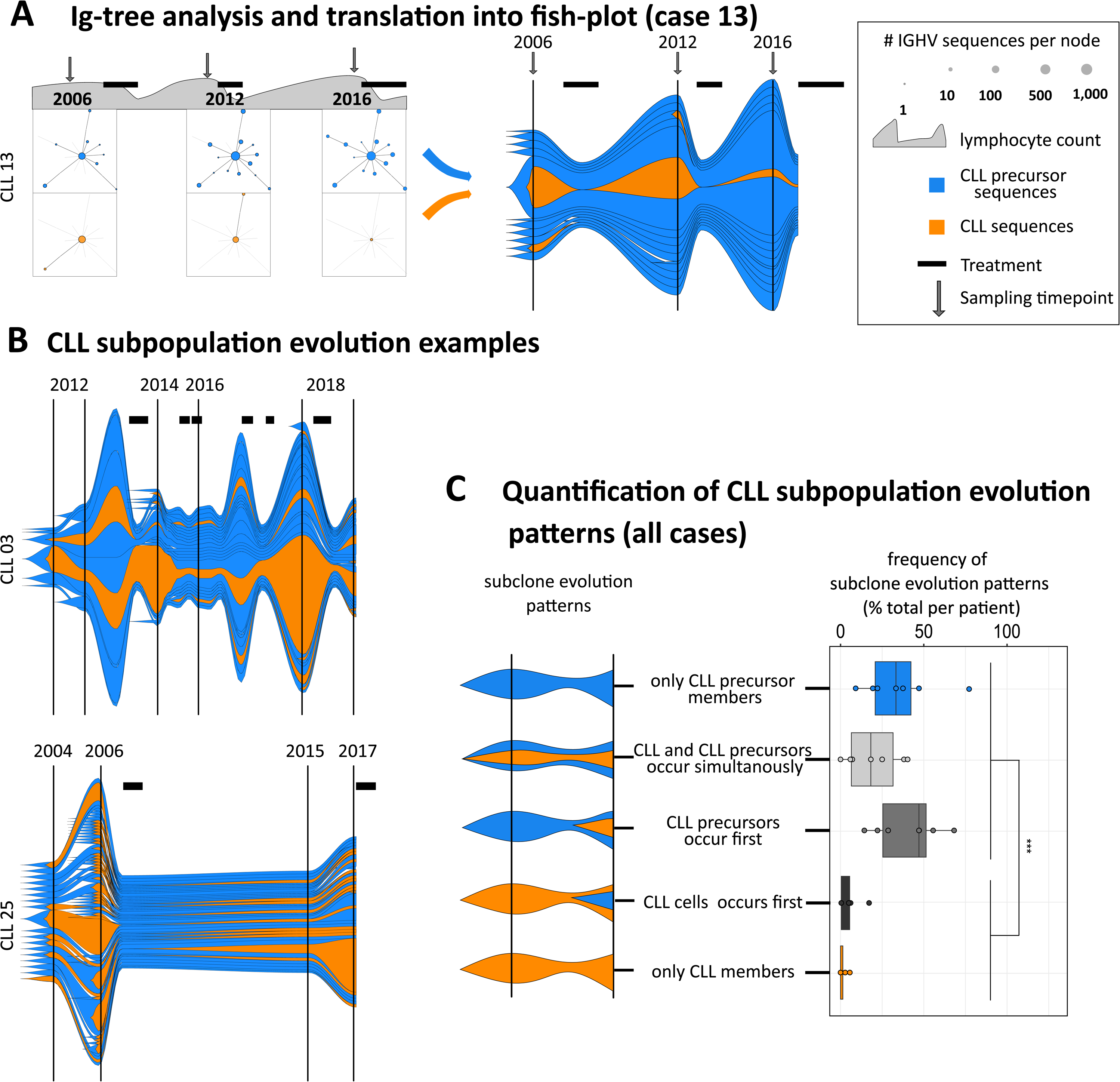
Longitudinal analysis of CLL subpopulation differentiation and evolution. IgV-mutation dendrograms of CLL precursor-(blue) and CLL-derived IGHV sequences (orange) from longitudinal samples were translated into fish plots to depict subclonal development. A) representative spread-dendrograms of CLL 13, presented as fish-plot diagram (B), normalized to the lymphocyte count and the number of IGHV sequences per node. Treatment and sampling timepoints are indicated by bars/arrows. C) Examples of clonal evolution from four individual CLL patients. D) shows the relative frequency of the five subpopulation differentiation categories, that are detectable in the fish plot analysis of each patient. **p < 0.01, Wilcoxon test.

To visualize development of each subclone (node) over time, we generated fish-plots which show the number of unique Ig-sequences per node and per timepoint, normalized to the PB lymphocyte count (Figure 5B). Fish-plots provide a visual representation of the size and individual trajectory of distinct precursor or mature CLL populations with similar immunoglobulins. These plots track changes over time, capturing stages of progression, treatment, and relapse (Figure 5B). If our hypothesis is correct, CLL precursors would give rise to CLL cells and might further evolve independently without maturing into CLL cells. We quantified the following hypothetical constellations to investigate if alternative scenarios not supporting our hypothesis were also possible (Figure 5B): a) CLL precursors may evolve independently, b) CLL precursors and CLL evolve in parallel but the maturation process from CLL precursors into CLL has not been captured by our time course c) CLL precursors have matured into CLL and evolve continuously, d) CLL may precede its related precursors e) CLL cells evolve independently of CLL precursors. Indeed, scenarios d) and e) were almost undetectable, reinforcing the hypothesis that mature CLL cells originate from CD20^high^/CD5^low^ CLL precursors. The remaining scenarios occur with nearly equal frequency.

### CLL precursor cells represent dedifferentiated memory B cells with leukemic stem cell signature

The scRNAseq of six patients identified 4,884 transcripts consistently expressed in all samples, of which 227 (4.64 %) were expressed at log2-fold-change `≥` 0.5 (Figure 6A and Table S1) between CLL precursor and CLL cells. In line with their elevated IgV-mutation diversity, CLL precursor cells showed elevated expression of memory B cell genes (Figure 6A and Table S1), and singleR projection of published transcriptome signatures^15,50^ showed that 25-55% of CLL precursor cells were more similar to (CD5^+^) memory B cells than to CLL cells (Figure 6B), this was independent of the M-CLL or U-CLL origin.

**Figure 6:**
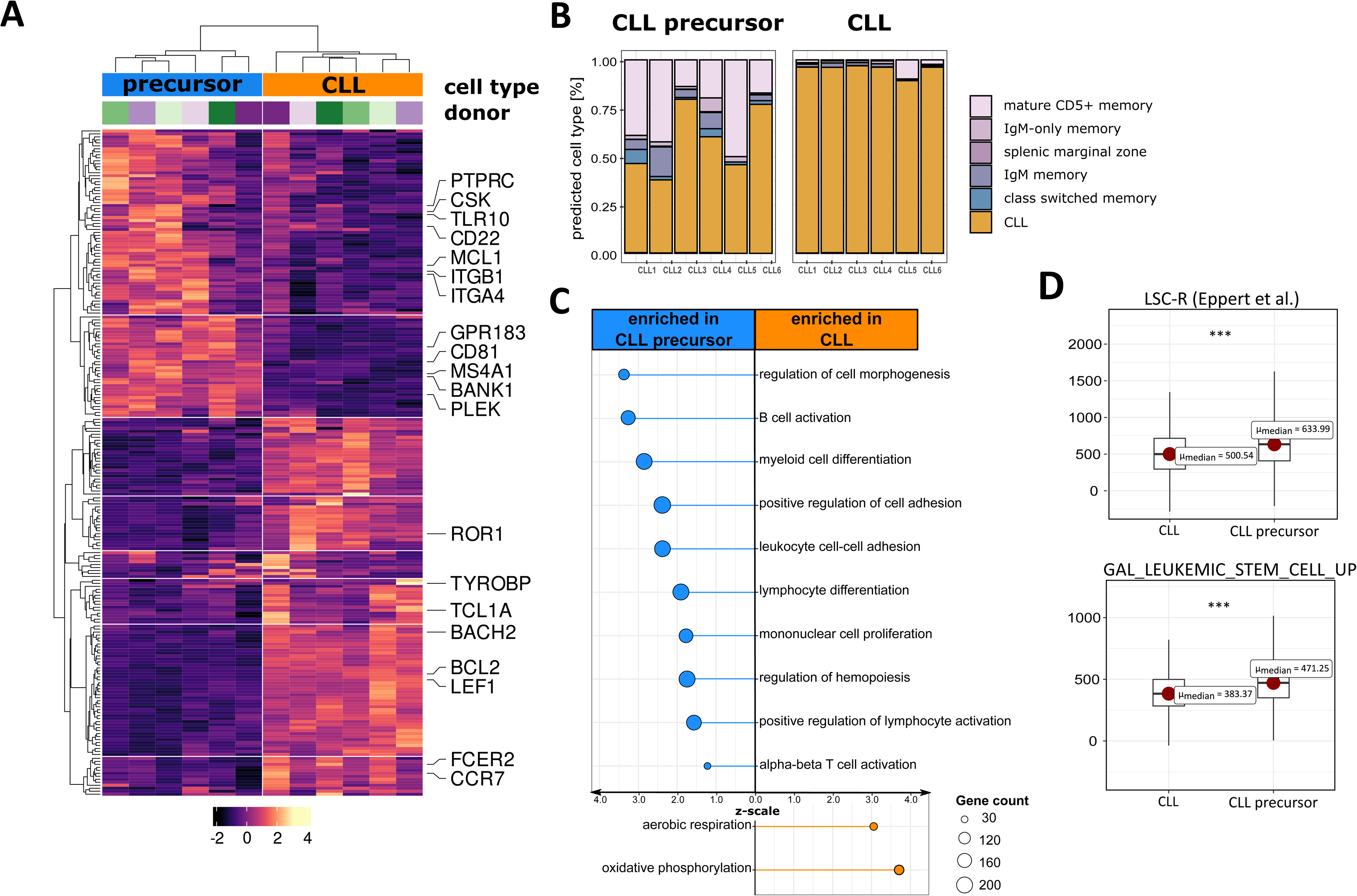
Transcriptomic characterization of CLL subpopulations. **A)** Heatmap of the statistically significant differentially expressed genes between CLL and CLL precursor cells (n=1,309), repeatedly identified in all six cases analyzed. **B)** Single-cell RNAseq annotation (SingleR) of CLL and paired CLL precursor cells, according to their global similarity to normal B cell signatures or to U-CLL/M-CLL signatures.^15^ For each cell only the predominantly assigned cell type is given. **C)** Gene Ontology pathway analysis of the genes enriched in CLL precursor versus CLL cells (see also Table S2). **D)** Enrichment plots of gene sets associated with human leukemic stem cells between CLL precursor versus CLL cells. ***p < 0.001, Wilcoxon test.

According to unsupervised pattern recognition algorithms, CLL precursor cells are enriched for signatures related to B cell activation, proliferation, adhesion, and cell differentiation programs (Figure 6C and Table S2), supporting our functional and immunogenetic data (Figures 4 and 5). Moreover, CLL precursor cells are significantly enriched for genes regulating hemopoiesis (Figure 6C) and gene sets curated from leukemic (but not solid) cancer stem cells (Figure 6D). The analysis of hallmark hematopoietic stem cell genes showed that CLL precursor cells do not share a typical CD34^+^CD38^−^ phenotype, nor express CD133 or CD117 (KIT).

We conclude, CLL precursor cells do not represent transformed stem cells, as seen in other leukemias, but show a mixed phenotype of conventional CLL cells and CD5^+^ memory B cells, and their transcriptional patterns suggest that a fraction of CLL precursor cells may have regained leukemic stem-cell-related signatures during transformation.

## Discussion

Through scRNAseq/scVDJseq sequencing, we identified a novel CLL precursor cell subpopulation characterized by high levels of CD20 and low CD5 expression. These cells share heavy and light chain IgV-rearrangements with their corresponding CD20^low^/CD5^high^ CLL tumor cells and simultaneously express normal memory B cell signatures as well as recurrent CLL transcription programs.

As normal residual B cells are underrepresented in CLL patient blood, and the CLL precursor cells often represent only a small fraction among the CD20^high^ normal B cell pool, the CLL precursor cells had to be enriched by cell sorting to be above detection limit, which is likely the reason they have not been detected in previous single-cell studies.^7,18,51–53^

A remarkable distinctness of CLL precursor cells is their genomic diversity. The genome-wide somatic variant landscape of the CLL precursor versus CLL population suggests that the former experienced extensive somatic diversification, of which on average only 13% of the somatic variants are shared by the CLL tumor majority. This, and the VAF analysis of genomic variants supports the clonal relation of CLL precursor and CLL cells, but also indicates developmental distinctness between both subpopulations. An additional characteristic of CLL precursor cells is their high intraclonal IgV-mutation diversity, which is present in both M-CLL and U-CLL subsets in our cohort of 70 CLL cases.

To deeply evaluate the developmental dynamics and temporal relationship of CLL precursor and CLL cells, we made use of IgV-mutation phylogenetic measures as molecular clock and developmental framework.^37,47,48^ IgV-mutation phylogenetics of 70 cases showed that the very rare CLL precursor cells represent a highly diversified pool of tumor cells that represent a “framework” from which the CLL tumor mass always represents a skewed sub-selection. In M-CLL cases, the CLL mass developed from IgV-mutation variants more downstream in the Ig-mutation-tree, in U-CLL cases from variants more upstream, but not necessarily the “germline”-node. Importantly, when calculating the average IgV-mutation frequency of all detected sequences, the CLL cells of U-CLL cases always showed an average IgV-mutation frequency below 2%, and M-CLL cases above 2%, hence, although the CLL precursor population is always highly Ig-diversified (> 2% average IgV-mutation frequency), the 2% cut-off rule is not violated in our cohort.^16^

The pathogenetically early generation and persistence of CLL precursor cells is supported by chromosomal CNVs, which are early, driving events in CLL pathogenesis.^13,14^ The prediction of chromosomal CNVs in our scRNAseq/scVDJseq data sets confirmed the common presence of a known, monoclonal 17p deletion in one patient, and absence of a known trisomy 12 only in CLL precursor cells of another patient, the latter of which strongly supports their generation before acquisition of trisomy 12. The idea that subclonal diversity is created early in CLL pathogenesis is also supported by epimutation studies^18^ and subclonal variants in the CLL precursor MBL.^26^ CLL precursor cells differ from the asymptomatic precursor state MBL by their lower frequency in PB, lack of the typical CLL-like immunophenotype and higher IgV-diversity.^26,54,55^

There is more to learn from IgV-mutation phylogenetics in CLL, in particular when analyzing longitudinal samples. First, during indolent course, the intraclonal diversity and subclonal composition of both, CLL precursor and CLL cells is stable. Only few “novel” IgV-mutation variants can be detected. It is unclear whether these derive from ongoing diversification, or just from sampling variation. During progressive disease, CLL precursor- and CLL cell numbers and diversity increase in the same order of magnitude, however, CLL cells remain a sub-selection of the CLL precursor diversity framework. Importantly, treatment affects both subpopulations, albeit CLL precursor cells may be tendentially less vulnerable. It is impossible to determine at this stage, whether the type of treatment impacts on this tendency, which would require drug screening approaches and substantially larger cohorts. Importantly, upon relapse – even after long periods of deep remission – CLL precursor cells recover conservatively, indicating self-replenishment, a hallmark of tumor-originating cells. In contrast, the main CLL tumor subclonal composition changes, with the dominant subclone locating often to a previously underrepresented node. The occurrence of novel Ig-variants is rare, and the majority of the IgV-variants that developed at one timepoint (often covered by thousands of sequences among the main CLL population), are already present years earlier among a small number of CLL precursor cells, but not vice versa. Hence, CLL precursor variants can develop into CLL, but there is barely indication that CLL subclones give rise to CLL precursor cells. Moreover, there is no indication that CLL variants can occur independently of CLL precursor cells, whereas there are multiple examples for independent CLL precursor subclones. Together, this suggests that the second hallmark of tumor-originating cells, i.e., the differentiation capacity, is confined to the CLL precursor population.

*In vitro* stimulation experiments further support that CLL precursor cells show proliferative capacity, and deviate from the mostly unresponsive, and apoptosis-prone behavior of bulk MBL/CLL cells, unless these are cultured with feeder cells.^56,57^ In addition, CLL precursor cells are enriched for leukemic stem cell expression signatures. We assume, CLL precursor cells develop from mature memory B cells which partly dedifferentiate during transformation. This is supported by their transcriptional and phenotypical similarity to normal memory B cells (which is more prominent than in the main CLL population), and their high IgV-mutation content, even in U-CLL cases.

Our study identified a previously unrecognized tumor subpopulation in M-CLL and U-CLL cases, which develops early in pathogenesis, is characterized by high intraclonal Ig-diversity, and persists. According to longitudinal Ig-dynamics, these “a-priori” diversified subclones show central features of lymphoma stem cells, i.e., proliferation capacity, self-renewing potential, and finally, differentiation into “newly” evolved subclones in the shape of conventional CLL cells. The existence of lymphoma stem cells is unclarified in non-Hodgkin lymphomas,^4–6^ but may be of decisive interest to predict tumor progress or resistance development. Especially in CLL, lymphoma-originating cells are supposed to locate to proliferation centers in the tissue, but we assume that these have a circulating counterpart, that disseminated and confer CLL spreading to previously uninvolved lymphoid tissues during the course of the disease.

## Materials and methods

### Sample preparation

B cells were isolated by Ficoll density centrifugation (density 1.077 g/ml, Pan BioTech, Aidenbach, Germany) followed by staining with anti-CD3 (BD Biosciences, Heidelberg, Germany), anti-CD5 (BioLegend, Koblenz, Germany), anti-CD20, anti-CD23, and anti-CD27 (each BD Biosciences) antibodies. Stained cells were analyzed on a CytoFLEX S flow cytometer (Beckman Coulter, Krefeld, Germany) using CytExpert v2.4 (Beckman Coulter) or FlowJo v10.6.2 (BD Biosciences) software. Sort-purification was performed with a FACSAria Fusion cell sorter (BD Biosciences).

### Single-cell RNA- and IGHV-sequencing

Combined scRNAseq and scVDJseq was performed with a Chromium Controller and Next GEM technology (10XGenomics, Pleasanton, CA, USA) using the “Chromium Single Cell V(D)J Enrichment Kit, Human B Cell” and “Chromium Next GEM Single Cell 5’ Library and Gel Bead Kit v1.1” (10XGenomics) for the recovery of up to 15,000 single cells. CLL precursor (CD20^high^/CD5^low^/ROR1^+^) and CLL cells (CD20^low^/CD5^high^/ROR1^+^) were sort-purified and separately loaded on the chip. The generated libraries were purified, and the VDJ libraries were paired end sequenced using the MiSeq v3 reagent kit (Illumina, San Diego, CA, USA), the GEX were paired end sequenced using the NextSeq2000 (100 cycles) v3. Reads were aligned with cellranger v3.1.0 (upon GRCh38 2020 version) and analyzed using Seurat^58^ with the following settings: all genes found at least in ten cells, all cells with at least 800 different genes, a count of at least 3000 reads per cell and cells with no more than 7 percent reads from mitochondrial RNA were further considered. Only cells for which an IGHV-rearrangement was obtained were processed as this was used to identify the cells from the CLL tumor clone. Identification of distinct cell phenotypes was carried out by SingleR^59^ using our own array-based datasets.^15,50^

### Pathway analysis

Pathway analysis among differentially expressed genes between subsets was performed via clusterProfiler and enrichR.^60^

### CNV detection

Copy numbers were detected using CONICSmat^35^ and compared with known copy number aberrations from FISH analyses.

### Whole genome sequencing, alignment, and variant calling

CLL, CLL precursor and T cells were sort-purified from three donors. Libraries were generated with the NEBNext Ultra II (NEB, Ipswich, Massachusetts, USA) kit and sequenced on a Nova Seq (Illumina, San Diego, California, USA) with standard settings and a desired coverage of 30 for each of the nine samples. The sequences were aligned with bwa mem (v. 0.7.17-r1188; standard settings)^61^ to GRCh38. PCR duplication was excluded with bammarkduplicates (v. 2.0.180;)^62^ and variant calling was perform using strelka (v. 2.9.10)^63^ with standard settings. Only sequences with *pass* were further considered.

### Whole genome subclonal architecture inference

The R-package sciClone^34^ was used to infer the subclonal architecture of each tumor patient. Variant allele frequency (VAF) plots and inferred trees were generated by ClonEvol,^33^ which uses the clonal information from sciClone together with the VAF to infer the clonal orderings and to reconstruct the clonal evolution trees. Driver events were mapped to the corresponding section of the tree.

### Library generation and BCR repertoire sequencing

UMI strategy (custom): We sort-purified CLL precursor (CD20^high^/CD5^low^/ROR1^+^) and CLL cells (CD20^low^/CD5^high^/ROR1^+^) from four CLL cases with known IGHV rearrangement in aliquots of 5,000 to 10,000 cells each and extracted genomic DNA (Gentra Puregene Core Kit, Qiagen, Hilden, Germany). Library generation was performed as described.^64^ In short, rearranged IGHV genes were amplified by PCR for the MiSeq Illumina platform (Illumina, San Diego, California, USA). In the first two PCR cycles 12 nucleotide UMIs are introduced and amplified in the second PCR. Sequences with identical UMI were classified as PCR duplicates and reduced to the longest detected consensus sequence. Each ambiguous nucleotide was replaced by the most frequent (majority vote) nucleotide.

Replicate strategy (InvivoScribe): Genomic DNA was extracted from sort-purified cells from 22 patients by QiaAmp DNA micro kit (Qiagen). For reliable identification of expanded clones, each B-cell population was analyzed in two replicates. IGHV gene rearrangements were amplified using the LymphoTrack IGH FR1 Assay kit (InvivoScribe, San Diego, CA, USA). Amplified DNA was processed for HTSeq according to the LymphoTrack protocol, amplified with a 1-step multiplex PCR for 29 cycles using subgroup-specific IGH framework region 1 primers and IGHJ primers, incorporating platform-specific adaptors. PCR products were purified and paired-end sequenced using the MiSeq v3 reagent kit (Illumina, San Diego, CA, USA). Reads with quality score of ≥ 30 and mean quality score ≥ 35 were further processed. Ambiguities between forward and reverse reads were denoted as ‘N’. Identical sequences were reduced to the longest detected sequence (collapsing) and each ‘N’ in a sequence was replaced by the most frequent (majority vote) nucleotide. Only sequences detected more than once were further processed. For full information regarding sequencing numbers, see Table S3.

### Determination of clonal sequences

Sequences were considered as clonally related when using the same IGHV gene and sharing at least 95% complementarity determining region III (CDRIII) nucleotide sequence homology, including up to 5% length variation. Identification of the IGHV gene was performed in R (www.R-project.org/) using rBLAST and the international ImMunoGeneTics information system (IMGT) database (www.imgt.org). Mutation dendrograms were calculated with IgTree.^65^

### Random drawing for phylogenetic dendrogram structure identification

For each patient, a fixed number of CLL IgV gene sequences, equal to the number of total CLL precursor IgV gene sequences from the same timepoint, was drawn randomly from the uncollapsed number of sequences. To remove putative sequence contaminations (e.g., by PCR artefacts or barcode hopping), a Gaussian Mixture Model was used and collapsed sequences that did not belong with 100% certainty to the CLL precursor cell distribution were excluded from the calculation. A Mixture Model is a mixture of k component distributions that collectively make a mixture distribution.

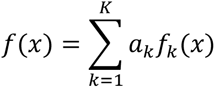

*α*_*k*_ represents a mixing weight for the *k*^*th*^ component where 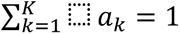. The R-package mixtools^66^ was used to calculate the probability with a log-likelihood approach.

### Phylogenetic analysis

All calculations (e.g., tree parameter etc.) were based on the whole un-simplified dendrograms including all available (drawn) sequences. Note that only for visualization in the figures, the full dendrograms were “simplified” by community calculations from the igraph R package,^67^ as otherwise the number of nodes and branches would be overwhelming and impossible to be displayed in a suitable resolution.

#### Maximum betweenness centrality

The Maximum betweenness centrality is calculated from the number of shortest paths through a node:

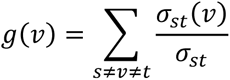

*v, s, t = nodes, σ_st_* =total number of shortest paths between *s* and *t*, *σ_st_*(*v*) = number of those paths which path through *v*. For each dendrogram the maximum number is given. This is a measurement of importance of nodes.

#### Branching

Branching calculates the mean number of outgoing edges (branches) per sample.

#### Intraclonal diversity

The intraclonal diversity is calculated from the approximate string distance between each sequence. The distance is a generalized Levenshtein (edit) distance, giving the minimal possibly weighted number of insertions, deletions and substitutions needed to transform one string into another. For each dendrogram, the maximum number is given. This is a measurement of the variability of the dendrogram.

### Clonal evolution analysis

To assess changes in CLL precursor abundance, the number of CLL versus non-tumor BCR-rearrangements retrieved from CD20^high^CD5^low^CD27^low^ B cells between sampling-timepoints was calculated and normalized to the total lymphocyte count; to assess changes in the overall tumor load, fold-changes of the total lymphocyte count were determined.

Dendrogram nodes representing Ig variants were translated into fishplots to illustrate longitudinal subclonal evolution of CLL precursor alongside main CLL cells. The plots were generated based on the R Fishplot Package (v0.5.2).^68^ In order to visualize rare Ig variants, an individual binning for each patient was applied.

### In vitro proliferation assay

NRB and CLL cells were sort purified and stained with 5 µM eFluor 670 cell proliferation dye (ThermoFischer, Oberhausen, Germany). eFluor670^high^ cells were cultured in RPMI 1640 medium supplemented with 20% fetal bovine serum, 100 U/ml penicillin and 100 μg/ml streptomycin (all from Pan Biotech, Aidenbach, Germany) at 37°C and 5% CO2. Stimulation was performed using 0.03 μg/μl anti-immunoglobulin (anti-Ig, Jackson ImmunoResearch, Newmarket, UK) and 1 μg/ml CD40-ligand-HA with 5 ng/ml anti-HA antibodies (both R&D Systems, Minneapolis, USA), and Type B CpG OND D-SL01 (InvivoGen, San Diego, USA). After 5 days of *in vitro* culture, proliferated (eFluor670^low^) and non-proliferated (eFluor670^high^) fractions were sort-purified and RNA extraction was performed (miRNeasy micro-Kit, Qiagen). After cDNA synthesis (High-capacity cDNA Reverse Transcription Kit, Applied Biosystems) two rounds of multiplex PCR using a mixture of custom IGHV- and IGHJ-primers^69^ with a total of 75 cycles were performed. PCR products were ligated into pGEM-T easy vectors (Promega, Walldorf, Germany) and transformed into JM109 E. coli (Promega, Walldorf, Germany). Bacteria were grown over night on LB/amp/IPTG/X-gal plates and non-colored colonies were selected for plasmid preparation. Plasmids were extracted by mi-Plasmid Miniprep Kit (Metabion, Planegg, Germany) and processed with Sanger sequencing. CLL tumor cells were identified by their respective IGHV-rearrangement.

### Statistics

Dendrogram parameters and proliferation capacities were analyzed with a Wilcoxon signed rank test, * = p < 0.05, ** = p < 0.01, *** = p < 0.001.

### Study approval

PB samples were obtained after written informed consent was received from participants prior to inclusion in the study according to the Declaration of Helsinki, and approval by the ethics committee of the Medical Faculty at the University of Duisburg-Essen, Germany (BO-10-4380).

## Author contribution

M.S. developed the concept. M.S., B.B., R.K., and J.D. designed and developed experimental strategies, contributed to data evaluation, and prepared the manuscript. B.B., M.B., A.K., K.B., M.D., E.H., and S.T. designed and carried out most of the experiments. B.B. and B.F. developed and performed bioinformatical and statistical analysis. P.J., J.B., J.v.T., H.C.R., S.S., and J.D. contributed samples, clinical data evaluation and helpful discussion.

## Acknowledgements

We thank Julia Jesdinsky-Elsenbruch for excellent technical assistance and Klaus Lennartz for his valuable engineering support. We thank Ludger Klein-Hitpass, Rene Scholtysik and the staff members of the Genomics and Transcriptomics Facility at the Institute of Cell Biology in Essen. This work was supported by the Deutsche Forschungsgemeinschaft (DFG) through SE1885/2-1, SE1885/2-2, SE1885/4-1, SFB1074 Subprojects B1 and B2, RE 2246/13-1, SFB1399-A01, SFB1430-A09, SFB1530-A01 and -C01, the DFG Clinician Scientist Program UMEA, FU 356/12-1, KU1315/16-1, the Deutsche Krebshilfe (1117240, 70115259, 70113041 and an Excellence Funding Program Grant to H.C.R.), the ERA-PerMed program HiRisk-HiGain (to H.C.R.), the intramural gender equity measure “Willkommen zurück”, the Dr. Werner Jackstädt-Stiftung, and the Deutsche Krebshilfe through grant 70112628.

## Notes

### Competing Interest Statement

The authors have declared no competing interest.

### Summary of Updates

author affiliations updated; Figures updated; supplemental files updated; additional analyses presented, section on CLL subpopulation identification updated to clarify relationship to the main tumor cells; longitudinal analyses presented in a way that self-replenishing and clonal evolution/hierarchy are now better visualized

