## Supplemental Figures 1-5, supplemental tables 1-3 for "A heterogeneous pool of tumor precursor cells with self-replenishing and tumor-originating properties in early pathogenesis of Chronic Lymphocytic Leukemia"

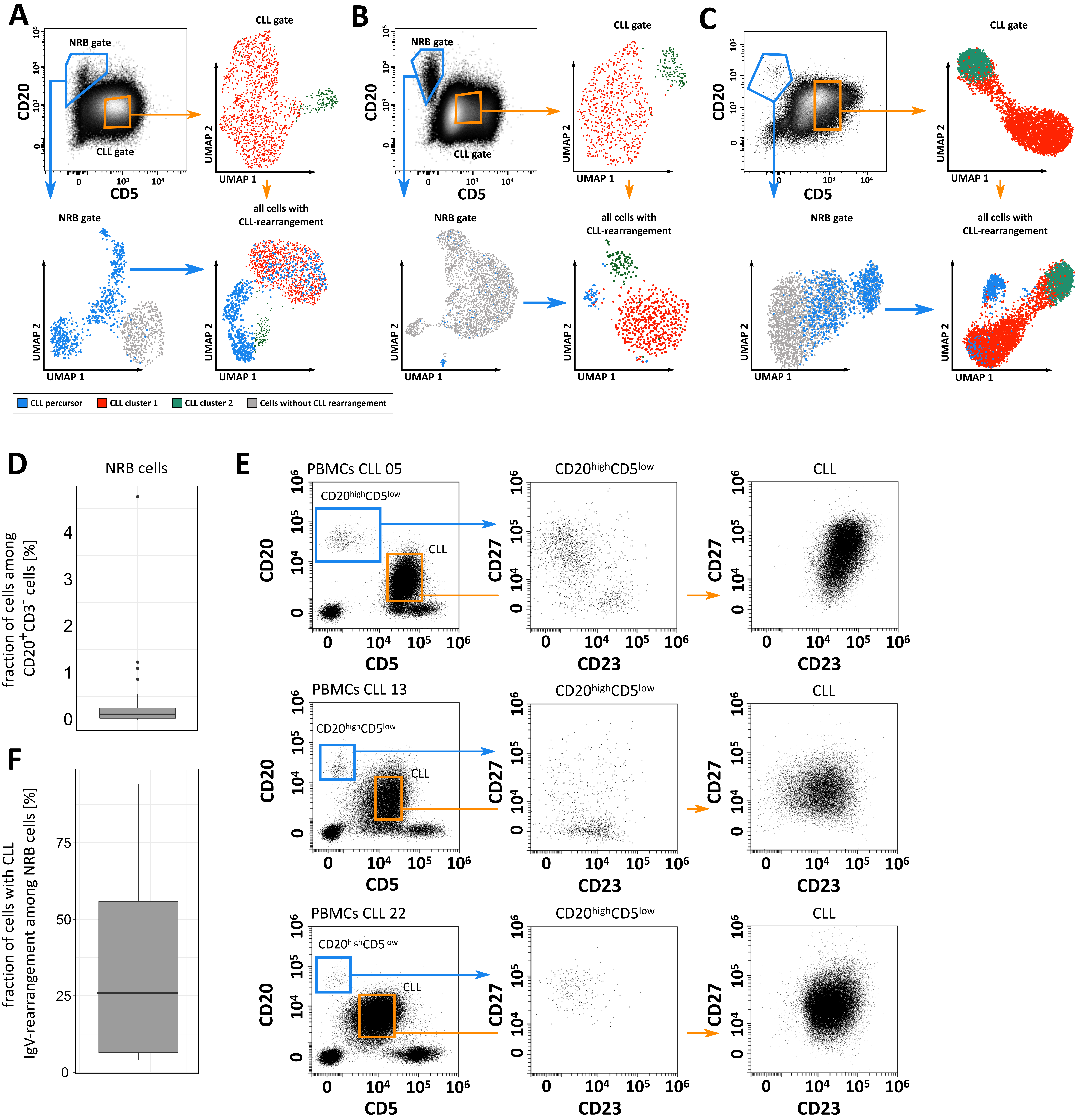


**Figure S1: Single-cell RNA/VDJseq analysis of circulating CLL cells**. A-C) The cell sorting strategy for CLL cells (orange) and normal B cells (blue) from three out of six cases (from left to right CLL 04 – 06) is shown in the top left plots. Top right plots: UMAP of scRNA profiles from the CLL gate (orange), showing two CLL clusters (red and green), all of which express the known CLL-IgH/IgL rearrangement. Lower left plots: UMAP of scRNA profiles from the normal B cell gate (blue), showing one cluster of normal B cells with polyclonal IgH/IgL rearrangements (grey) and one cluster of normal-associated CLL cells expressing the known CLL-IgH/IgL rearrangement (blue). Lower right plots: integrated UMAP and clustering from all cells expressing the known CLL-IgH/IgL rearrangement (red, green, and blue). D) Frequency of normal residual B cells among CD20+ cells in the PB of 60 CLL patients. E) Flow cytometric analysis of NRB cells (blue gate) and CLL cells (orange gate) among three selected CLL patients. The gates in the first column are representative for the cell sorting strategy. The cells were analyzed for expression of CD23 and CD27. PBMC: peripheral blood mononuclear cells. F) Frequency of cells with CLL-IGHV-rearrangement among CD20+ normal residual B (NRB) cells in the PB of 60 CLL patients.

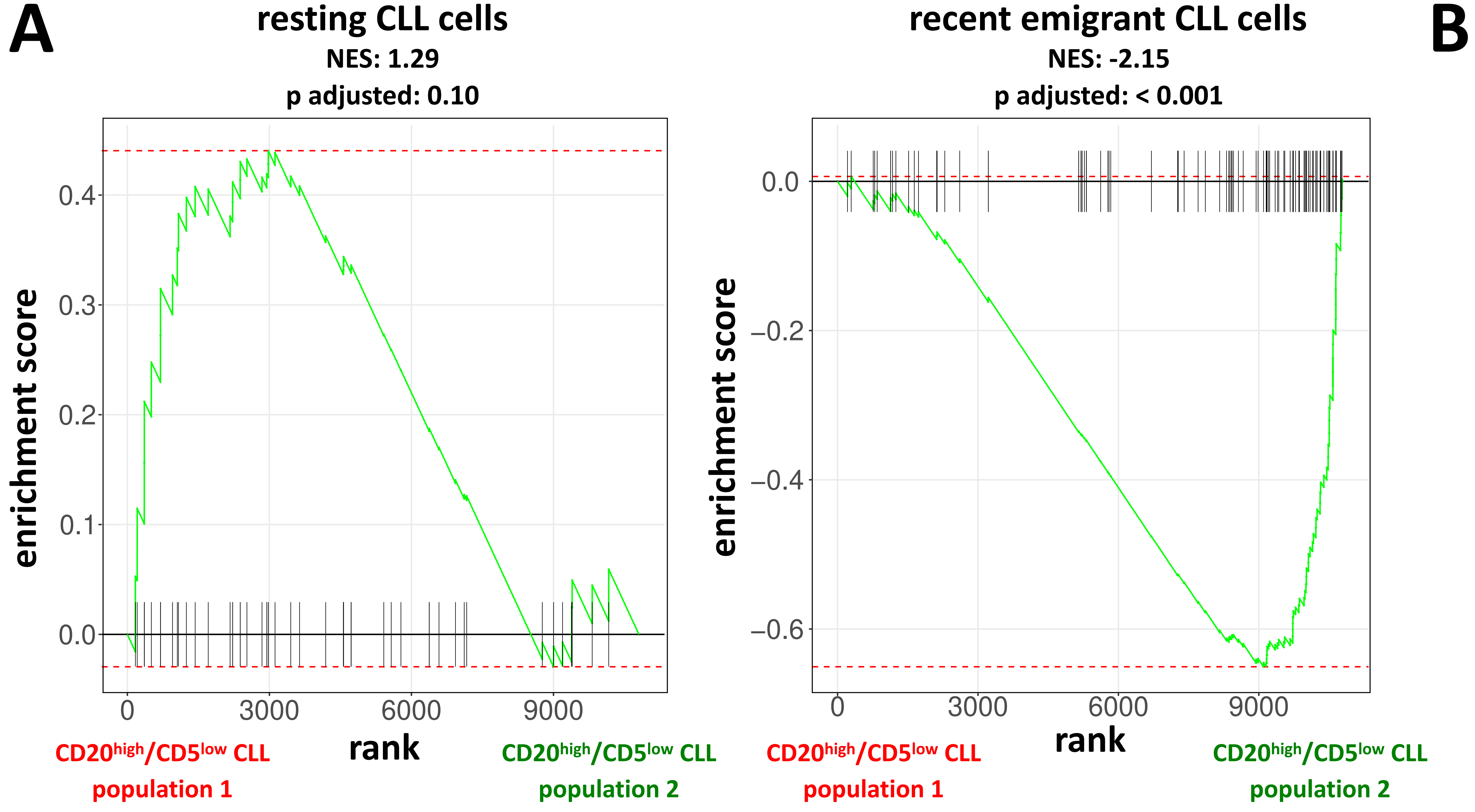


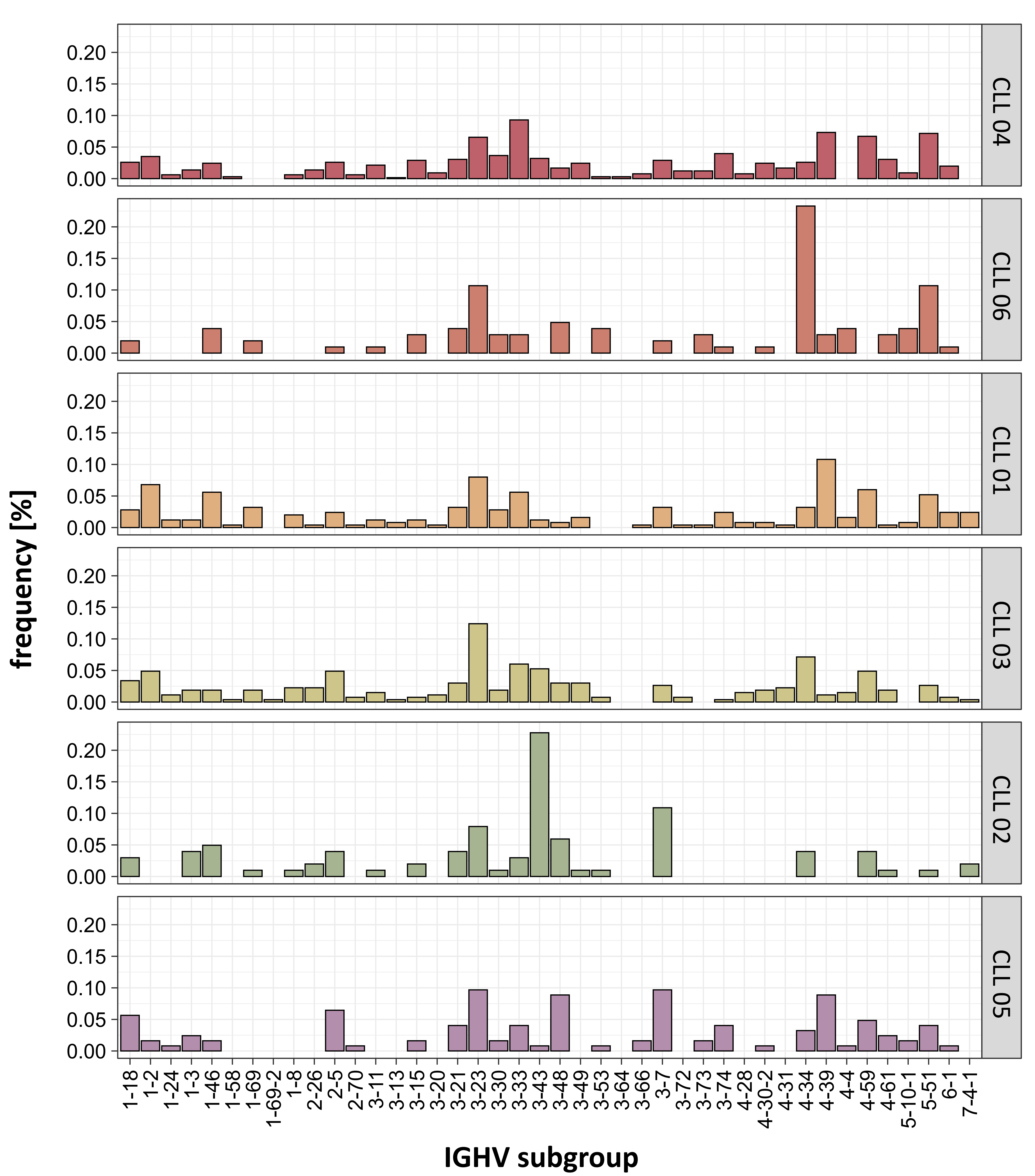


**Figure S2: scRNAseq profiling of CLL subsets.** (A) IGHV gene usage among normal residual B cells, i.e., lacking the CLL-IGHV-rearrangement in the PB of six CLL patients used for scRNA/VDJseq analysis. B) Gene set enrichment analysis of CLL precursor cells and CLL cells from six patients (CLL 01-06), based on the 159 genes differentially expressed in scRNAseq between resting and recently divided CLL cells.[24]

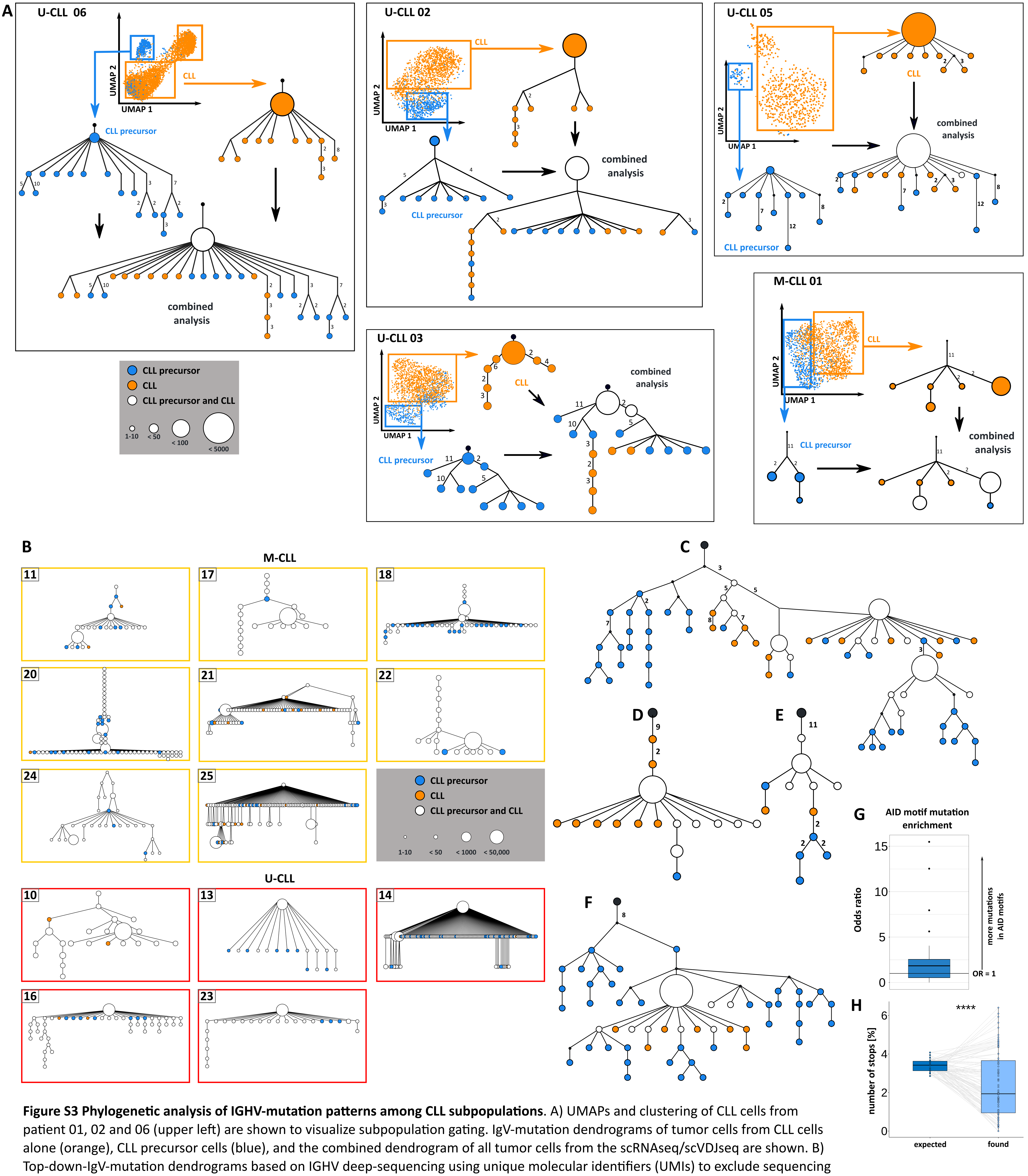


**Figure S3 Phylogenetic analysis of IGHV-mutation patterns among CLL subpopulations**. A) UMAPs and clustering of CLL cells from patient 01, 02 and 06 (upper left) are shown to visualize subpopulation gating. IgV-mutation dendrograms of tumor cells from CLL cells alone (orange), CLL precursor cells (blue), and the combined dendrogram of all tumor cells from the scRNAseq/scVDJseq are shown. B) Top-down-IgV-mutation dendrograms based on IGHV deep-sequencing using unique molecular identifiers (UMIs) to exclude sequencing artifacts. C-F) Top-down-IgV-mutation dendrograms based on IGHV deep-sequencing using replicate analysis to confirm subclonal expansions. G) CLL-IGHV-mutation patterns were significantly enriched in AID-hotspot-motifs ($p < 0.01$), and nonsense mutations H) were counter-selected ($p < 0.001$).

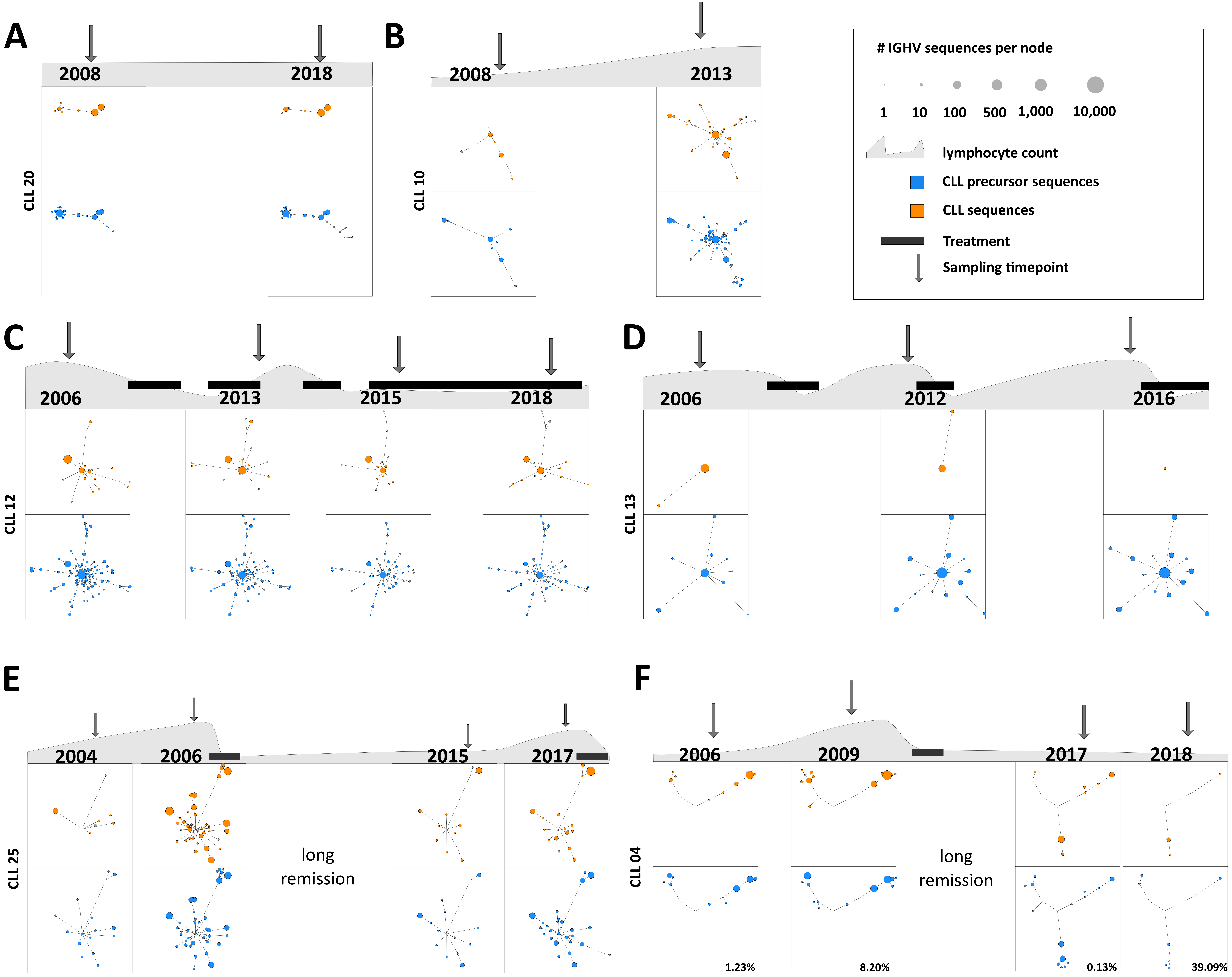


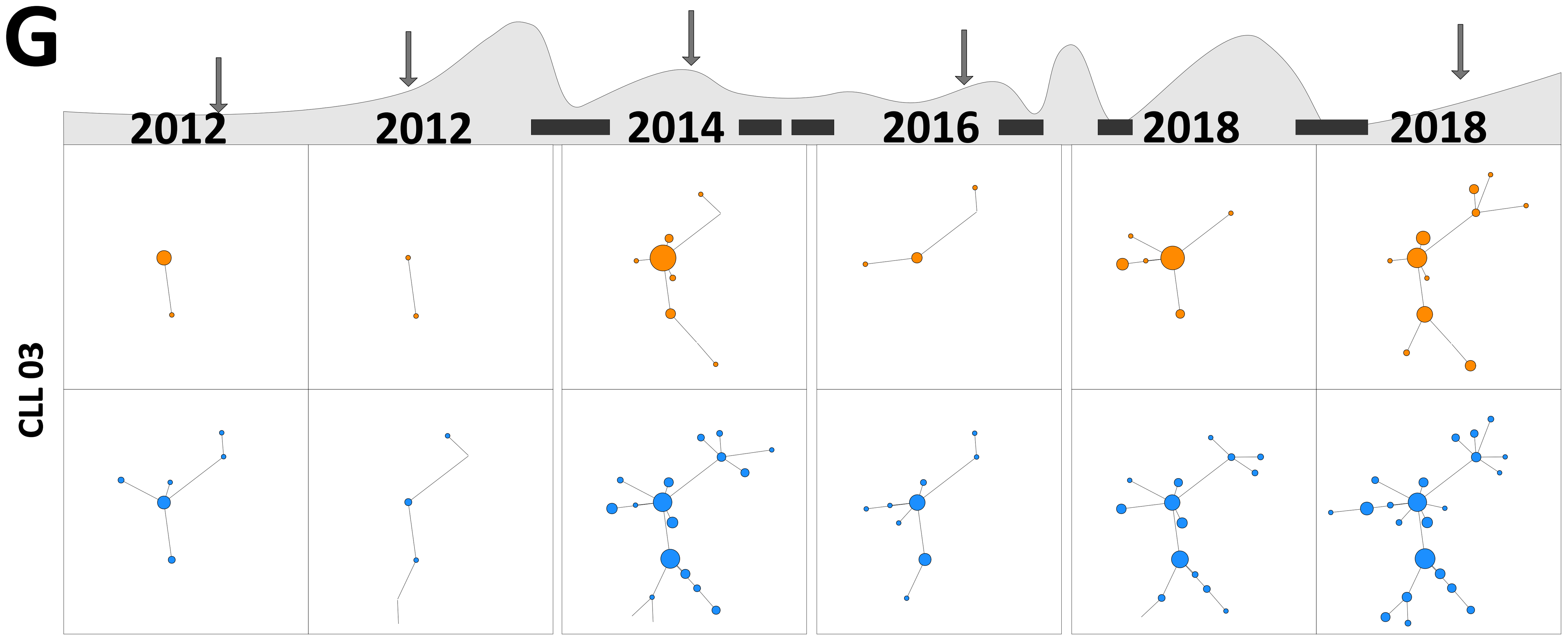


**Figure S4: Longitudinal analysis of IGHV-mutation phylogenetics among CLL subsets.** Representative dendrogram structures of wide-spread IgV-mutation dendrograms from sequential samples of seven patients are shown. Lymphocyte counts are shown as area plots for each patient. Treatment phases are marked as black bars, for treatment information, see Table 4. For each IGHV deep sequencing timepoint (marked by arrow) the number of CLL-derived sequences is identical to the number of sequences displayed from CLL precursor cells. The node size (exponentially displayed, see scale legend) reflects the number of distinct, non-collapsed sequences per node.

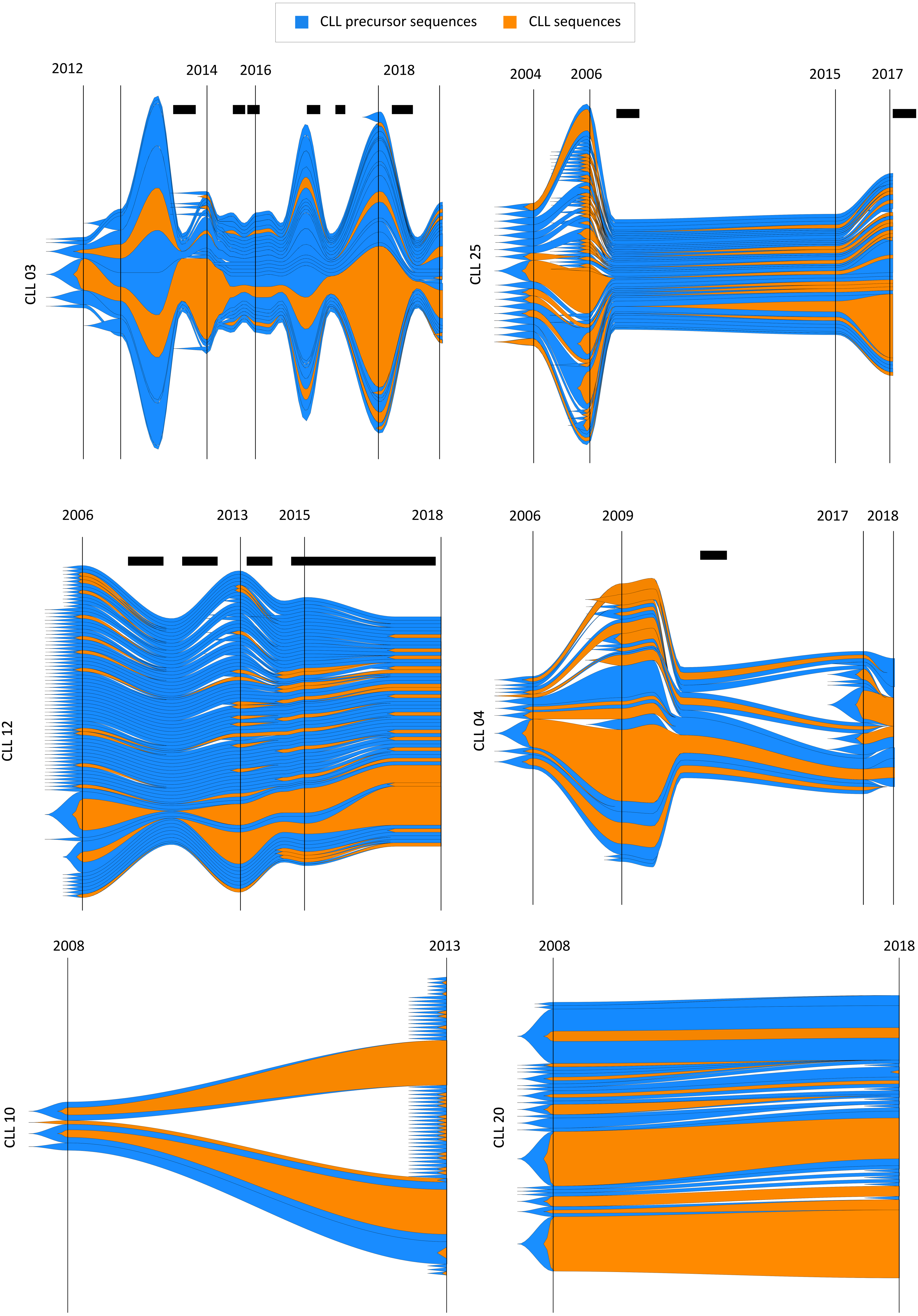


**Figure S5: Longitudinal analysis of IGHV-mutation phylogenetics among CLL subsets.** IgV-mutation dendrograms of CLL precursor- (blue) and CLL-derived IGHV sequences (orange) from longitudinal samples were translated into fish plots to depict subclonal development. Plots were normalized to the lymphocyte count and the number of IGHV sequences per node. Treatment and sampling timepoints are indicated by bars/arrows.

**Table S1: The 856 differential expressed genes between CLL cells and NAC-CLL cells and their normalized expression (batch controlled) values per group as mean value (min-max).** Genes showing an absolute average log2 fold change >= 0.5 (green) are shown in Figure 6A.

| Gene Symbol | CLL cells | NAC-CLL cells | P value | Avg $log_2$ FC | P value adjusted |
|---|---|---|---|---|---|
| ABCA6 | 4 (2.94 - 5.49) | 1.34 (0.51 - 2.19) | <0.001 | 1.46 | <0.001 |
| ABHD14B | 0.86 (0.75 - 0.96) | 1.28 (1.07 - 1.44) | <0.001 | -0.36 | 1 |
| ABI1 | 2.48 (2.03 - 2.63) | 1.85 (1.61 - 2.14) | <0.001 | 0.26 | <0.001 |
| AC007364.1 | 0.5 (0.45 - 0.62) | 0.48 (0.44 - 0.63) | <0.001 | 1 | <0.001 |
| AC007384.1 | 0.97 (0.73 - 1.11) | 0.72 (0.57 - 1.02) | <0.001 | 0.23 | <0.001 |
| AC007569.1 | 0.79 (0.62 - 0.94) | 0.46 (0.3 - 0.62) | <0.001 | 0.48 | <0.001 |
| AC092821.3 | 0.92 (0.68 - 1.19) | 0.5 (0.43 - 0.64) | <0.001 | 0.63 | <0.001 |
| ACBD3 | 1.22 (0.96 - 1.53) | 1.13 (0.95 - 1.45) | 0.03 | -0.31 | 1 |
| ACBD6 | 0.95 (0.77 - 1.09) | 0.84 (0.69 - 1.07) | <0.001 | 0.24 | <0.001 |
| ACSM3 | 2.33 (1.51 - 3.09) | 0.58 (0.17 - 1.35) | <0.001 | 1.37 | <0.001 |
| ACTB | 28.64 (25.79 - 31.96) | 41.25 (36.2 - 45.26) | <0.001 | -0.53 | <0.001 |
| ACTG1 | 8.78 (7.12 - 12.39) | 10.78 (7.93 - 12.58) | <0.001 | -0.44 | <0.001 |
| ACTR2 | 3.59 (3.09 - 4.5) | 4.16 (3.15 - 5.13) | <0.001 | -0.5 | <0.001 |
| ACTR3 | 3.26 (2.99 - 3.74) | 3.67 (3.06 - 3.95) | <0.001 | -0.33 | 1 |
| ADAM17 | 1.08 (0.97 - 1.28) | 0.66 (0.57 - 0.74) | <0.001 | 0.25 | <0.001 |
| ADAM28 | 4.16 (3.11 - 5.45) | 2.25 (1.46 - 2.67) | <0.001 | 1.11 | <0.001 |
| ADD1 | 1.84 (1.3 - 2.27) | 2.17 (1.62 - 2.63) | <0.001 | -0.57 | <0.001 |
| ADD3 | 1.29 (0.75 - 1.94) | 1.76 (1.12 - 2.22) | <0.001 | -0.6 | <0.001 |
| ADK | 1.94 (1.51 - 2.25) | 1.65 (1.33 - 2.05) | <0.001 | 0.39 | <0.001 |
| ADPGK | 1.42 (1.16 - 1.53) | 1.07 (0.95 - 1.34) | <0.001 | 0.32 | <0.001 |
| AFF3 | 6.7 (5.84 - 8.17) | 4.16 (3.39 - 4.8) | <0.001 | 0.72 | <0.001 |
| AFF4 | 3.24 (2.82 - 3.82) | 2.14 (1.89 - 2.56) | <0.001 | 0.5 | <0.001 |
| AGPAT5 | 3.33 (2.48 - 4.38) | 1.17 (0.4 - 1.66) | <0.001 | 1.23 | <0.001 |
| AHCTF1 | 1.15 (0.99 - 1.37) | 1.15 (1.01 - 1.41) | 0.15 | -0.25 | 1 |
| AHNAK | 8.82 (7.36 - 9.58) | 9.58 (8.68 - 10.82) | <0.001 | -0.59 | <0.001 |
| AKAP11 | 1.07 (0.89 - 1.28) | 1.14 (0.91 - 1.37) | 0.47 | -0.23 | 1 |
| AKAP6 | 0.65 (0.42 - 1.19) | 0.37 (0.02 - 0.5) | <0.001 | 0.59 | <0.001 |
| AKAP9 | 2.18 (1.95 - 2.35) | 1.82 (1.62 - 2.09) | <0.001 | 0.24 | <0.001 |
| AKIRIN2 | 1.43 (1.27 - 1.62) | 1.48 (1.3 - 1.64) | 0.42 | -0.21 | 1 |
| AKNA | 1.24 (1.13 - 1.33) | 1.34 (1.13 - 1.47) | <0.001 | -0.32 | 1 |
| AL592429.2 | 1.86 (1.42 - 2.57) | 0.85 (0.47 - 1.12) | <0.001 | 0.92 | <0.001 |
| ALG13 | 1.05 (0.74 - 1.21) | 0.82 (0.72 - 1.1) | <0.001 | 0.23 | <0.001 |
| ALOX5 | 2.25 (1.74 - 3.01) | 1.26 (0.84 - 1.68) | <0.001 | 0.65 | <0.001 |
| AMFR | 1.11 (1.04 - 1.21) | 1.05 (0.87 - 1.23) | <0.001 | -0.31 | 1 |
| ANAPC5 | 1.31 (1.15 - 1.61) | 0.88 (0.63 - 0.98) | <0.001 | 0.46 | <0.001 |

| | | | | | |
|---|---|---|---|---|---|
| ANKLE2 | 1.64 (1.34 - 1.93) | 1.36 (1.05 - 1.64) | <0.001 | 0.31 | <0.001 |
| ANKRD10 | 1.44 (1.28 - 1.66) | 1.17 (1.06 - 1.27) | <0.001 | 0.24 | <0.001 |
| ANKRD11 | 5.78 (4.89 - 7.26) | 4.34 (3.84 - 4.78) | <0.001 | 0.4 | <0.001 |
| ANKRD12 | 5.87 (5.43 - 6.22) | 4.64 (4.17 - 5.01) | <0.001 | 0.36 | <0.001 |
| ANKRD13A | 1.46 (1.22 - 1.78) | 1.81 (1.36 - 2.21) | <0.001 | -0.59 | <0.001 |
| ANKRD36 | 1.29 (0.99 - 1.53) | 1.12 (0.83 - 1.51) | <0.001 | 0.34 | <0.001 |
| ANKRD36C | 1 (0.79 - 1.2) | 0.48 (0.36 - 0.65) | <0.001 | 0.43 | <0.001 |
| ANKRD44 | 3.66 (3.04 - 4.27) | 2.86 (2.55 - 3.47) | <0.001 | 0.49 | <0.001 |
| ANP32B | 2.07 (1.77 - 2.81) | 2.46 (1.87 - 2.77) | 0.02 | -0.36 | 1 |
| ANP32E | 1.42 (1.13 - 1.68) | 1.61 (1.27 - 1.96) | 0.58 | -0.26 | 1 |
| ANXA6 | 0.82 (0.62 - 1.08) | 1.09 (0.8 - 1.55) | <0.001 | -0.51 | <0.001 |
| AP1G1 | 1.85 (1.58 - 2.08) | 1.52 (1.29 - 2.19) | <0.001 | 0.22 | <0.001 |
| AP1S3 | 0.97 (0.93 - 1.01) | 0.58 (0.52 - 0.65) | <0.001 | 0.38 | <0.001 |
| AP3B1 | 1.8 (1.56 - 2.35) | 1.15 (0.85 - 1.27) | <0.001 | 0.35 | <0.001 |
| APBB1IP | 1.36 (1.07 - 1.69) | 1.71 (1.34 - 1.97) | 0.01 | -0.37 | 1 |
| APBB2 | 0.88 (0.57 - 1.23) | 0.5 (0.24 - 0.78) | <0.001 | 0.34 | <0.001 |
| APH1A | 1.11 (0.9 - 1.24) | 1.59 (1.31 - 1.85) | 0.79 | -0.21 | 1 |
| APH1B | 0.82 (0.46 - 0.99) | 0.5 (0.36 - 0.89) | <0.001 | 0.44 | <0.001 |
| APOBEC3G | 0.79 (0.58 - 0.98) | 0.61 (0.53 - 0.77) | <0.001 | 0.3 | <0.001 |
| APOD | 1.56 (1.22 - 2.07) | 1.49 (1.21 - 1.92) | <0.001 | 1.15 | <0.001 |
| APPL1 | 1.93 (1.62 - 2.26) | 1.38 (1.13 - 1.6) | <0.001 | 0.41 | <0.001 |
| ARF6 | 1.92 (1.56 - 2.24) | 2.48 (1.99 - 3.24) | <0.001 | -0.84 | <0.001 |
| ARGLU1 | 5.84 (5.23 - 6.26) | 4.25 (3.94 - 4.58) | <0.001 | 0.52 | <0.001 |
| ARHGAP15 | 1.79 (1.57 - 2.08) | 1.34 (1.11 - 1.51) | <0.001 | 0.4 | <0.001 |
| ARHGAP17 | 2.07 (1.77 - 2.53) | 1.89 (1.55 - 2.32) | 0.63 | -0.25 | 1 |
| ARHGAP24 | 5.48 (4.22 - 6.54) | 3.39 (2.56 - 4.3) | <0.001 | 0.85 | <0.001 |
| ARHGAP30 | 0.88 (0.69 - 1.04) | 1.1 (0.87 - 1.41) | <0.001 | -0.36 | 1 |
| ARHGAP45 | 1.27 (1.23 - 1.4) | 1.49 (1.2 - 1.66) | 0.04 | -0.27 | 1 |
| ARHGDIA | 1.64 (1.56 - 1.85) | 2.21 (1.95 - 2.47) | 0.02 | -0.36 | 1 |
| ARID1A | 1.07 (0.88 - 1.23) | 1.06 (0.89 - 1.23) | 0.19 | -0.24 | 1 |
| ARID1B | 3.08 (2.65 - 3.71) | 2.07 (1.64 - 2.62) | <0.001 | 0.46 | <0.001 |
| ARID4B | 3.23 (2.88 - 3.44) | 2.38 (2.15 - 2.57) | <0.001 | 0.4 | <0.001 |
| ARID5B | 4.68 (3.58 - 6.02) | 3.89 (3.42 - 4.52) | <0.001 | 0.45 | <0.001 |
| ARL5A | 1.05 (0.92 - 1.18) | 1.47 (1.36 - 1.65) | 0.33 | -0.33 | 1 |
| ARL6IP5 | 1.88 (1.56 - 2.39) | 2.39 (1.97 - 2.97) | 0.01 | -0.38 | 1 |
| ASXL1 | 2 (1.76 - 2.36) | 2.04 (1.77 - 2.34) | <0.001 | -0.36 | 1 |
| ATAD2B | 1.58 (1.48 - 1.7) | 1.4 (1.2 - 1.62) | <0.001 | 0.27 | <0.001 |
| ATF7IP | 4.32 (2.75 - 5.62) | 3.8 (2.82 - 5.27) | <0.001 | 0.25 | <0.001 |
| ATP2A3 | 2.13 (1.62 - 2.51) | 2.35 (1.92 - 2.82) | <0.001 | -0.42 | 0.02 |

| | | | | | |
|---|---|---|---|---|---|
| ATP5MC2 | 4.35 (4 - 4.96) | 4.59 (4.2 - 5.37) | <0.001 | 0.22 | <0.001 |
| ATP6V1G1 | 2.53 (2.37 - 2.84) | 3.15 (2.72 - 3.67) | <0.001 | 0.24 | <0.001 |
| ATP9B | 0.92 (0.72 - 1.05) | 0.61 (0.51 - 0.81) | <0.001 | 0.37 | <0.001 |
| ATRX | 2.49 (2.19 - 2.78) | 2.21 (2.01 - 2.49) | <0.001 | 0.25 | <0.001 |
| ATXN1 | 1.29 (1.08 - 1.58) | 0.7 (0.57 - 0.84) | <0.001 | 0.4 | <0.001 |
| ATXN2 | 0.78 (0.66 - 0.87) | 0.53 (0.45 - 0.62) | <0.001 | 0.27 | <0.001 |
| B4GALT1 | 3.23 (2.54 - 4.29) | 3.48 (2.49 - 4.19) | <0.001 | -0.37 | 1 |
| BACH2 | 5.15 (3.85 - 6.55) | 2.35 (1.37 - 3.5) | <0.001 | 1.02 | <0.001 |
| BANK1 | 2.23 (1.44 - 2.81) | 5.23 (4.44 - 6.84) | <0.001 | -1.38 | <0.001 |
| BCAS4 | 1.17 (0.99 - 1.29) | 0.81 (0.74 - 0.87) | <0.001 | 0.24 | <0.001 |
| BCL11A | 3.22 (2.81 - 3.43) | 1.99 (1.9 - 2.11) | <0.001 | 0.54 | <0.001 |
| BCL2 | 5.69 (4.88 - 6.69) | 2.83 (2.13 - 3.46) | <0.001 | 0.86 | <0.001 |
| BICD1 | 1.18 (1.02 - 1.38) | 0.68 (0.59 - 0.79) | <0.001 | 0.54 | <0.001 |
| BIRC3 | 2.54 (1.83 - 3.29) | 2.02 (1.47 - 2.56) | <0.001 | 0.47 | <0.001 |
| BLOC1S1 | 0.91 (0.76 - 1.03) | 0.87 (0.79 - 1.02) | <0.001 | 0.38 | <0.001 |
| BLOC1S2 | 0.88 (0.74 - 1.07) | 0.9 (0.75 - 1.07) | <0.001 | 0.27 | <0.001 |
| BNIP2 | 1.76 (1.61 - 2.08) | 1.97 (1.57 - 2.3) | 0.75 | -0.27 | 1 |
| BNIP3L | 1.08 (0.95 - 1.17) | 0.85 (0.72 - 0.99) | <0.001 | 0.21 | <0.001 |
| BPTF | 2.43 (2.14 - 2.75) | 2.42 (1.93 - 3.05) | 0.02 | -0.24 | 1 |
| BRD2 | 1.83 (1.63 - 1.99) | 1.95 (1.65 - 2.27) | 0.86 | -0.23 | 1 |
| BRD4 | 2.44 (2.14 - 2.75) | 2.5 (2.18 - 2.75) | 0.83 | -0.25 | 1 |
| BRWD1 | 1.62 (1.24 - 1.8) | 1.22 (1.01 - 1.78) | <0.001 | 0.39 | <0.001 |
| BTAF1 | 2.17 (1.98 - 2.33) | 1.49 (1.28 - 1.68) | <0.001 | 0.43 | <0.001 |
| BTG1 | 17.48 (15.35 - 22.01) | 16.1 (14.73 - 18.81) | <0.001 | 0.34 | <0.001 |
| BTLA | 1.73 (1.39 - 2.36) | 1.07 (0.86 - 1.34) | <0.001 | 0.38 | <0.001 |
| C12orf57 | 1.79 (1.57 - 2.25) | 1.62 (1.38 - 1.81) | <0.001 | 0.38 | <0.001 |
| C16orf72 | 1.25 (1.14 - 1.34) | 1.44 (1.25 - 1.75) | <0.001 | -0.35 | 1 |
| CALCOCO1 | 0.78 (0.52 - 0.92) | 0.57 (0.46 - 0.82) | <0.001 | 0.36 | <0.001 |
| CALM1 | 4.64 (4.22 - 5.01) | 6.25 (6.16 - 6.35) | <0.001 | -0.39 | 0.01 |
| CALM2 | 2.85 (2.48 - 3.33) | 3.31 (2.82 - 3.71) | 0.26 | -0.26 | 1 |
| CAMK1D | 0.69 (0.54 - 1.02) | 1.24 (0.88 - 1.45) | <0.001 | -0.64 | <0.001 |
| CAMK2D | 3.19 (2.59 - 3.96) | 1.65 (1.27 - 2.04) | <0.001 | 0.39 | <0.001 |
| CANX | 1.62 (1.42 - 2.2) | 1.92 (1.32 - 2.32) | 0.03 | -0.35 | 1 |
| CAP1 | 1.67 (1.36 - 2.15) | 2.04 (1.6 - 2.52) | <0.001 | -0.36 | 1 |
| CAPN7 | 1.27 (1.07 - 1.84) | 0.89 (0.61 - 1.12) | <0.001 | 0.3 | <0.001 |
| CAPNS1 | 0.81 (0.71 - 1.08) | 1.11 (0.74 - 1.34) | <0.001 | -0.63 | <0.001 |
| CAPZB | 1.48 (1.42 - 1.53) | 2.59 (2.44 - 2.73) | <0.001 | -0.56 | <0.001 |
| CAST | 3.66 (3.21 - 4.15) | 3.25 (2.84 - 3.77) | <0.001 | 0.3 | <0.001 |
| CCDC12 | 1.67 (1.47 - 1.99) | 1.22 (0.92 - 1.41) | <0.001 | 0.37 | <0.001 |

| | | | | | |
|---|---|---|---|---|---|
| CCDC141 | 2.14 (1.72 - 2.56) | 0.77 (0.5 - 1.11) | <0.001 | 0.9 | <0.001 |
| CCDC18-AS1 | 2 (1.36 - 2.57) | 1.14 (0.79 - 1.81) | <0.001 | 0.74 | <0.001 |
| CCDC32 | 1.02 (0.89 - 1.23) | 0.85 (0.7 - 0.98) | <0.001 | 0.24 | <0.001 |
| CCDC50 | 2.01 (1.74 - 2.56) | 2.81 (2.32 - 3.06) | <0.001 | -0.44 | 1 |
| CCDC6 | 2.37 (1.98 - 2.78) | 2.49 (1.88 - 3.25) | 0.07 | -0.36 | 1 |
| CCDC88A | 5.96 (4.83 - 7.11) | 2.23 (1.92 - 2.75) | <0.001 | 1.22 | <0.001 |
| CCDC91 | 0.78 (0.71 - 0.84) | 0.56 (0.48 - 0.66) | <0.001 | 0.35 | <0.001 |
| CCDC93 | 1.27 (1.07 - 1.73) | 1.03 (0.87 - 1.26) | <0.001 | 0.36 | <0.001 |
| CCND3 | 2.46 (1.91 - 3.18) | 2 (1.57 - 2.35) | <0.001 | 0.23 | <0.001 |
| CCNG2 | 0.79 (0.49 - 1) | 1.03 (0.71 - 1.43) | <0.001 | -0.65 | <0.001 |
| CCNH | 1.07 (0.95 - 1.26) | 0.62 (0.5 - 0.75) | <0.001 | 0.34 | <0.001 |
| CCNI | 4.42 (3.85 - 5.27) | 5.58 (4.74 - 6.14) | <0.001 | -0.41 | <0.001 |
| CCNL1 | 4.89 (4.22 - 5.94) | 3.28 (2.93 - 3.65) | <0.001 | 0.55 | <0.001 |
| CCNT2 | 1.31 (1.22 - 1.35) | 0.97 (0.87 - 1.22) | <0.001 | 0.21 | <0.001 |
| CCR7 | 3.81 (3.03 - 5.39) | 2.22 (1.62 - 2.6) | <0.001 | 1.45 | <0.001 |
| CD164 | 2.36 (2.08 - 2.65) | 2.62 (2.21 - 3.08) | <0.001 | -0.42 | 0.39 |
| CD200 | 1.39 (1.19 - 1.65) | 0.6 (0.47 - 0.83) | <0.001 | 0.4 | <0.001 |
| CD22 | 1.07 (0.7 - 1.62) | 1.32 (0.69 - 2) | <0.001 | -0.65 | <0.001 |
| CD27 | 1.71 (1.48 - 2.37) | 1.7 (1.36 - 2.18) | <0.001 | 0.43 | <0.001 |
| CD40 | 0.91 (0.76 - 1.06) | 0.91 (0.79 - 1.12) | <0.001 | 0.24 | <0.001 |
| CD46 | 1.68 (1.44 - 1.91) | 1.21 (0.99 - 1.39) | <0.001 | 0.31 | <0.001 |
| CD48 | 1.75 (1.53 - 2.08) | 2.35 (2.06 - 2.57) | 0.07 | -0.33 | 1 |
| CD53 | 2.48 (2.09 - 2.77) | 3.09 (2.62 - 3.98) | <0.001 | -0.49 | 0.02 |
| CD55 | 2.79 (2.16 - 3.55) | 2.26 (1.78 - 3.06) | <0.001 | 0.28 | <0.001 |
| CD81 | 0.72 (0.44 - 1.02) | 1.27 (0.95 - 1.69) | <0.001 | -1.08 | <0.001 |
| CD82 | 1.09 (0.86 - 1.49) | 1.65 (1.35 - 1.87) | 0.02 | -0.32 | 1 |
| CD84 | 1.51 (1.26 - 1.95) | 1.04 (0.74 - 1.51) | <0.001 | 0.31 | <0.001 |
| CD99 | 2.17 (1.81 - 2.57) | 2.57 (2.17 - 3.14) | 0.8 | -0.26 | 1 |
| CDCA7L | 1.27 (1.03 - 1.64) | 1.16 (0.96 - 1.43) | <0.001 | 0.22 | <0.001 |
| CDK11A | 1.07 (0.85 - 1.33) | 0.8 (0.65 - 1.02) | <0.001 | 0.37 | <0.001 |
| CDK13 | 2.1 (1.85 - 2.28) | 1.83 (1.69 - 2.16) | <0.001 | 0.22 | <0.001 |
| CDK14 | 4.15 (3.36 - 4.85) | 2.22 (1.87 - 2.86) | <0.001 | 0.77 | <0.001 |
| CDK5RAP3 | 1.01 (0.7 - 1.16) | 0.69 (0.57 - 0.99) | <0.001 | 0.26 | <0.001 |
| CDV3 | 2.1 (1.65 - 2.46) | 2.99 (2.32 - 3.41) | <0.001 | -0.92 | <0.001 |
| CEMIP2 | 3.32 (2.33 - 4.37) | 2.07 (1.54 - 2.36) | <0.001 | 0.4 | <0.001 |
| CENPC | 1.7 (1.19 - 2.13) | 1.33 (1.04 - 2) | <0.001 | 0.4 | <0.001 |
| CEP350 | 1.78 (1.59 - 1.95) | 1.67 (1.44 - 2) | 0.47 | -0.25 | 1 |
| CEP95 | 1.7 (1.39 - 2.07) | 1.04 (0.88 - 1.29) | <0.001 | 0.46 | <0.001 |
| CFL1 | 4.97 (4.41 - 5.44) | 6.65 (6.11 - 7.2) | <0.001 | -0.31 | 1 |

| | | | | | |
|---|---|---|---|---|---|
| CHD2 | 5.4 (4.99 - 5.69) | 3.72 (3.29 - 4.23) | <0.001 | 0.61 | <0.001 |
| CIB1 | 1.14 (0.74 - 1.65) | 1.53 (0.91 - 2.09) | 0.58 | -0.34 | 1 |
| CIITA | 2.23 (1.93 - 2.61) | 2.23 (1.86 - 2.59) | 0.59 | -0.25 | 1 |
| CLEC17A | 1 (0.62 - 1.31) | 0.67 (0.52 - 0.99) | <0.001 | 0.36 | <0.001 |
| CLIC1 | 2.37 (2.16 - 2.77) | 2.95 (2.31 - 3.49) | 0.28 | -0.27 | 1 |
| CLK1 | 1.99 (1.64 - 2.34) | 1.57 (1.41 - 1.69) | <0.001 | 0.2 | <0.001 |
| CLK4 | 0.93 (0.8 - 1.04) | 0.7 (0.59 - 0.82) | <0.001 | 0.26 | <0.001 |
| CLNK | 1.57 (1.26 - 1.93) | 0.35 (0.16 - 0.52) | <0.001 | 1.08 | <0.001 |
| CLTC | 1.27 (1.1 - 1.44) | 1.33 (1.05 - 1.48) | 0.89 | -0.21 | 1 |
| CMPK1 | 1.14 (0.76 - 1.34) | 1.64 (1.31 - 2.17) | <0.001 | -0.58 | <0.001 |
| CMTM6 | 0.86 (0.62 - 1.4) | 1.37 (0.73 - 1.65) | <0.001 | -0.59 | <0.001 |
| CNOT1 | 1.41 (1.3 - 1.55) | 1.51 (1.29 - 1.65) | 0.69 | -0.22 | 1 |
| CNOT2 | 1.59 (1.49 - 1.77) | 1.21 (1.01 - 1.41) | <0.001 | 0.36 | <0.001 |
| CNOT4 | 1.54 (1.34 - 1.7) | 0.94 (0.79 - 1.16) | <0.001 | 0.51 | <0.001 |
| CNST | 1.82 (1.42 - 2.1) | 1.51 (1.21 - 1.99) | <0.001 | 0.35 | <0.001 |
| COBLL1 | 1.62 (1.21 - 2.12) | 2.64 (2.26 - 2.98) | 0.13 | -0.27 | 1 |
| COCH | 1.24 (0.61 - 2.46) | 1.04 (0.44 - 1.71) | <0.001 | 0.75 | <0.001 |
| COL19A1 | 0.98 (0.62 - 1.38) | 0.42 (0.17 - 0.72) | <0.001 | 0.81 | <0.001 |
| COL9A3 | 0.81 (0.68 - 0.94) | 0.44 (0.29 - 0.54) | <0.001 | 0.47 | <0.001 |
| CORO1A | 3.1 (2.92 - 3.38) | 4.93 (4.31 - 5.4) | <0.001 | -0.33 | <0.001 |
| COTL1 | 2.64 (2.07 - 3.3) | 3.77 (3.18 - 4.39) | <0.001 | -0.67 | <0.001 |
| COX4I1 | 7.66 (7.04 - 9.67) | 7.87 (7.24 - 9.02) | <0.001 | 0.26 | <0.001 |
| COX6A1 | 1.77 (1.57 - 2.08) | 1.87 (1.68 - 2.1) | <0.001 | 0.26 | <0.001 |
| CPNE1 | 0.8 (0.66 - 0.93) | 0.7 (0.59 - 0.86) | <0.001 | 0.27 | <0.001 |
| CREB3L2 | 3.08 (2.49 - 4.32) | 1.54 (1.08 - 1.89) | <0.001 | 0.49 | <0.001 |
| CREBZF | 1.23 (0.98 - 1.47) | 0.86 (0.66 - 1.07) | <0.001 | 0.27 | <0.001 |
| CRIP1 | 2.07 (1.25 - 2.79) | 4.32 (3.25 - 5.64) | <0.001 | -1.47 | <0.001 |
| CSDE1 | 3.81 (3.03 - 4.75) | 3.92 (2.9 - 4.77) | 0.01 | -0.32 | 1 |
| CSK | 1.64 (1.46 - 1.9) | 1.95 (1.57 - 2.35) | <0.001 | -0.57 | <0.001 |
| CSNK1G3 | 1.67 (1.05 - 2.02) | 1.62 (1.29 - 2.51) | <0.001 | 0.25 | <0.001 |
| CTDSPL2 | 1.13 (1.01 - 1.21) | 0.8 (0.69 - 0.89) | <0.001 | 0.21 | <0.001 |
| CTNNB1 | 1.59 (1.48 - 1.72) | 1.54 (1.31 - 1.73) | 0.59 | -0.25 | 1 |
| CTSZ | 1.33 (1.14 - 1.75) | 1.45 (1.25 - 1.72) | <0.001 | -0.34 | 1 |
| CUL5 | 1.19 (1 - 1.37) | 1.06 (0.92 - 1.38) | <0.001 | 0.22 | <0.001 |
| CWF19L2 | 1.08 (0.82 - 1.24) | 0.75 (0.62 - 0.88) | <0.001 | 0.31 | <0.001 |
| CXCR4 | 6.49 (4.99 - 7.63) | 5.55 (4.95 - 6.19) | <0.001 | 0.32 | <0.001 |
| CXXC5 | 2.54 (2.35 - 2.79) | 2.56 (2.23 - 2.92) | 0.76 | -0.21 | 1 |
| CYBA | 3.32 (3.04 - 3.54) | 4.48 (4.12 - 4.98) | <0.001 | -0.44 | <0.001 |
| CYBB | 0.51 (0.11 - 0.84) | 1.41 (1 - 1.89) | <0.001 | -0.87 | <0.001 |

| | | | | | |
|---|---|---|---|---|---|
| CYFIP2 | 2.56 (2.06 - 2.94) | 2.45 (2 - 2.85) | 0.34 | -0.25 | 1 |
| CYTH1 | 3.46 (3.16 - 3.84) | 2.68 (2.21 - 3.22) | <0.001 | 0.28 | <0.001 |
| DAPP1 | 0.93 (0.76 - 1.32) | 1.66 (1.31 - 2.13) | <0.001 | -0.51 | <0.001 |
| DBI | 1.94 (1.55 - 2.3) | 1.96 (1.55 - 2.16) | <0.001 | 0.42 | <0.001 |
| DBNL | 1.39 (1.21 - 1.6) | 1.62 (1.39 - 2.12) | 0.02 | -0.32 | 1 |
| DCAF8 | 1.51 (1.15 - 1.66) | 1.12 (0.95 - 1.56) | <0.001 | 0.34 | <0.001 |
| DCLK2 | 0.9 (0.61 - 1.06) | 0.36 (0.21 - 0.56) | <0.001 | 0.64 | <0.001 |
| DDHD1 | 1.68 (1.47 - 1.78) | 0.95 (0.83 - 1.11) | <0.001 | 0.55 | <0.001 |
| DDX17 | 7.48 (6.38 - 9.38) | 5.74 (5.4 - 6.51) | <0.001 | 0.33 | <0.001 |
| DDX24 | 4.25 (3.8 - 4.77) | 3.8 (3.53 - 4.19) | <0.001 | 0.2 | <0.001 |
| DDX27 | 1.12 (0.92 - 1.38) | 0.79 (0.63 - 0.89) | <0.001 | 0.26 | <0.001 |
| DDX39B | 3.05 (2.47 - 3.3) | 2.26 (1.87 - 2.8) | <0.001 | 0.26 | <0.001 |
| DDX5 | 14.03 (12.93 - 16.19) | 12.37 (10 - 13.74) | <0.001 | 0.23 | <0.001 |
| DEK | 1.12 (1.02 - 1.27) | 1.28 (1.11 - 1.39) | 0.86 | -0.2 | 1 |
| DENND2D | 0.67 (0.51 - 0.91) | 0.47 (0.34 - 0.54) | <0.001 | 0.4 | <0.001 |
| DGKA | 1.7 (1.28 - 1.91) | 0.71 (0.57 - 0.93) | <0.001 | 0.9 | <0.001 |
| DGKD | 1.46 (1.31 - 1.82) | 1.74 (1.27 - 1.92) | 0.01 | -0.36 | 1 |
| DGKG | 1.07 (0.69 - 1.67) | 0.47 (0.2 - 0.74) | <0.001 | 0.79 | <0.001 |
| DIP2B | 1.02 (0.82 - 1.35) | 0.95 (0.65 - 1.2) | <0.001 | 0.22 | <0.001 |
| DLGAP4 | 0.82 (0.63 - 1.16) | 0.68 (0.41 - 0.92) | <0.001 | 0.28 | <0.001 |
| DMD | 1.48 (0.97 - 2.2) | 0.56 (0.15 - 0.97) | <0.001 | 1.4 | <0.001 |
| DMTF1 | 1.67 (1.43 - 1.88) | 1.2 (1.03 - 1.5) | <0.001 | 0.31 | <0.001 |
| DNAJC15 | 0.84 (0.64 - 0.99) | 0.7 (0.55 - 0.97) | <0.001 | 0.23 | <0.001 |
| DNMBP | 2.43 (1.93 - 3.26) | 0.82 (0.45 - 1.22) | <0.001 | 0.94 | <0.001 |
| DOCK2 | 1.28 (1.21 - 1.45) | 1.49 (1.22 - 1.8) | <0.001 | -0.38 | 1 |
| DRAM2 | 1.34 (0.96 - 1.65) | 1.1 (0.9 - 1.42) | <0.001 | 0.39 | <0.001 |
| DUSP1 | 2.06 (1.72 - 2.72) | 2.68 (2.08 - 3.52) | 0.01 | -0.32 | 1 |
| DYM | 1.05 (0.88 - 1.25) | 0.94 (0.72 - 1.22) | <0.001 | 0.23 | <0.001 |
| DYNLL1 | 1.05 (0.98 - 1.24) | 1.01 (0.82 - 1.27) | <0.001 | 0.22 | <0.001 |
| EAPP | 0.99 (0.9 - 1.18) | 0.76 (0.68 - 0.84) | <0.001 | 0.26 | <0.001 |
| EBLN3P | 2.25 (2.2 - 2.39) | 1.69 (1.57 - 1.79) | <0.001 | 0.28 | <0.001 |
| EEF1A1 | 95.13 (91.17 - 111.59) | 114.53 (108.22 - 121.79) | <0.001 | -0.23 | <0.001 |
| EEF1B2 | 12.38 (11.33 - 14.01) | 15.26 (14.02 - 17.01) | <0.001 | -0.26 | <0.001 |
| EEF2 | 14.76 (12.85 - 18.28) | 18.03 (13.27 - 21.87) | <0.001 | -0.81 | <0.001 |
| EHMT1 | 2.49 (2.27 - 2.88) | 1.46 (1.13 - 1.71) | <0.001 | 0.7 | <0.001 |
| EIF2AK1 | 1.01 (0.76 - 1.4) | 1.22 (0.77 - 1.5) | 0.61 | -0.25 | 1 |
| EIF2S3 | 2.19 (1.99 - 2.57) | 2.47 (2 - 2.93) | 0.17 | -0.32 | 1 |
| EIF3L | 2.99 (2.6 - 3.47) | 3.52 (3.22 - 3.76) | 0.48 | -0.25 | 1 |

| | | | | | |
|---|---|---|---|---|---|
| EIF4A1 | 1.8 (1.56 - 2.33) | 2.24 (1.67 - 2.63) | 0.77 | -0.21 | 1 |
| EIF4H | 2.22 (2.07 - 2.62) | 2.78 (2.18 - 3.39) | <0.001 | -0.51 | <0.001 |
| ELF2 | 1.58 (1.44 - 1.76) | 1.21 (1.07 - 1.45) | <0.001 | 0.33 | <0.001 |
| ELK4 | 1.16 (1.03 - 1.38) | 1.62 (1.25 - 1.98) | <0.001 | -0.75 | <0.001 |
| ELP2 | 1.03 (0.7 - 1.21) | 0.82 (0.62 - 1.2) | <0.001 | 0.22 | <0.001 |
| EMB | 0.74 (0.61 - 0.98) | 1.34 (1.09 - 1.46) | <0.001 | -0.62 | <0.001 |
| EMP3 | 3.58 (3.14 - 4.16) | 5.02 (4.48 - 5.45) | <0.001 | -0.4 | 0.01 |
| ENO1 | 1.38 (1.1 - 1.59) | 1.9 (1.47 - 2.3) | <0.001 | -0.35 | 1 |
| ENTPD4 | 1.43 (1.19 - 1.7) | 1.04 (0.91 - 1.27) | <0.001 | 0.22 | <0.001 |
| EP400 | 1.46 (1.2 - 1.57) | 1.02 (0.89 - 1.21) | <0.001 | 0.3 | <0.001 |
| EPB41 | 2.98 (1.98 - 3.7) | 2.43 (1.95 - 3.27) | <0.001 | 0.43 | <0.001 |
| EPC1 | 3.46 (2.88 - 3.66) | 3.3 (2.78 - 4.36) | <0.001 | 0.31 | <0.001 |
| ERC1 | 1.24 (1.11 - 1.38) | 0.77 (0.64 - 0.85) | <0.001 | 0.51 | <0.001 |
| ERICH1 | 1.06 (0.74 - 1.19) | 0.89 (0.76 - 1.28) | <0.001 | 0.34 | <0.001 |
| ETS1 | 3.28 (2.53 - 3.93) | 3.14 (2.39 - 4.34) | <0.001 | -0.63 | <0.001 |
| ETV6 | 1.5 (1.19 - 2.12) | 0.76 (0.4 - 1.04) | <0.001 | 0.5 | <0.001 |
| EVI2B | 3.42 (2.87 - 4.23) | 4.44 (3.65 - 4.97) | <0.001 | -0.51 | <0.001 |
| EVL | 2.11 (1.83 - 2.44) | 1.54 (1.28 - 1.75) | <0.001 | 0.43 | <0.001 |
| EWSR1 | 2.67 (2.15 - 2.91) | 2.25 (1.98 - 2.76) | <0.001 | 0.26 | <0.001 |
| EXOC4 | 1 (0.89 - 1.16) | 0.64 (0.57 - 0.7) | <0.001 | 0.3 | <0.001 |
| EXOSC10 | 1.08 (0.96 - 1.24) | 0.9 (0.79 - 1.02) | <0.001 | 0.24 | <0.001 |
| EZH1 | 1.18 (0.82 - 1.29) | 0.83 (0.65 - 1.16) | <0.001 | 0.3 | <0.001 |
| FAM102A | 1.18 (0.84 - 1.88) | 1.33 (1.01 - 1.62) | 0.01 | -0.3 | 1 |
| FAM107B | 1.33 (1.18 - 1.57) | 1.56 (1.26 - 1.99) | <0.001 | -0.74 | <0.001 |
| FAM117B | 0.93 (0.87 - 1.01) | 0.9 (0.78 - 1.02) | 0.29 | -0.25 | 1 |
| FAM133B | 1.26 (1.11 - 1.46) | 1.23 (1.01 - 1.56) | <0.001 | 0.26 | <0.001 |
| FAM13B | 1.36 (1.26 - 1.51) | 0.96 (0.8 - 1.12) | <0.001 | 0.25 | <0.001 |
| FAM30A | 0.69 (0.39 - 1.35) | 0.86 (0.75 - 0.99) | <0.001 | -0.36 | 1 |
| FAM49A | 0.46 (0.12 - 1.09) | 1.69 (0.99 - 2.33) | <0.001 | -0.98 | <0.001 |
| FAM49B | 1.1 (1.01 - 1.33) | 1.43 (1.21 - 1.52) | 0.8 | -0.25 | 1 |
| FBL | 1.18 (1.06 - 1.27) | 1.34 (1.23 - 1.45) | 0.49 | -0.21 | 1 |
| FBXO21 | 2.65 (1.94 - 3.53) | 1.62 (1.21 - 1.93) | <0.001 | 0.62 | <0.001 |
| FBXO9 | 0.86 (0.54 - 1.05) | 0.8 (0.62 - 1.16) | <0.001 | 0.22 | <0.001 |
| FCER2 | 4.82 (3.91 - 6.14) | 2.7 (2.08 - 3.27) | <0.001 | 0.91 | <0.001 |
| FCGR2B | 1.38 (1.09 - 1.89) | 1.54 (1.15 - 2.4) | 0.02 | -0.34 | 1 |
| FCMR | 8.87 (7.11 - 10.79) | 6.83 (5.9 - 8.37) | <0.001 | 0.34 | <0.001 |
| FCRL2 | 1.69 (1.37 - 2.24) | 1.34 (1.16 - 1.54) | <0.001 | 0.42 | <0.001 |
| FCRL3 | 1.6 (1.1 - 2.69) | 1.47 (0.58 - 2.42) | <0.001 | 0.38 | <0.001 |
| FCRL5 | 2.52 (1.78 - 3.87) | 2.14 (0.92 - 3.83) | <0.001 | 0.33 | <0.001 |

| | | | | | |
|---|---|---|---|---|---|
| FDFT1 | 0.92 (0.81 - 1.06) | 1.04 (0.87 - 1.33) | <0.001 | -0.44 | 1 |
| FGD2 | 0.87 (0.67 - 1.06) | 1.09 (0.87 - 1.52) | <0.001 | -0.35 | 1 |
| FKBP8 | 1.03 (0.9 - 1.16) | 0.97 (0.82 - 1.18) | 0.63 | -0.23 | 1 |
| FLI1 | 0.91 (0.66 - 1.24) | 0.9 (0.62 - 1.39) | 0.85 | -0.2 | 1 |
| FLNA | 1.68 (1.32 - 2.23) | 2.5 (1.83 - 3.04) | <0.001 | -0.77 | <0.001 |
| FLOT2 | 0.89 (0.57 - 1.21) | 1.13 (0.77 - 1.39) | 0.05 | -0.34 | 1 |
| FMNL1 | 1.31 (1.19 - 1.42) | 1.57 (1.33 - 1.78) | <0.001 | -0.44 | <0.001 |
| FMOD | 1.22 (1.02 - 1.46) | 0.57 (0.44 - 0.71) | <0.001 | 0.42 | <0.001 |
| FNBP1 | 1.79 (1.68 - 2) | 2.65 (2.34 - 2.97) | <0.001 | -0.66 | <0.001 |
| FNBP4 | 3.7 (3.32 - 4.11) | 2.95 (2.68 - 3.69) | <0.001 | 0.37 | <0.001 |
| FNDC3A | 0.93 (0.72 - 1.08) | 0.51 (0.42 - 0.72) | <0.001 | 0.45 | <0.001 |
| FOXP1 | 10.84 (9.81 - 11.81) | 6.84 (5.85 - 7.45) | <0.001 | 0.73 | <0.001 |
| FRY | 0.95 (0.77 - 1.36) | 0.38 (0.2 - 0.52) | <0.001 | 0.65 | <0.001 |
| FRYL | 1.16 (0.86 - 1.31) | 0.86 (0.74 - 1.12) | <0.001 | 0.26 | <0.001 |
| FTH1 | 15.59 (14.6 - 17.68) | 21.15 (18.72 - 26.64) | <0.001 | -0.25 | <0.001 |
| FTL | 16.22 (15.28 - 18.58) | 17.03 (16.08 - 18.4) | <0.001 | 0.29 | <0.001 |
| FTX | 1.56 (1.23 - 1.93) | 1.01 (0.86 - 1.27) | <0.001 | 0.33 | <0.001 |
| FUS | 5.67 (5.12 - 6.21) | 4.89 (4.46 - 5.9) | <0.001 | 0.59 | <0.001 |
| FXYD5 | 4.61 (4.07 - 6.17) | 4.12 (3.62 - 5.14) | <0.001 | 0.62 | <0.001 |
| GAB1 | 1.92 (1.63 - 2.49) | 0.65 (0.51 - 0.76) | <0.001 | 0.89 | <0.001 |
| GAB2 | 1.3 (0.96 - 1.79) | 0.68 (0.3 - 0.95) | <0.001 | 0.33 | <0.001 |
| GABPB1-AS1 | 1.69 (1.29 - 2.44) | 0.97 (0.64 - 1.32) | <0.001 | 0.73 | <0.001 |
| GGA1 | 0.85 (0.55 - 1.05) | 0.58 (0.42 - 0.98) | <0.001 | 0.5 | <0.001 |
| GGA2 | 4.33 (3.41 - 6.03) | 3.21 (2.45 - 4.03) | <0.001 | 0.32 | <0.001 |
| GHITM | 0.86 (0.81 - 0.93) | 1.1 (0.96 - 1.19) | 0.45 | -0.22 | 1 |
| GIT2 | 1.27 (1.08 - 1.53) | 0.89 (0.71 - 1.08) | <0.001 | 0.35 | <0.001 |
| GLS | 2.73 (2.51 - 3.1) | 2.56 (2.28 - 2.88) | 0.02 | -0.28 | 1 |
| GMFG | 1.5 (1.23 - 1.7) | 1.49 (1.38 - 1.66) | <0.001 | 0.22 | <0.001 |
| GNA13 | 2.22 (1.89 - 2.53) | 2.54 (2.16 - 3.03) | <0.001 | -0.42 | 0.01 |
| GNAI2 | 1.7 (1.51 - 2.05) | 2.23 (1.66 - 2.64) | <0.001 | -0.83 | <0.001 |
| GNAS | 5.61 (4.8 - 6.82) | 4.81 (4.43 - 5.29) | 0.03 | -0.21 | 1 |
| GNB1 | 1.39 (1.21 - 1.49) | 1.66 (1.41 - 1.99) | <0.001 | -0.61 | <0.001 |
| GNG2 | 2.01 (1.84 - 2.36) | 1.48 (1.32 - 1.73) | <0.001 | 0.23 | <0.001 |
| GNPTAB | 2.17 (1.64 - 2.51) | 1.22 (1.09 - 1.67) | <0.001 | 0.58 | <0.001 |
| GOLGA4 | 2.1 (1.93 - 2.43) | 1.53 (1.36 - 1.69) | <0.001 | 0.34 | <0.001 |
| GOLGB1 | 1.58 (1.43 - 1.77) | 1.21 (1.02 - 1.47) | <0.001 | 0.24 | <0.001 |
| GON4L | 1.05 (0.96 - 1.15) | 0.64 (0.56 - 0.74) | <0.001 | 0.32 | <0.001 |
| GPBP1 | 2.46 (2.16 - 2.69) | 2.22 (1.94 - 2.69) | <0.001 | 0.34 | <0.001 |
| GPCPD1 | 1.59 (1.54 - 1.65) | 1.16 (1.11 - 1.27) | <0.001 | 0.31 | <0.001 |

| | | | | | |
|---|---|---|---|---|---|
| GPR183 | 0.74 (0.55 - 1.12) | 1.92 (1.52 - 2.43) | <0.001 | -0.91 | <0.001 |
| GPT2 | 1.13 (0.83 - 1.52) | 0.37 (0.06 - 0.66) | <0.001 | 0.69 | <0.001 |
| GRK2 | 1.05 (0.82 - 1.41) | 1.1 (0.9 - 1.46) | <0.001 | -0.39 | 0.01 |
| GRN | 1.07 (0.72 - 1.78) | 1.31 (0.82 - 1.73) | 0.49 | -0.21 | 1 |
| GSAP | 1.46 (1.32 - 1.63) | 0.83 (0.68 - 0.93) | <0.001 | 0.51 | <0.001 |
| GSPT1 | 1.01 (0.96 - 1.11) | 1.06 (0.84 - 1.23) | 0.01 | -0.33 | 1 |
| GSTK1 | 1.07 (0.94 - 1.33) | 1.12 (0.86 - 1.41) | 0.55 | -0.27 | 1 |
| GSTP1 | 1.4 (1.18 - 1.98) | 1.19 (0.75 - 1.51) | <0.001 | 0.6 | <0.001 |
| H2AFY | 1.02 (0.91 - 1.12) | 0.86 (0.73 - 0.91) | <0.001 | 0.35 | <0.001 |
| H3F3B | 22.4 (19.16 - 29.73) | 18.09 (14.63 - 21.31) | <0.001 | 0.6 | <0.001 |
| HEXA | 0.92 (0.79 - 1.06) | 0.71 (0.57 - 0.89) | <0.001 | 0.34 | <0.001 |
| HIF1A | 1.01 (0.5 - 1.64) | 1.97 (1.33 - 2.79) | <0.001 | -0.44 | <0.001 |
| HIPK2 | 0.83 (0.65 - 0.99) | 1.4 (1.09 - 1.68) | <0.001 | -0.59 | <0.001 |
| HIST1H1C | 4.95 (3.75 - 6.14) | 3.69 (3.11 - 4.81) | <0.001 | 0.27 | <0.001 |
| HIST1H1D | 2.48 (1.46 - 4.26) | 2.63 (1.22 - 3.63) | <0.001 | 0.45 | <0.001 |
| HIST1H1E | 12.39 (10.5 - 14.04) | 9.23 (7.35 - 11.24) | <0.001 | 0.54 | <0.001 |
| HIST1H4C | 2.76 (2.38 - 3.12) | 2.89 (2.5 - 3.36) | <0.001 | 0.22 | <0.001 |
| HIVEP1 | 1.95 (1.74 - 2.25) | 1.3 (1.15 - 1.48) | <0.001 | 0.44 | <0.001 |
| HIVEP2 | 1.05 (0.85 - 1.37) | 0.64 (0.46 - 0.87) | <0.001 | 0.29 | <0.001 |
| HMGN3 | 0.87 (0.7 - 0.99) | 0.78 (0.58 - 1.13) | <0.001 | 0.42 | <0.001 |
| HNRNPA1 | 7.62 (7.05 - 8.33) | 9.67 (8.71 - 10.43) | <0.001 | -0.24 | 1 |
| HNRNPA2B1 | 12.06 (10.84 - 12.94) | 10.41 (9.57 - 11.34) | <0.001 | 0.22 | <0.001 |
| HNRNPD | 2.53 (2.38 - 2.62) | 2.74 (2.56 - 3.19) | <0.001 | -0.29 | 1 |
| HNRNPF | 1.28 (1.05 - 1.75) | 1.51 (1.09 - 1.87) | 0.8 | -0.26 | 1 |
| HNRNPK | 3.83 (3.45 - 4.48) | 4.14 (3.35 - 5.02) | 0.37 | -0.23 | 1 |
| HNRNPL | 2.52 (2.3 - 2.73) | 2.87 (2.38 - 3.53) | <0.001 | -0.5 | <0.001 |
| HNRNPM | 1.86 (1.61 - 2.21) | 1.95 (1.5 - 2.16) | 0.16 | -0.21 | 1 |
| HNRNPUL1 | 1.26 (1.14 - 1.39) | 1.53 (1.3 - 1.8) | <0.001 | -0.37 | 1 |
| HNRNPUL2 | 1.15 (1 - 1.31) | 1.34 (1.03 - 1.56) | <0.001 | -0.52 | <0.001 |
| HSD17B4 | 0.94 (0.7 - 1.29) | 0.52 (0.32 - 0.65) | <0.001 | 0.39 | <0.001 |
| HSP90AB1 | 9.59 (8.48 - 11.88) | 11.49 (9.02 - 14.19) | <0.001 | -0.3 | 0.16 |
| HSP90B1 | 2.63 (1.93 - 3.44) | 3.22 (2.41 - 4.17) | 0.42 | -0.27 | 1 |
| HSPA8 | 4.53 (3.76 - 5.87) | 5.1 (3.91 - 6.51) | <0.001 | -0.39 | <0.001 |
| HUWE1 | 1.66 (1.55 - 1.89) | 1.8 (1.43 - 2) | 0.16 | -0.3 | 1 |
| HVCN1 | 2.74 (1.93 - 4.04) | 2.27 (1.76 - 3.04) | <0.001 | 0.69 | <0.001 |
| ID3 | 2.67 (2.29 - 3.04) | 1.67 (1.29 - 2.25) | <0.001 | 0.29 | <0.001 |
| IDS | 1.25 (1.07 - 1.48) | 1.74 (1.41 - 2.08) | <0.001 | -0.47 | 0.05 |
| IER2 | 1.07 (0.85 - 1.29) | 1.84 (1.53 - 2.27) | <0.001 | -0.62 | <0.001 |
| IFI16 | 1.98 (1.72 - 2.26) | 1.57 (1.4 - 1.93) | <0.001 | 0.23 | <0.001 |

| | | | | | |
|---|---|---|---|---|---|
| IFI30 | 0.78 (0.49 - 0.92) | 1.47 (1.06 - 2.01) | <0.001 | -0.53 | <0.001 |
| IGFBP4 | 0.88 (0.55 - 1.25) | 0.44 (0.15 - 0.65) | <0.001 | 0.58 | <0.001 |
| IGFLR1 | 1.08 (0.86 - 1.37) | 0.92 (0.82 - 1.2) | <0.001 | 0.25 | <0.001 |
| IKZF1 | 2.44 (1.96 - 3.16) | 2.74 (1.9 - 3.3) | <0.001 | -0.39 | 1 |
| IL10RA | 4.31 (3.5 - 5.46) | 3.3 (2.94 - 3.67) | <0.001 | 0.27 | <0.001 |
| IL16 | 1.87 (1.5 - 2.41) | 2.02 (1.5 - 2.54) | 0.45 | -0.27 | 1 |
| IL2RG | 2.89 (2.45 - 3.41) | 2.87 (2.61 - 3.38) | <0.001 | 0.27 | <0.001 |
| IL4R | 4.49 (3.17 - 5.82) | 1.88 (0.92 - 2.88) | <0.001 | 1.18 | <0.001 |
| IMMP2L | 1.29 (0.94 - 1.55) | 0.55 (0.38 - 0.89) | <0.001 | 0.77 | <0.001 |
| INO80D | 1.51 (1.26 - 1.64) | 1.2 (0.99 - 1.49) | <0.001 | 0.2 | <0.001 |
| INPP5F | 2.35 (1.94 - 2.76) | 1.17 (0.99 - 1.35) | <0.001 | 0.53 | <0.001 |
| IQGAP1 | 2.06 (1.76 - 2.53) | 2.12 (1.56 - 2.49) | 0.52 | -0.24 | 1 |
| IQSEC1 | 0.62 (0.16 - 1.43) | 2.15 (1.21 - 2.82) | <0.001 | -1.5 | <0.001 |
| IRAK4 | 0.8 (0.74 - 0.86) | 0.76 (0.63 - 0.89) | <0.001 | 0.24 | <0.001 |
| IRF1 | 1.57 (1.04 - 1.99) | 2.32 (1.79 - 3.15) | 0.94 | -0.2 | 1 |
| IRF4 | 1.04 (0.91 - 1.24) | 1.24 (0.95 - 1.44) | <0.001 | -0.42 | 0.15 |
| IRF9 | 1.06 (0.95 - 1.17) | 0.89 (0.81 - 1.07) | <0.001 | 0.29 | <0.001 |
| ISCU | 3.52 (3.08 - 3.85) | 2.86 (2.69 - 3.09) | <0.001 | 0.49 | <0.001 |
| ITFG2 | 0.7 (0.61 - 0.85) | 0.47 (0.38 - 0.54) | <0.001 | 0.46 | <0.001 |
| ITGA4 | 0.74 (0.17 - 1.47) | 2.17 (1.51 - 3.01) | <0.001 | -1.33 | <0.001 |
| ITGB1 | 1.55 (1.36 - 1.9) | 2 (1.59 - 2.5) | <0.001 | -0.57 | <0.001 |
| ITGB7 | 0.82 (0.69 - 1.14) | 0.99 (0.61 - 1.35) | <0.001 | -0.37 | 1 |
| ITM2B | 2.46 (2.13 - 3.05) | 2.58 (2.17 - 3.33) | <0.001 | 0.26 | <0.001 |
| ITPR2 | 2.16 (1.77 - 2.64) | 1.85 (1.61 - 2.11) | <0.001 | 0.35 | <0.001 |
| JAK2 | 1.27 (1.12 - 1.35) | 0.71 (0.64 - 0.79) | <0.001 | 0.45 | <0.001 |
| JAZF1 | 1.97 (1.67 - 2.26) | 1.53 (1.22 - 1.84) | <0.001 | 0.3 | <0.001 |
| JCHAIN | 1.61 (1.06 - 2.97) | 1.72 (1.22 - 2.33) | <0.001 | 0.66 | <0.001 |
| JMJD1C | 4.35 (4.09 - 4.73) | 3.32 (3.09 - 3.65) | <0.001 | 0.24 | <0.001 |
| JPX | 0.95 (0.83 - 1.05) | 0.73 (0.61 - 0.86) | <0.001 | 0.32 | <0.001 |
| JUN | 4.08 (3.44 - 4.71) | 3.67 (3.27 - 4.45) | <0.001 | 0.32 | <0.001 |
| KANSL1 | 2.1 (1.85 - 2.27) | 1.64 (1.46 - 1.75) | <0.001 | 0.35 | <0.001 |
| KAT6B | 1.74 (1.59 - 1.98) | 1.16 (1.1 - 1.21) | <0.001 | 0.33 | <0.001 |
| KDM5A | 2.38 (2.13 - 2.6) | 2.16 (1.9 - 2.38) | <0.001 | 0.27 | <0.001 |
| KDM6A | 1.23 (1.12 - 1.39) | 0.93 (0.83 - 1.23) | <0.001 | 0.3 | <0.001 |
| KHDRBS1 | 1.73 (1.68 - 1.77) | 1.86 (1.68 - 2.02) | 0.01 | -0.39 | 1 |
| KHDRBS2 | 1.44 (1.28 - 1.91) | 0.56 (0.26 - 0.74) | <0.001 | 0.88 | <0.001 |
| KIF2A | 1.17 (0.95 - 1.62) | 1.57 (1.1 - 1.9) | 0.28 | -0.3 | 1 |
| KLF13 | 1.21 (0.92 - 1.45) | 1.54 (1.19 - 1.76) | <0.001 | -0.58 | <0.001 |
| KLF2 | 4.1 (2.41 - 5.14) | 6.58 (5.06 - 8.35) | <0.001 | -1.73 | <0.001 |

| | | | | | |
|---|---|---|---|---|---|
| KLF3 | 0.57 (0.24 - 1.1) | 1.26 (0.77 - 1.68) | <0.001 | -0.89 | <0.001 |
| KLF6 | 6.7 (5.54 - 9.16) | 7.16 (6.06 - 8.19) | <0.001 | -0.3 | 1 |
| KLHL14 | 0.84 (0.74 - 0.96) | 0.46 (0.36 - 0.59) | <0.001 | 0.39 | <0.001 |
| KMT2A | 2.86 (2.15 - 3.27) | 2.65 (2.26 - 3.23) | 0.95 | -0.28 | 1 |
| KMT2E | 6.29 (5.59 - 7) | 5.28 (5.18 - 5.52) | <0.001 | 0.34 | <0.001 |
| LAPTM5 | 19.15 (16 - 22.35) | 21.52 (19.28 - 25.25) | <0.001 | -0.5 | <0.001 |
| LBH | 3.29 (2.75 - 4.45) | 3.7 (2.77 - 5.18) | 0.65 | -0.21 | 1 |
| LBR | 0.99 (0.68 - 1.25) | 1.58 (1.33 - 1.86) | <0.001 | -0.69 | <0.001 |
| LCP1 | 2.67 (2.19 - 4.15) | 4.28 (2.72 - 5.1) | <0.001 | -0.86 | <0.001 |
| LEF1 | 1.88 (1.59 - 2.33) | 0.57 (0.27 - 0.84) | <0.001 | 0.99 | <0.001 |
| LEMD3 | 1.05 (0.96 - 1.2) | 0.91 (0.78 - 1.11) | <0.001 | 0.25 | <0.001 |
| LINC-PINT | 1.11 (0.8 - 1.35) | 0.75 (0.57 - 1.18) | <0.001 | 0.34 | <0.001 |
| LINC00926 | 5.1 (3.48 - 6.56) | 4.22 (3.07 - 5.8) | <0.001 | 0.8 | <0.001 |
| LINC01480 | 0.84 (0.53 - 1.11) | 0.57 (0.33 - 0.89) | <0.001 | 0.35 | <0.001 |
| LINC01857 | 1.28 (1.02 - 1.8) | 1.05 (0.85 - 1.27) | <0.001 | 0.6 | <0.001 |
| LINC02245 | 0.75 (0.63 - 0.9) | 0.33 (0.22 - 0.41) | <0.001 | 0.61 | <0.001 |
| LINC02397 | 1.22 (0.96 - 1.55) | 0.91 (0.82 - 1.04) | <0.001 | 0.65 | <0.001 |
| LMBRD1 | 2.4 (2.17 - 2.5) | 1.79 (1.67 - 1.92) | <0.001 | 0.4 | <0.001 |
| LONP2 | 1.75 (1.61 - 1.92) | 1.11 (0.91 - 1.26) | <0.001 | 0.35 | <0.001 |
| LPGAT1 | 0.88 (0.72 - 1.03) | 0.98 (0.74 - 1.17) | <0.001 | -0.43 | 0.36 |
| LPP | 2.23 (1.94 - 2.56) | 1.41 (1.23 - 1.55) | <0.001 | 0.61 | <0.001 |
| LRMP | 3.62 (3.1 - 4.29) | 2.62 (2.01 - 3.18) | <0.001 | 0.45 | <0.001 |
| LRRFIP1 | 1.67 (1.41 - 1.99) | 2.57 (1.98 - 3.2) | 0.05 | -0.33 | 1 |
| LSM10 | 0.88 (0.73 - 1.08) | 0.8 (0.65 - 0.92) | <0.001 | 0.39 | <0.001 |
| LSM14A | 2.15 (1.96 - 2.44) | 2.44 (2.07 - 2.83) | 0.06 | -0.33 | 1 |
| LSM7 | 0.89 (0.75 - 1.01) | 1.24 (1.04 - 1.51) | 0.21 | -0.28 | 1 |
| LSP1 | 3.48 (3.1 - 4.03) | 4.9 (4.72 - 5.13) | 0.01 | -0.22 | 1 |
| LTA4H | 1.88 (1.58 - 2.41) | 1.3 (0.98 - 1.44) | <0.001 | 0.56 | <0.001 |
| LTB | 4.3 (3.64 - 5.12) | 6.53 (5.2 - 7.63) | <0.001 | 0.34 | <0.001 |
| LUC7L | 1.8 (1.55 - 2.14) | 1.2 (0.97 - 1.44) | <0.001 | 0.56 | <0.001 |
| LUC7L2 | 2.88 (2.66 - 3.21) | 2.47 (2.24 - 2.82) | <0.001 | 0.21 | <0.001 |
| LUC7L3 | 2.28 (1.61 - 2.6) | 1.69 (1.41 - 2.4) | <0.001 | 0.51 | <0.001 |
| LY6E | 1.17 (0.95 - 1.38) | 1.89 (1.55 - 2.1) | <0.001 | -0.52 | 0.02 |
| LY9 | 1.79 (0.89 - 3.2) | 1.47 (0.57 - 2.57) | <0.001 | 0.35 | <0.001 |
| LYST | 1.97 (1.44 - 2.47) | 1.2 (0.79 - 1.7) | <0.001 | 0.43 | <0.001 |
| MACF1 | 1.09 (0.56 - 1.91) | 1.57 (0.73 - 2.34) | <0.001 | -0.54 | 0.08 |
| MALAT1 | 444.13 (382.59 - 485.82) | 388.78 (336.48 - 454.57) | <0.001 | 0.37 | <0.001 |
| MALT1 | 1.1 (0.99 - 1.23) | 1.27 (1 - 1.37) | <0.001 | -0.42 | <0.001 |

| | | | | | |
|---|---|---|---|---|---|
| MAML2 | 1.09 (0.84 - 1.46) | 0.75 (0.45 - 0.95) | <0.001 | 0.24 | <0.001 |
| MAN2A1 | 1.82 (1.56 - 2.15) | 1.12 (0.92 - 1.29) | <0.001 | 0.35 | <0.001 |
| MAP1LC3B | 1.14 (1.05 - 1.24) | 1.37 (1.24 - 1.5) | 0.94 | -0.23 | 1 |
| MAP3K1 | 3.18 (3 - 3.53) | 2.96 (2.65 - 3.26) | 0.18 | -0.26 | 1 |
| MAP4K1 | 0.69 (0.6 - 0.76) | 1.07 (0.97 - 1.18) | <0.001 | -0.38 | <0.001 |
| MAP4K4 | 0.75 (0.2 - 1.1) | 1.4 (0.95 - 2.02) | <0.001 | -0.76 | <0.001 |
| MAPK1IP1L | 1.47 (1.33 - 1.59) | 1.66 (1.46 - 1.93) | 0.76 | -0.25 | 1 |
| MAPK8IP3 | 2.34 (2.09 - 2.84) | 1.94 (1.63 - 2.67) | <0.001 | 0.53 | <0.001 |
| MARCH1 | 3.11 (2.16 - 3.83) | 2.5 (2.18 - 3.06) | <0.001 | 0.41 | <0.001 |
| MARCKSL1 | 0.53 (0.31 - 0.85) | 0.68 (0.4 - 1.17) | <0.001 | -0.43 | <0.001 |
| MATR3.1 | 1.52 (1.31 - 1.82) | 0.9 (0.7 - 1.14) | <0.001 | 0.47 | <0.001 |
| MBD2 | 0.91 (0.77 - 1.21) | 1.1 (0.87 - 1.48) | <0.001 | -0.51 | <0.001 |
| MBD4 | 1.79 (1.53 - 2.04) | 1.71 (1.65 - 1.75) | <0.001 | 0.24 | <0.001 |
| MBNL1 | 5.57 (4.86 - 5.97) | 4.85 (4.43 - 5.37) | <0.001 | 0.21 | <0.001 |
| MBP | 1.18 (0.83 - 1.47) | 0.95 (0.72 - 1.37) | <0.001 | 0.22 | <0.001 |
| MBTPS1 | 1.23 (1.04 - 1.39) | 1.05 (0.91 - 1.25) | <0.001 | 0.22 | <0.001 |
| MCL1 | 2.41 (2.22 - 2.57) | 3.03 (2.58 - 3.39) | <0.001 | -0.64 | <0.001 |
| MCOLN2 | 1.59 (1.21 - 1.82) | 0.85 (0.66 - 1.16) | <0.001 | 0.55 | <0.001 |
| MCTP2 | 1.5 (1.22 - 2.11) | 0.57 (0.25 - 0.84) | <0.001 | 0.8 | <0.001 |
| MDFIC | 1.08 (0.93 - 1.38) | 1.09 (1.01 - 1.25) | <0.001 | -0.34 | 1 |
| MEF2A | 1.35 (1.14 - 1.74) | 1.74 (1.24 - 2.09) | <0.001 | -0.44 | 0.01 |
| MEF2C | 3.15 (2.65 - 4.03) | 3.77 (3.02 - 4.48) | 0.02 | -0.32 | 1 |
| METTL8 | 1.48 (0.95 - 2) | 0.78 (0.52 - 1.12) | <0.001 | 1.08 | <0.001 |
| MKNK2 | 1.85 (1.45 - 2.36) | 3.09 (2.33 - 4.3) | <0.001 | -0.98 | <0.001 |
| MOB1A | 3.26 (2.76 - 4.11) | 4.14 (3.06 - 4.85) | <0.001 | -0.54 | <0.001 |
| MOB3A | 0.72 (0.42 - 1.07) | 0.97 (0.61 - 1.37) | <0.001 | -0.37 | 0.13 |
| MON2 | 1.73 (1.51 - 1.95) | 1.2 (1.07 - 1.42) | <0.001 | 0.49 | <0.001 |
| MORC3 | 1.9 (1.54 - 2.26) | 1.21 (1.07 - 1.57) | <0.001 | 0.29 | <0.001 |
| MPHOSPH8 | 2.04 (1.62 - 2.3) | 1.65 (1.42 - 2.14) | <0.001 | 0.38 | <0.001 |
| MRNIP | 1.15 (0.95 - 1.5) | 0.63 (0.51 - 0.83) | <0.001 | 0.51 | <0.001 |
| MS4A1 | 5.96 (3.46 - 8.31) | 14.24 (11.62 - 20.2) | <0.001 | -1.97 | <0.001 |
| MSI2 | 3.25 (2.66 - 3.88) | 1.47 (1.04 - 2.05) | <0.001 | 1.14 | <0.001 |
| MSN | 2.53 (2.03 - 3.44) | 3.09 (2.08 - 3.84) | <0.001 | -0.55 | <0.001 |
| MT-CO1 | 109.67 (102.65 - 119.13) | 107.94 (95.67 - 121.05) | <0.001 | -0.34 | <0.001 |
| MT-CO2 | 72.02 (66.5 - 77.09) | 71.13 (66.58 - 74.92) | <0.001 | 0.31 | <0.001 |
| MT-CO3 | 73.77 (66.33 - 79.88) | 66.32 (61.59 - 68.82) | <0.001 | 0.36 | <0.001 |
| MT-CYB | 72.4 (64.53 - 78.15) | 64.49 (60.32 - 67.34) | <0.001 | 0.38 | <0.001 |
| MT-ND1 | 26.06 (23.84 - 29.06) | 22.67 (21.56 - | <0.001 | 0.24 | <0.001 |

| | | | | | |
|---|---|---|---|---|---|
| | | 23.71) | | | |
| MT-ND2 | 35.63 (27.2 - 54.92) | 25.68 (22.22 - 27.6) | 0.69 | 0.42 | 1 |
| MT-ND4 | 19.51 (17.54 - 22.53) | 20.18 (17.16 - 22.85) | <0.001 | -0.28 | <0.001 |
| MT-ND4L | 36.86 (32.94 - 42.49) | 40.93 (34.6 - 44.86) | <0.001 | -0.4 | <0.001 |
| MT-ND5 | 26.35 (23.68 - 28.96) | 26.34 (23.38 - 32.29) | <0.001 | -0.69 | <0.001 |
| MTERF4 | 0.8 (0.56 - 0.92) | 0.67 (0.51 - 0.9) | <0.001 | 0.23 | <0.001 |
| MTPN | 3.95 (3.37 - 4.77) | 4.71 (3.87 - 5.35) | <0.001 | -0.44 | 0.03 |
| MTRNR2L12 | 32.37 (27.26 - 39.7) | 33.19 (27.76 - 37.74) | <0.001 | -0.26 | <0.001 |
| MTRNR2L8 | 42.5 (36.98 - 58.79) | 43.78 (37.05 - 55.06) | <0.001 | 2.9 | <0.001 |
| MYH9 | 2.97 (2.44 - 3.72) | 3.33 (2.4 - 3.98) | <0.001 | -0.49 | <0.001 |
| MYL6 | 4.75 (4.4 - 5.11) | 4.64 (3.88 - 5.59) | <0.001 | 0.31 | <0.001 |
| MYO1E | 1.24 (1.04 - 1.73) | 0.58 (0.29 - 0.74) | <0.001 | 0.73 | <0.001 |
| MYO9B | 1.53 (1.28 - 1.93) | 1.46 (1.12 - 1.82) | 0.02 | -0.29 | 1 |
| MZB1 | 0.9 (0.38 - 1.76) | 0.76 (0.44 - 1.27) | <0.001 | 0.26 | <0.001 |
| N4BP2L2 | 2.89 (2.53 - 3.06) | 2.07 (1.88 - 2.46) | <0.001 | 0.46 | <0.001 |
| NAA50 | 1.35 (1.19 - 1.76) | 1.73 (1.19 - 1.9) | <0.001 | -0.45 | <0.001 |
| NAGK | 1.05 (0.99 - 1.2) | 0.7 (0.61 - 0.85) | <0.001 | 0.34 | <0.001 |
| NAP1L1 | 2.76 (2.37 - 3.16) | 3.2 (2.88 - 3.57) | 0.13 | -0.28 | 1 |
| NAP1L4 | 1.21 (1.17 - 1.26) | 0.85 (0.76 - 0.9) | <0.001 | 0.21 | <0.001 |
| NCF1 | 3.79 (3.17 - 4.5) | 3.41 (2.92 - 4.45) | <0.001 | 0.23 | <0.001 |
| NCKAP1L | 2.01 (1.77 - 2.3) | 1.55 (1.38 - 1.74) | <0.001 | 0.37 | <0.001 |
| NCL | 5.83 (5.36 - 6.73) | 6.65 (5.57 - 7.08) | 0.04 | -0.29 | 1 |
| NCOA3 | 4.52 (4.32 - 4.71) | 4.08 (3.72 - 4.54) | 0.84 | -0.21 | 1 |
| NCOR1 | 2.77 (2.38 - 3.12) | 2.33 (2.03 - 2.62) | <0.001 | 0.25 | <0.001 |
| NDFIP1 | 1.31 (1.13 - 1.49) | 0.82 (0.71 - 0.92) | <0.001 | 0.24 | <0.001 |
| NDUFV1 | 0.91 (0.8 - 1.05) | 0.71 (0.61 - 0.9) | <0.001 | 0.35 | <0.001 |
| NDUFV2 | 2.19 (2.07 - 2.31) | 1.73 (1.6 - 1.87) | <0.001 | 0.32 | <0.001 |
| NEMF | 1.91 (1.58 - 2.26) | 1.61 (1.38 - 2.1) | <0.001 | 0.4 | <0.001 |
| NFAT5 | 2.41 (1.63 - 2.76) | 1.79 (1.47 - 2.76) | <0.001 | 0.43 | <0.001 |
| NFKB1 | 1.59 (1.21 - 1.8) | 2.06 (1.65 - 3.23) | <0.001 | -0.41 | 0.13 |
| NFKBID | 1.59 (1.38 - 1.89) | 1.02 (0.75 - 1.27) | <0.001 | 0.61 | <0.001 |
| NFX1 | 1.13 (0.94 - 1.23) | 0.77 (0.66 - 0.92) | <0.001 | 0.22 | <0.001 |
| NGLY1 | 1.8 (1.61 - 1.93) | 1.57 (1.39 - 1.78) | <0.001 | 0.21 | <0.001 |
| NIBAN3 | 3.57 (2.99 - 4.27) | 3.04 (2.57 - 3.7) | <0.001 | 0.37 | <0.001 |
| NIN | 1.45 (1.05 - 1.86) | 2.16 (1.59 - 2.74) | <0.001 | -0.68 | <0.001 |
| NKTR | 3.1 (2.61 - 3.46) | 2.26 (1.96 - 3.02) | <0.001 | 0.4 | <0.001 |
| NME2 | 1.81 (1.63 - 2.05) | 2.59 (2.31 - 2.78) | 0.01 | -0.34 | 1 |
| NOP53 | 3.71 (3.16 - 4.41) | 3.69 (3.22 - 4.22) | 0.32 | -0.24 | 1 |

| | | | | | |
|---|---|---|---|---|---|
| NOP58 | 3.41 (2.81 - 4.1) | 2.55 (2.38 - 2.63) | <0.001 | 0.36 | <0.001 |
| NORAD | 2.33 (2.12 - 2.61) | 2.24 (1.88 - 2.58) | 0.3 | -0.25 | 1 |
| NOSIP | 2.33 (1.79 - 3.1) | 1.58 (1.35 - 1.82) | <0.001 | 1.42 | <0.001 |
| NPEPPS | 2.78 (2.21 - 3.05) | 2.15 (1.78 - 2.72) | <0.001 | 0.37 | <0.001 |
| NPM1 | 5.95 (5.52 - 6.58) | 7.11 (6.47 - 7.93) | <0.001 | -0.32 | 0.68 |
| NSF | 2.56 (2.16 - 3.01) | 1.93 (1.52 - 2.41) | <0.001 | 0.32 | <0.001 |
| NSMCE1 | 0.72 (0.59 - 0.83) | 0.51 (0.42 - 0.56) | <0.001 | 0.56 | <0.001 |
| NT5C3A | 0.79 (0.68 - 0.94) | 0.61 (0.49 - 0.76) | <0.001 | 0.33 | <0.001 |
| NUDT4 | 1.3 (1.03 - 1.7) | 1.86 (1.41 - 2.28) | <0.001 | -0.67 | <0.001 |
| NUP214 | 1.38 (1.13 - 1.65) | 0.97 (0.79 - 1.21) | <0.001 | 0.33 | <0.001 |
| ODC1 | 1.62 (1.34 - 2.06) | 1.7 (1.39 - 2.19) | <0.001 | -0.47 | <0.001 |
| OFD1 | 1.21 (0.8 - 1.35) | 1.09 (0.94 - 1.57) | <0.001 | 0.28 | <0.001 |
| OGA | 4.03 (3.33 - 4.39) | 3.28 (2.86 - 4.34) | <0.001 | 0.36 | <0.001 |
| OGT | 3.08 (2.67 - 3.81) | 2.16 (1.83 - 2.35) | <0.001 | 0.26 | <0.001 |
| OIP5-AS1 | 1.35 (1.21 - 1.48) | 1.37 (1.23 - 1.53) | 0.62 | -0.22 | 1 |
| ORAI2 | 1.51 (1.11 - 1.74) | 1.26 (1.11 - 1.43) | <0.001 | 0.27 | <0.001 |
| OSBPL10 | 0.89 (0.72 - 1.1) | 0.95 (0.76 - 1.14) | 0.01 | -0.3 | 1 |
| OSBPL8 | 1.23 (1.01 - 1.49) | 1.4 (1.1 - 1.66) | 0.25 | -0.28 | 1 |
| P2RX5 | 1.08 (0.49 - 1.72) | 3.42 (2.61 - 4.53) | <0.001 | -1.8 | <0.001 |
| PABPC1 | 12.55 (11.34 - 15.64) | 16.02 (12.32 - 18.51) | <0.001 | -0.54 | <0.001 |
| PABPC4 | 0.84 (0.72 - 1.03) | 1.13 (0.84 - 1.52) | <0.001 | -0.54 | <0.001 |
| PABPN1 | 1.28 (1.08 - 1.41) | 1.17 (1.07 - 1.41) | 0.3 | -0.23 | 1 |
| PALM2-AKAP2 | 1.66 (1.03 - 2.16) | 1.91 (1.34 - 2.62) | 0.01 | -0.45 | 1 |
| PATL2 | 0.75 (0.65 - 0.91) | 0.42 (0.35 - 0.5) | <0.001 | 0.58 | <0.001 |
| PCBP1 | 1.76 (1.46 - 2.08) | 2.22 (1.77 - 2.68) | <0.001 | -0.66 | <0.001 |
| PCBP2 | 2.55 (2.34 - 2.77) | 2.48 (2.24 - 2.98) | <0.001 | 0.32 | <0.001 |
| PCDH9 | 3.62 (2.16 - 5.31) | 1.95 (1.06 - 2.76) | <0.001 | 0.98 | <0.001 |
| PCM1 | 3.49 (3.2 - 3.95) | 2.96 (2.64 - 3.3) | <0.001 | 0.25 | <0.001 |
| PDCD10 | 1 (0.79 - 1.12) | 0.91 (0.78 - 1.13) | <0.001 | 0.24 | <0.001 |
| PDE4B | 3.67 (3.06 - 4.7) | 3.18 (2.65 - 4.2) | <0.001 | 0.5 | <0.001 |
| PDE4DIP | 0.8 (0.59 - 1.31) | 0.53 (0.29 - 0.61) | <0.001 | 0.72 | <0.001 |
| PDIA6 | 1.11 (0.87 - 1.33) | 1.25 (0.98 - 1.46) | 0.86 | -0.29 | 1 |
| PDLIM1 | 1.18 (1.04 - 1.37) | 1.54 (1.47 - 1.69) | 0.66 | -0.25 | 1 |
| PDS5A | 1.49 (1.37 - 1.64) | 1.44 (1.32 - 1.59) | 0.32 | -0.23 | 1 |
| PDXDC1 | 1.24 (1.1 - 1.4) | 0.96 (0.87 - 1.09) | <0.001 | 0.31 | <0.001 |
| PEBP1 | 3.94 (3.44 - 4.51) | 3.63 (3.42 - 4.09) | <0.001 | 0.33 | <0.001 |
| PFDN5 | 8.73 (8.24 - 9.72) | 8.44 (7.86 - 9.52) | <0.001 | 0.5 | <0.001 |
| PFN1 | 5.02 (4.35 - 5.86) | 8.01 (7.12 - 8.49) | <0.001 | -0.51 | <0.001 |
| PHC3 | 2.74 (2.44 - 2.94) | 1.89 (1.64 - 2.08) | <0.001 | 0.21 | <0.001 |

| | | | | | |
|---|---|---|---|---|---|
| PHF1 | 1.29 (1.19 - 1.44) | 1.06 (0.93 - 1.19) | <0.001 | 0.35 | <0.001 |
| PHF20L1 | 1.08 (0.84 - 1.19) | 0.81 (0.7 - 1.04) | <0.001 | 0.24 | <0.001 |
| PHF3 | 3.35 (3.14 - 3.59) | 2.52 (2.36 - 2.72) | <0.001 | 0.26 | <0.001 |
| PIGR | 1.05 (0.67 - 1.44) | 0.7 (0.42 - 1.26) | <0.001 | 0.5 | <0.001 |
| PIK3AP1 | 1.31 (1.08 - 1.71) | 1.53 (1.03 - 2.1) | <0.001 | -0.56 | <0.001 |
| PIK3IP1 | 1.36 (1.11 - 1.61) | 0.89 (0.75 - 1.08) | <0.001 | 0.36 | <0.001 |
| PIKFYVE | 1.26 (1.07 - 1.59) | 1.55 (1.25 - 1.86) | <0.001 | -0.42 | 0.03 |
| PIP4K2A | 0.64 (0.4 - 0.92) | 1.29 (0.88 - 1.75) | <0.001 | -1.01 | <0.001 |
| PKM | 1.27 (1.01 - 1.8) | 1.8 (1.27 - 2.18) | <0.001 | -0.43 | 0.53 |
| PKN2 | 1.69 (1.5 - 1.92) | 1.22 (1.03 - 1.5) | <0.001 | 0.55 | <0.001 |
| PLAC8 | 1.33 (0.27 - 2.11) | 2.2 (1.45 - 3.69) | <0.001 | -0.48 | <0.001 |
| PLCL2 | 2.38 (1.85 - 2.82) | 1.3 (1 - 1.66) | <0.001 | 0.32 | <0.001 |
| PLEK | 0.71 (0.49 - 0.94) | 1.89 (1.59 - 2.27) | <0.001 | -1.29 | <0.001 |
| PLEKHF2 | 1.64 (1.4 - 2.09) | 2.06 (1.52 - 2.4) | <0.001 | -0.48 | 0.01 |
| PLEKHG1 | 2.7 (2.05 - 3.09) | 1.75 (1.45 - 2.2) | <0.001 | 0.46 | <0.001 |
| PLEKHO1 | 0.93 (0.79 - 1.12) | 1.2 (0.92 - 1.57) | <0.001 | -0.45 | <0.001 |
| PLPP5 | 1.91 (1.13 - 2.43) | 1.32 (0.94 - 1.86) | <0.001 | 0.57 | <0.001 |
| PMAIP1 | 3.04 (2.35 - 3.83) | 2.05 (1.59 - 2.59) | <0.001 | 0.42 | <0.001 |
| PNISR | 4.09 (3.68 - 4.5) | 3.89 (3.33 - 4.53) | <0.001 | 0.42 | <0.001 |
| PNN | 3.21 (2.84 - 3.43) | 2.4 (2.18 - 2.83) | <0.001 | 0.42 | <0.001 |
| PNRC1 | 11.87 (11.01 - 13.49) | 12.82 (10.61 - 14.85) | <0.001 | -0.25 | <0.001 |
| POLR2J3 | 1.05 (0.74 - 1.31) | 0.75 (0.5 - 1.08) | <0.001 | 0.46 | <0.001 |
| POU2AF1 | 1.34 (1.03 - 2) | 1.81 (1.08 - 2.31) | <0.001 | -0.58 | <0.001 |
| POU2F2 | 1.91 (1.64 - 2.18) | 2.14 (1.75 - 2.69) | 0.73 | -0.22 | 1 |
| PPHLN1 | 1.25 (1.15 - 1.4) | 1.17 (1.01 - 1.28) | <0.001 | 0.24 | <0.001 |
| PPIA | 8.06 (7.45 - 8.64) | 9.39 (8.79 - 10.04) | <0.001 | -0.22 | 1 |
| PPP1CB | 0.99 (0.88 - 1.27) | 1.15 (0.8 - 1.31) | 0.03 | -0.28 | 1 |
| PPP1R10 | 1.83 (1.39 - 2.3) | 1.52 (1.22 - 1.92) | <0.001 | 0.25 | <0.001 |
| PPP1R12A | 2.36 (2.16 - 2.65) | 1.82 (1.6 - 2.01) | <0.001 | 0.44 | <0.001 |
| PPP1R16B | 0.98 (0.79 - 1.11) | 1.23 (1.02 - 1.35) | <0.001 | -0.35 | 1 |
| PPP1R18 | 0.59 (0.3 - 0.91) | 1.24 (0.92 - 1.83) | <0.001 | -1.01 | <0.001 |
| PPP1R2 | 2.65 (2.22 - 2.9) | 2.38 (2.01 - 3.01) | <0.001 | 0.26 | <0.001 |
| PPP2R5C | 1.65 (1.54 - 1.7) | 1.43 (1.26 - 1.57) | <0.001 | 0.2 | <0.001 |
| PPP3CA | 1.35 (0.94 - 1.71) | 1.86 (1.4 - 2.39) | <0.001 | -0.51 | <0.001 |
| PPP3R1 | 1.17 (1.01 - 1.31) | 1.16 (0.97 - 1.4) | <0.001 | -0.37 | 1 |
| PPP4R3B | 1.94 (1.41 - 2.15) | 1.58 (1.29 - 2.24) | <0.001 | 0.2 | <0.001 |
| PRDX1 | 0.88 (0.8 - 1.05) | 1.09 (0.94 - 1.14) | 0.92 | -0.22 | 1 |
| PRKAR1A | 0.85 (0.7 - 1.02) | 1.09 (0.86 - 1.23) | <0.001 | -0.39 | 1 |
| PRKCE | 2.83 (2.25 - 3.71) | 1.31 (0.9 - 1.71) | <0.001 | 0.73 | <0.001 |

| | | | | | |
|---|---|---|---|---|---|
| PRKD3 | 0.96 (0.84 - 1.16) | 1.23 (0.92 - 1.44) | <0.001 | -0.56 | <0.001 |
| PRMT2 | 1.19 (1.01 - 1.33) | 0.9 (0.78 - 1.09) | <0.001 | 0.41 | <0.001 |
| PRRC2A | 1.02 (0.9 - 1.17) | 1.1 (0.86 - 1.28) | 0.09 | -0.27 | 1 |
| PSAP | 1.91 (1.36 - 2.69) | 3.08 (2.19 - 3.83) | <0.001 | -0.74 | <0.001 |
| PSMA3-AS1 | 0.99 (0.81 - 1.09) | 0.88 (0.69 - 1.17) | <0.001 | 0.22 | <0.001 |
| PSMB9 | 1.03 (0.85 - 1.4) | 1.4 (0.93 - 1.77) | 0.06 | -0.34 | 1 |
| PTBP3 | 2.23 (1.82 - 2.76) | 2.31 (1.62 - 2.89) | 0.37 | -0.29 | 1 |
| PTEN | 1.56 (1.28 - 1.75) | 1.53 (1.33 - 1.88) | 0.13 | -0.29 | 1 |
| PTMA | 23.25 (21.45 - 24.81) | 33.04 (29.89 - 38.33) | <0.001 | -0.39 | <0.001 |
| PTP4A2 | 1.36 (1 - 1.8) | 2.27 (1.66 - 2.87) | <0.001 | -0.89 | <0.001 |
| PTPN1 | 4.14 (3.54 - 4.67) | 4 (3.67 - 4.57) | 0.98 | -0.24 | 1 |
| PTPN22 | 1.27 (1.06 - 1.58) | 0.89 (0.75 - 1.03) | <0.001 | 0.33 | <0.001 |
| PTPRC | 9.68 (8.6 - 11.52) | 10.47 (8.13 - 12.65) | <0.001 | -0.5 | <0.001 |
| PXK | 0.74 (0.53 - 0.95) | 0.98 (0.66 - 1.41) | <0.001 | -0.61 | <0.001 |
| R3HDM4 | 0.93 (0.83 - 1.18) | 1.13 (0.85 - 1.31) | <0.001 | -0.39 | 1 |
| RAB14 | 1.41 (1.27 - 1.74) | 1.57 (1.22 - 1.8) | 0.44 | -0.25 | 1 |
| RAB31 | 0.83 (0.73 - 0.94) | 0.62 (0.54 - 0.75) | <0.001 | 0.23 | <0.001 |
| RAB3GAP1 | 1.57 (1.42 - 1.66) | 0.97 (0.86 - 1.19) | <0.001 | 0.49 | <0.001 |
| RAB8B | 1.21 (1.06 - 1.47) | 1.42 (1.05 - 1.74) | 0.16 | -0.31 | 1 |
| RABEP2 | 1 (0.89 - 1.07) | 0.88 (0.78 - 1.01) | <0.001 | 0.2 | <0.001 |
| RABGAP1L | 2.39 (1.71 - 2.68) | 1.88 (1.45 - 2.5) | <0.001 | 0.26 | <0.001 |
| RAC2 | 3.41 (3.03 - 4.22) | 4.8 (3.93 - 5.45) | <0.001 | -0.29 | 1 |
| RAD23B | 1.24 (0.93 - 1.54) | 1.51 (1.07 - 1.93) | <0.001 | -0.41 | 1 |
| RALGAPA1 | 1.08 (0.86 - 1.22) | 0.63 (0.51 - 0.84) | <0.001 | 0.36 | <0.001 |
| RAN | 2.66 (2.19 - 2.98) | 2.41 (1.99 - 3.03) | <0.001 | 0.21 | <0.001 |
| RAPGEF1 | 1.6 (1.35 - 2.13) | 1.77 (1.51 - 2.03) | <0.001 | -0.42 | <0.001 |
| RASA1 | 1.28 (1.02 - 1.57) | 0.63 (0.44 - 0.75) | <0.001 | 0.35 | <0.001 |
| RASGRF1 | 0.99 (0.91 - 1.24) | 0.41 (0.25 - 0.48) | <0.001 | 0.6 | <0.001 |
| RASSF5 | 0.9 (0.7 - 1.31) | 1.21 (0.73 - 1.41) | <0.001 | -0.34 | 1 |
| RB1CC1 | 1.64 (1.4 - 1.87) | 1.14 (1.06 - 1.21) | <0.001 | 0.2 | <0.001 |
| RBM25 | 4.02 (3.51 - 4.62) | 3.6 (3.21 - 4.17) | <0.001 | 0.33 | <0.001 |
| RBM3 | 3.41 (2.96 - 3.78) | 4.22 (3.71 - 4.68) | <0.001 | -0.35 | 1 |
| RBM39 | 6.28 (5.61 - 6.99) | 5.49 (4.85 - 6.3) | <0.001 | 0.29 | <0.001 |
| RBM6 | 1.4 (1.27 - 1.54) | 1.01 (0.88 - 1.13) | <0.001 | 0.35 | <0.001 |
| RECQL | 0.78 (0.65 - 0.97) | 0.56 (0.42 - 0.74) | <0.001 | 0.23 | <0.001 |
| RESF1 | 7.35 (6.59 - 8.01) | 5.83 (4.96 - 6.88) | <0.001 | 0.22 | <0.001 |
| RHOA | 6.26 (5.78 - 7.41) | 7.75 (6.36 - 8.81) | <0.001 | -0.48 | <0.001 |
| RHOH | 5.85 (5.19 - 6.88) | 3.67 (3.29 - 4.11) | <0.001 | 0.64 | <0.001 |
| RIC1 | 2.18 (1.79 - 3.05) | 1.47 (1.15 - 1.76) | <0.001 | 0.44 | <0.001 |

| | | | | | |
|---|---|---|---|---|---|
| RIC3 | 0.97 (0.89 - 1.1) | 0.56 (0.47 - 0.64) | <0.001 | 0.63 | <0.001 |
| RIPOR2 | 4.54 (3.71 - 5.87) | 4.25 (3.46 - 5.21) | <0.001 | 0.22 | <0.001 |
| RLF | 1.29 (1.02 - 1.55) | 0.91 (0.76 - 1.21) | <0.001 | 0.34 | <0.001 |
| RNASE6 | 0.57 (0.49 - 0.75) | 0.51 (0.42 - 0.7) | <0.001 | 0.81 | <0.001 |
| RNASET2 | 2.37 (2.24 - 2.47) | 3.29 (2.93 - 3.54) | 0.28 | -0.3 | 1 |
| RNF13 | 0.79 (0.69 - 0.86) | 0.59 (0.51 - 0.68) | <0.001 | 0.28 | <0.001 |
| RNF130 | 0.89 (0.77 - 1.07) | 0.53 (0.48 - 0.58) | <0.001 | 0.34 | <0.001 |
| RNF141 | 0.73 (0.56 - 0.93) | 0.99 (0.72 - 1.38) | <0.001 | -0.49 | <0.001 |
| RNF19A | 3.8 (2.95 - 6.42) | 2.69 (2.25 - 2.98) | <0.001 | 0.25 | <0.001 |
| RNF213 | 1.91 (1.52 - 2.17) | 2.63 (2.25 - 3.24) | <0.001 | -0.38 | 1 |
| RNF216 | 1.29 (1.11 - 1.46) | 0.84 (0.7 - 0.93) | <0.001 | 0.41 | <0.001 |
| RNF41 | 1.7 (1.38 - 2.07) | 0.92 (0.55 - 1.15) | <0.001 | 0.41 | <0.001 |
| RNH1 | 1 (0.75 - 1.23) | 0.94 (0.81 - 1.24) | <0.001 | 0.22 | <0.001 |
| RNMT | 2.07 (1.78 - 2.6) | 1.61 (1.29 - 2.13) | <0.001 | 0.4 | <0.001 |
| RNPC3 | 1.65 (1.47 - 1.84) | 1.02 (0.88 - 1.24) | <0.001 | 0.59 | <0.001 |
| ROR1 | 2.29 (1.53 - 2.82) | 0.64 (0.22 - 1.28) | <0.001 | 1.13 | <0.001 |
| RRAS2 | 1.47 (1.22 - 1.66) | 0.94 (0.84 - 1.07) | <0.001 | 0.2 | <0.001 |
| RRBP1 | 1.57 (1.17 - 2.28) | 0.74 (0.15 - 1.13) | <0.001 | 0.62 | <0.001 |
| RUBCNL | 3.9 (3.13 - 4.85) | 2.23 (1.64 - 2.62) | <0.001 | 0.75 | <0.001 |
| S100A10 | 0.61 (0.32 - 0.82) | 1.28 (0.96 - 1.84) | <0.001 | -0.97 | <0.001 |
| S100A4 | 2.04 (1.69 - 2.77) | 2.46 (1.72 - 3.19) | <0.001 | 0.73 | <0.001 |
| S100A6 | 2.37 (1.85 - 2.7) | 2.39 (2.06 - 2.81) | <0.001 | 0.37 | <0.001 |
| S1PR1 | 0.55 (0.29 - 0.9) | 0.79 (0.4 - 1.15) | <0.001 | -0.65 | <0.001 |
| SAFB2 | 1.5 (1.3 - 1.7) | 0.97 (0.87 - 1.08) | <0.001 | 0.36 | <0.001 |
| SARAF | 4.68 (4.13 - 5.83) | 4.32 (3.53 - 5.16) | <0.001 | 0.49 | <0.001 |
| SCAF11 | 3.8 (3.36 - 4.35) | 2.99 (2.47 - 3.57) | <0.001 | 0.31 | <0.001 |
| SEC14L1 | 1.54 (1.23 - 1.83) | 0.92 (0.78 - 1.12) | <0.001 | 0.41 | <0.001 |
| SECISBP2 | 1.45 (1.34 - 1.58) | 1.03 (0.96 - 1.09) | <0.001 | 0.31 | <0.001 |
| SECISBP2L | 2.92 (2.4 - 3.47) | 2.21 (1.88 - 2.72) | <0.001 | 0.34 | <0.001 |
| SEL1L3 | 3.57 (2.44 - 4.64) | 3.1 (2.45 - 3.9) | 0.99 | -0.2 | 1 |
| SEMA4B | 0.71 (0.54 - 0.91) | 1.05 (0.77 - 1.42) | <0.001 | -0.64 | <0.001 |
| SENP6 | 2.18 (1.87 - 2.39) | 1.87 (1.63 - 2.19) | <0.001 | 0.26 | <0.001 |
| SEPTIN2 | 2.14 (1.87 - 2.34) | 1.84 (1.54 - 2.26) | <0.001 | 0.22 | <0.001 |
| SERBP1 | 2.2 (2.05 - 2.33) | 2.6 (2.52 - 2.72) | 0.41 | -0.22 | 1 |
| SERF2 | 6.03 (5.22 - 7.19) | 5.66 (4.94 - 6.89) | <0.001 | 0.4 | <0.001 |
| SERINC5 | 1.55 (1.26 - 2.34) | 0.73 (0.61 - 0.91) | <0.001 | 0.61 | <0.001 |
| SESN1 | 1.12 (0.78 - 1.57) | 0.52 (0.26 - 0.82) | <0.001 | 0.65 | <0.001 |
| SESN3 | 5.35 (4.07 - 6.84) | 3.72 (3.13 - 4.34) | <0.001 | 0.45 | <0.001 |
| SESTD1 | 1.2 (0.65 - 1.47) | 0.65 (0.43 - 0.99) | <0.001 | 0.43 | <0.001 |

| | | | | | |
|---|---|---|---|---|---|
| SET | 2.8 (2.15 - 3.42) | 3.37 (2.55 - 4.12) | <0.001 | -0.7 | <0.001 |
| SF1 | 6.04 (5.36 - 6.46) | 5.48 (4.82 - 6.7) | <0.001 | 0.31 | <0.001 |
| SF3A1 | 1.16 (1.01 - 1.39) | 1.15 (0.84 - 1.44) | 0.16 | -0.26 | 1 |
| SF3B1 | 5.49 (5.1 - 6.06) | 4.92 (4.4 - 5.65) | <0.001 | 0.21 | <0.001 |
| SFMBT1 | 2.3 (1.89 - 2.75) | 0.82 (0.55 - 1.02) | <0.001 | 0.95 | <0.001 |
| SFSWAP | 2.02 (1.72 - 2.57) | 1.29 (1.07 - 1.5) | <0.001 | 0.69 | <0.001 |
| SGPP1 | 1.98 (1.52 - 2.65) | 1.21 (0.89 - 1.49) | <0.001 | 0.23 | <0.001 |
| SH3BGRL3 | 3.88 (3.7 - 4.22) | 5.21 (4.83 - 5.59) | <0.001 | -0.37 | 0.01 |
| SH3BP5 | 3.48 (2.91 - 3.86) | 2.78 (2.41 - 3.36) | <0.001 | 0.28 | <0.001 |
| SH3KBP1 | 0.81 (0.66 - 0.94) | 1.23 (1.07 - 1.47) | <0.001 | -0.44 | <0.001 |
| SHMT2 | 1.15 (0.98 - 1.33) | 1.13 (1.02 - 1.35) | <0.001 | 0.48 | <0.001 |
| SHOC2 | 1.24 (1.12 - 1.41) | 1.42 (1.23 - 1.54) | 0.12 | -0.3 | 1 |
| SHPRH | 1.01 (0.93 - 1.09) | 0.65 (0.59 - 0.71) | <0.001 | 0.26 | <0.001 |
| SKAP1 | 1.89 (1.52 - 2.61) | 0.82 (0.37 - 1.16) | <0.001 | 0.72 | <0.001 |
| SKP1 | 3.8 (3.48 - 4.2) | 3.16 (2.91 - 3.58) | <0.001 | 0.39 | <0.001 |
| SLC23A2 | 1.12 (1.02 - 1.33) | 0.52 (0.37 - 0.59) | <0.001 | 0.57 | <0.001 |
| SLC25A5 | 1.33 (1.26 - 1.46) | 1.96 (1.77 - 2.22) | 0.95 | -0.25 | 1 |
| SLC38A1 | 5.66 (4.89 - 6.86) | 4.73 (3.65 - 5.55) | <0.001 | 0.36 | <0.001 |
| SLC38A2 | 2.06 (1.78 - 2.32) | 1.63 (1.35 - 1.91) | <0.001 | 0.2 | <0.001 |
| SLC44A2 | 0.97 (0.83 - 1.25) | 1.15 (0.8 - 1.42) | <0.001 | -0.37 | 1 |
| SLC9A7 | 2.04 (1.59 - 2.31) | 1.25 (1.06 - 1.56) | <0.001 | 0.21 | <0.001 |
| SMAD4 | 1.42 (1.16 - 1.58) | 0.87 (0.78 - 1.03) | <0.001 | 0.24 | <0.001 |
| SMAP2 | 9.85 (8.15 - 10.9) | 9.93 (8.46 - 11.26) | <0.001 | -0.29 | <0.001 |
| SMARCA5 | 1.88 (1.69 - 2) | 1.46 (1.37 - 1.68) | <0.001 | 0.23 | <0.001 |
| SMARCB1 | 1.76 (1.6 - 2.06) | 1.54 (1.37 - 1.75) | <0.001 | 0.29 | <0.001 |
| SMARCC1 | 1.32 (1.17 - 1.42) | 1.33 (1.08 - 1.6) | <0.001 | -0.39 | 1 |
| SMARCC2 | 1.7 (1.52 - 1.94) | 1.25 (1.04 - 1.43) | <0.001 | 0.36 | <0.001 |
| SMCHD1 | 14.52 (12.92 - 15.73) | 11.63 (10.9 - 13.09) | <0.001 | 0.5 | <0.001 |
| SMIM14 | 2.09 (1.68 - 2.55) | 2.01 (1.7 - 2.56) | 0.19 | -0.21 | 1 |
| SND1 | 1.12 (0.95 - 1.3) | 1.13 (1.01 - 1.31) | 0.24 | -0.23 | 1 |
| SNHG14 | 1.54 (1.28 - 1.73) | 0.88 (0.71 - 1.06) | <0.001 | 0.47 | <0.001 |
| SNHG29 | 2.75 (2.33 - 3.59) | 3.47 (3.02 - 4.04) | 0.44 | -0.26 | 1 |
| SNHG7 | 1.03 (0.86 - 1.18) | 0.98 (0.86 - 1.17) | <0.001 | -0.37 | 0.25 |
| SNRNP40 | 0.9 (0.83 - 1.01) | 0.67 (0.58 - 0.74) | <0.001 | 0.24 | <0.001 |
| SNRNP70 | 2.15 (1.81 - 2.41) | 1.43 (1.19 - 1.75) | <0.001 | 0.35 | <0.001 |
| SNX18 | 1.37 (1.22 - 1.64) | 0.75 (0.61 - 0.89) | <0.001 | 0.42 | <0.001 |
| SNX29 | 2.43 (1.86 - 2.96) | 1.5 (1.09 - 2.05) | <0.001 | 0.89 | <0.001 |
| SNX3 | 2.12 (1.93 - 2.5) | 2.81 (2.19 - 3.14) | <0.001 | -0.63 | <0.001 |
| SNX9 | 5.19 (3.63 - 7.29) | 3.15 (2.19 - 3.72) | <0.001 | 0.6 | <0.001 |

| | | | | | |
|---|---|---|---|---|---|
| SOD1 | 1.21 (0.95 - 1.55) | 1.2 (1.07 - 1.45) | <0.001 | 0.3 | <0.001 |
| SP100 | 5.71 (5.06 - 6.13) | 4.39 (4.07 - 4.73) | <0.001 | 0.46 | <0.001 |
| SP140L | 1.34 (1.09 - 1.56) | 1.11 (0.97 - 1.39) | <0.001 | 0.33 | <0.001 |
| SPAG9 | 1.57 (1.44 - 1.87) | 1.12 (0.89 - 1.33) | <0.001 | 0.36 | <0.001 |
| SPECC1L | 1 (0.76 - 1.18) | 0.78 (0.62 - 1.03) | <0.001 | 0.22 | <0.001 |
| SPG7 | 1.21 (1.01 - 1.42) | 0.79 (0.61 - 1.01) | <0.001 | 0.35 | <0.001 |
| SPOCK2 | 2.51 (1.93 - 2.93) | 1.1 (0.85 - 1.41) | <0.001 | 0.9 | <0.001 |
| SPTAN1 | 1.45 (1.03 - 1.79) | 1.52 (1.09 - 1.93) | <0.001 | -0.38 | 1 |
| SPTBN1 | 1.78 (1.58 - 1.98) | 1.91 (1.62 - 2.1) | 0.72 | -0.21 | 1 |
| SRGN | 1.74 (1.23 - 2.17) | 1.24 (0.97 - 1.97) | <0.001 | 0.27 | <0.001 |
| SRRM2 | 7.88 (7.48 - 8.79) | 8.13 (7.61 - 8.65) | <0.001 | -0.35 | 0.01 |
| SSBP1 | 1.08 (0.84 - 1.24) | 1.03 (0.78 - 1.32) | <0.001 | 0.26 | <0.001 |
| SSH2 | 3.16 (2.62 - 3.43) | 2.46 (2.24 - 2.96) | <0.001 | 0.37 | <0.001 |
| ST6GAL1 | 3.2 (2.4 - 3.73) | 3.01 (2.31 - 3.46) | 0.31 | -0.23 | 1 |
| STAM | 1.83 (1.39 - 2.32) | 0.77 (0.49 - 1.13) | <0.001 | 0.66 | <0.001 |
| STAMBPL1 | 0.96 (0.71 - 1.2) | 0.55 (0.41 - 0.78) | <0.001 | 0.31 | <0.001 |
| STAP1 | 0.86 (0.62 - 1.06) | 0.75 (0.6 - 1) | <0.001 | 0.38 | <0.001 |
| STAT2 | 1.08 (0.79 - 1.39) | 0.7 (0.47 - 0.93) | <0.001 | 0.43 | <0.001 |
| STAU1 | 1.13 (1.02 - 1.21) | 0.94 (0.83 - 1.08) | <0.001 | 0.21 | <0.001 |
| STIM2 | 2.34 (1.89 - 3.3) | 1.53 (1.16 - 1.84) | <0.001 | 0.35 | <0.001 |
| STK17A | 1.92 (1.73 - 2.01) | 1.55 (1.35 - 1.79) | <0.001 | 0.32 | <0.001 |
| STK17B | 1.89 (1.43 - 2.67) | 3.25 (2.39 - 3.88) | <0.001 | -0.77 | <0.001 |
| STK24 | 1.51 (1.27 - 1.92) | 1.54 (1.21 - 1.97) | <0.001 | -0.38 | 0.65 |
| STK4 | 5.13 (4.51 - 5.77) | 4.14 (3.68 - 5.11) | <0.001 | 0.47 | <0.001 |
| STX16 | 2.2 (1.8 - 2.4) | 1.63 (1.49 - 2.11) | <0.001 | 0.44 | <0.001 |
| STX7 | 4.31 (3.23 - 5.66) | 3.19 (2.67 - 3.67) | <0.001 | 0.49 | <0.001 |
| SUB1 | 2 (1.75 - 2.3) | 2.83 (2.51 - 3.1) | <0.001 | -0.55 | <0.001 |
| SUDS3 | 0.89 (0.8 - 1.06) | 0.7 (0.55 - 0.8) | <0.001 | 0.26 | <0.001 |
| SUN1 | 1.26 (1.05 - 1.42) | 0.77 (0.68 - 0.97) | <0.001 | 0.36 | <0.001 |
| SUN2 | 0.58 (0.45 - 0.77) | 0.84 (0.57 - 1.17) | <0.001 | -0.7 | <0.001 |
| SUZ12 | 1.32 (1.09 - 1.56) | 1.5 (1.27 - 1.81) | 0.27 | -0.35 | 1 |
| SYK | 2.27 (1.94 - 3.09) | 2.54 (2.2 - 3.04) | 0.01 | -0.3 | 1 |
| SYNE2 | 4.29 (3.6 - 5.85) | 1.89 (1.46 - 2.44) | <0.001 | 0.84 | <0.001 |
| SYNGR2 | 3.12 (2.79 - 3.53) | 3.29 (3.07 - 3.55) | <0.001 | 0.23 | <0.001 |
| SYVN1 | 0.87 (0.59 - 1.11) | 0.94 (0.64 - 1.23) | 0.13 | -0.21 | 1 |
| TAF1D | 1.94 (1.75 - 2.16) | 1.74 (1.51 - 2.17) | <0.001 | 0.33 | <0.001 |
| TAGAP | 2.59 (2.19 - 3.26) | 2.74 (2.18 - 3.18) | 0.01 | -0.34 | 1 |
| TAGLN2 | 3.33 (2.92 - 3.7) | 5.2 (4.52 - 5.73) | <0.001 | -0.44 | <0.001 |
| TAP2 | 0.84 (0.66 - 1.13) | 1 (0.67 - 1.15) | 0.21 | -0.26 | 1 |

| | | | | | |
|---|---|---|---|---|---|
| TAPBP | 2.91 (2.62 - 3.32) | 3.48 (3.15 - 3.72) | 0.35 | -0.23 | 1 |
| TBC1D1 | 2.04 (1.73 - 2.19) | 1.23 (1.07 - 1.4) | <0.001 | 0.34 | <0.001 |
| TBC1D22A | 1.76 (1.43 - 1.96) | 1.12 (0.87 - 1.45) | <0.001 | 0.44 | <0.001 |
| TBRG1 | 1.22 (1.01 - 1.33) | 0.93 (0.8 - 1.18) | <0.001 | 0.41 | <0.001 |
| TCEA1 | 3.12 (2.97 - 3.52) | 3.59 (3.52 - 3.68) | 0.47 | -0.23 | 1 |
| TCF12 | 1.11 (0.96 - 1.22) | 0.79 (0.68 - 0.9) | <0.001 | 0.26 | <0.001 |
| TCF25 | 2.72 (2.22 - 3.51) | 2.33 (1.93 - 2.89) | <0.001 | 0.37 | <0.001 |
| TCF4 | 6.02 (4.95 - 6.92) | 3.3 (2.56 - 4.1) | <0.001 | 0.69 | <0.001 |
| TCL1A | 3.29 (2.39 - 4.67) | 2.18 (1.43 - 2.49) | <0.001 | 0.5 | <0.001 |
| TCP11L2 | 1.62 (1.02 - 2.08) | 1.02 (0.7 - 1.45) | <0.001 | 0.46 | <0.001 |
| TERF2 | 0.68 (0.59 - 0.83) | 1.05 (0.82 - 1.24) | <0.001 | -0.64 | <0.001 |
| TESC | 1.72 (1.46 - 2.04) | 0.78 (0.5 - 1.11) | <0.001 | 0.75 | <0.001 |
| TGFB1 | 1.41 (1.31 - 1.53) | 1.72 (1.44 - 1.94) | <0.001 | -0.57 | <0.001 |
| THRAP3 | 2.48 (2.34 - 2.67) | 2.26 (2.04 - 2.54) | <0.001 | 0.2 | <0.001 |
| THUMPD3-AS1 | 0.92 (0.77 - 1.08) | 0.48 (0.38 - 0.65) | <0.001 | 0.49 | <0.001 |
| TKT | 0.51 (0.37 - 0.75) | 0.93 (0.65 - 1.22) | <0.001 | -0.78 | <0.001 |
| TLE4 | 1.97 (1.51 - 2.66) | 0.99 (0.67 - 1.31) | <0.001 | 0.53 | <0.001 |
| TLE5 | 2.03 (1.79 - 2.2) | 3.12 (2.83 - 3.57) | <0.001 | -0.81 | <0.001 |
| TLK1 | 4.26 (3.78 - 4.59) | 3.11 (2.85 - 3.79) | <0.001 | 0.49 | <0.001 |
| TLN1 | 2.21 (1.8 - 2.58) | 2.64 (2.07 - 3.51) | <0.001 | -0.64 | <0.001 |
| TLR10 | 0.38 (0.12 - 0.62) | 0.79 (0.51 - 1.23) | <0.001 | -0.84 | <0.001 |
| TM9SF3 | 1.22 (1.02 - 1.44) | 1.03 (0.9 - 1.29) | 0.11 | -0.26 | 1 |
| TMED5 | 1.24 (1.09 - 1.44) | 1.36 (1.09 - 1.47) | 0.42 | -0.24 | 1 |
| TMED8 | 0.95 (0.83 - 1.05) | 0.93 (0.75 - 1.17) | 0.46 | -0.27 | 1 |
| TMEM131L | 2.88 (2.74 - 3.15) | 1.67 (1.48 - 1.82) | <0.001 | 0.56 | <0.001 |
| TMEM154 | 1.13 (0.74 - 1.31) | 1.26 (1 - 1.71) | <0.001 | -0.45 | 1 |
| TMEM19 | 0.78 (0.61 - 0.94) | 0.66 (0.62 - 0.71) | <0.001 | 0.29 | <0.001 |
| TMEM30A | 1.06 (0.98 - 1.27) | 1.05 (0.75 - 1.2) | 0.79 | -0.21 | 1 |
| TMPO | 2.1 (1.79 - 2.77) | 1.71 (1.15 - 1.99) | <0.001 | 0.37 | <0.001 |
| TNKS | 1.01 (0.89 - 1.1) | 0.74 (0.66 - 0.94) | <0.001 | 0.24 | <0.001 |
| TNRC6A | 1.32 (1.18 - 1.53) | 0.87 (0.8 - 0.93) | <0.001 | 0.23 | <0.001 |
| TNRC6B | 3.04 (2.8 - 3.33) | 2.39 (2.21 - 2.56) | <0.001 | 0.31 | <0.001 |
| TOMM6 | 1.3 (0.89 - 1.62) | 1.34 (1.05 - 1.75) | <0.001 | 0.2 | <0.001 |
| TPI1 | 1.33 (1.15 - 1.62) | 1.49 (1.22 - 1.72) | <0.001 | 0.22 | <0.001 |
| TPM3 | 5.06 (4.19 - 5.84) | 5.84 (5 - 6.58) | <0.001 | -0.35 | 0.14 |
| TPM4 | 1.74 (1.47 - 2.21) | 1.95 (1.55 - 2.28) | 0.67 | -0.22 | 1 |
| TPP1 | 0.92 (0.76 - 1.06) | 0.88 (0.66 - 1.16) | 0.11 | -0.28 | 1 |
| TPR | 2.48 (2.29 - 2.63) | 2.02 (1.79 - 2.18) | <0.001 | 0.23 | <0.001 |
| TPT1 | 27.89 (25.76 - 30.26) | 35.03 (33.09 - 37.45) | <0.001 | -0.4 | <0.001 |

| | | | | | |
|---|---|---|---|---|---|
| TRA2A | 1.13 (1.04 - 1.25) | 0.97 (0.9 - 1.17) | <0.001 | 0.24 | <0.001 |
| TRAC | 1.1 (0.81 - 1.47) | 0.85 (0.77 - 0.93) | <0.001 | 0.82 | <0.001 |
| TRAF3IP3 | 1.72 (1.22 - 2) | 1.11 (0.99 - 1.38) | <0.001 | 0.62 | <0.001 |
| TRAF5 | 1.76 (1.32 - 2.1) | 1.28 (1 - 1.54) | <0.001 | 0.41 | <0.001 |
| TRAK1 | 1.14 (0.94 - 1.27) | 1.12 (0.98 - 1.33) | 0.4 | -0.2 | 1 |
| TRAPPC2L | 1.15 (0.91 - 1.29) | 0.81 (0.6 - 1.18) | <0.001 | 0.53 | <0.001 |
| TRBC2 | 1.74 (1.46 - 2.45) | 1.64 (1.38 - 2.06) | <0.001 | 0.21 | <0.001 |
| TRIM73 | 0.88 (0.61 - 1.09) | 0.49 (0.35 - 0.69) | <0.001 | 0.62 | <0.001 |
| TRIP11 | 0.86 (0.79 - 0.91) | 0.64 (0.56 - 0.76) | <0.001 | 0.22 | <0.001 |
| TRIR | 1.89 (1.7 - 2.11) | 2.44 (2.18 - 2.65) | <0.001 | -0.42 | 1 |
| TSC22D3 | 13.28 (12.29 - 14.52) | 13.47 (12.48 - 14.52) | <0.001 | -0.36 | <0.001 |
| TSPAN3 | 0.77 (0.64 - 1) | 1.43 (1.17 - 1.76) | <0.001 | -0.69 | <0.001 |
| TSPYL2 | 2.86 (2.11 - 4.28) | 1.68 (1.29 - 2.1) | <0.001 | 0.69 | <0.001 |
| TSTD1 | 1.39 (1.07 - 1.77) | 1.36 (1.02 - 1.89) | <0.001 | 0.33 | <0.001 |
| TTC14 | 1.21 (1.07 - 1.36) | 0.83 (0.76 - 0.86) | <0.001 | 0.35 | <0.001 |
| TTC17 | 1.64 (1.54 - 1.76) | 1.07 (0.88 - 1.15) | <0.001 | 0.32 | <0.001 |
| TTN | 10.77 (8.66 - 14.33) | 5.04 (2.89 - 6.57) | <0.001 | 0.94 | <0.001 |
| TUBA1A | 1.32 (0.99 - 1.61) | 1 (0.84 - 1.18) | <0.001 | 0.46 | <0.001 |
| TUBA1B | 2.5 (2.1 - 2.92) | 2.5 (2.21 - 2.8) | <0.001 | 0.23 | <0.001 |
| TUBB | 1.53 (1.17 - 2.21) | 1.99 (1.41 - 2.39) | <0.001 | -0.35 | 1 |
| TUBGCP2 | 0.87 (0.81 - 0.98) | 0.71 (0.61 - 0.8) | <0.001 | 0.34 | <0.001 |
| TUT4 | 3.98 (3.36 - 4.37) | 2.88 (2.57 - 3.42) | <0.001 | 0.46 | <0.001 |
| TXNDC5 | 1.61 (1.24 - 1.9) | 1.76 (1.46 - 2.23) | <0.001 | -0.77 | <0.001 |
| TYROBP | 0.8 (0.5 - 1.9) | 0.5 (0.18 - 0.6) | <0.001 | 0.53 | <0.001 |
| UBAC2 | 0.96 (0.87 - 1.09) | 0.67 (0.54 - 0.74) | <0.001 | 0.31 | <0.001 |
| UBE2D2 | 3.01 (2.79 - 3.13) | 2.56 (2.44 - 2.71) | <0.001 | 0.21 | <0.001 |
| UBE2D3 | 2.85 (2.64 - 3.25) | 3.2 (2.69 - 3.5) | 0.39 | -0.26 | 1 |
| UBE2H | 1.07 (0.96 - 1.14) | 0.97 (0.9 - 1.14) | 0.4 | -0.23 | 1 |
| UBE2J1 | 1.29 (1.11 - 1.48) | 2.31 (2.04 - 2.81) | <0.001 | -1.01 | <0.001 |
| UBE2N | 1.01 (0.79 - 1.27) | 1.13 (0.75 - 1.45) | 0.17 | -0.32 | 1 |
| UBE3A | 2.13 (1.8 - 2.43) | 1.49 (1.24 - 1.69) | <0.001 | 0.32 | <0.001 |
| UHMK1 | 1.57 (1.43 - 1.78) | 1.75 (1.49 - 1.89) | 0.29 | -0.27 | 1 |
| UPF3A | 0.89 (0.82 - 1.03) | 0.76 (0.68 - 0.88) | <0.001 | 0.28 | <0.001 |
| USP11 | 1.22 (1 - 1.39) | 0.98 (0.81 - 1.22) | <0.001 | 0.23 | <0.001 |
| USP15 | 3.95 (3.69 - 4.24) | 2.9 (2.61 - 3.29) | <0.001 | 0.67 | <0.001 |
| USP24 | 1.39 (1.19 - 1.55) | 1.31 (1.08 - 1.52) | 0.23 | -0.21 | 1 |
| USP3 | 2.02 (1.72 - 2.18) | 1.9 (1.62 - 2.31) | <0.001 | 0.2 | <0.001 |
| USP47 | 1.77 (1.57 - 2) | 1.15 (1.08 - 1.23) | <0.001 | 0.33 | <0.001 |
| USP8 | 2.43 (2.22 - 2.57) | 1.74 (1.66 - 1.89) | <0.001 | 0.41 | <0.001 |

| | | | | | |
|---|---|---|---|---|---|
| USP9Y | 1.07 (0 - 1.95) | 0.63 (0 - 1.17) | <0.001 | 0.23 | <0.001 |
| UTY | 2.12 (0 - 3.52) | 1.03 (0 - 1.5) | <0.001 | 0.51 | <0.001 |
| UVRAG | 5.44 (3.88 - 9.71) | 4 (2.44 - 4.87) | <0.001 | 0.53 | <0.001 |
| VAMP2 | 3.14 (2.83 - 3.63) | 3.85 (3.52 - 4.23) | 0.44 | -0.25 | 1 |
| VASP | 0.58 (0.42 - 0.79) | 0.8 (0.63 - 1.01) | <0.001 | -0.36 | 0.16 |
| VPS29 | 0.88 (0.8 - 0.97) | 0.66 (0.57 - 0.75) | <0.001 | 0.35 | <0.001 |
| VTI1A | 1.06 (0.96 - 1.14) | 0.84 (0.73 - 0.95) | <0.001 | 0.31 | <0.001 |
| WAC | 2.04 (1.72 - 2.26) | 2.15 (1.72 - 2.44) | 0.29 | -0.3 | 1 |
| WAS | 1.04 (0.9 - 1.28) | 1.26 (1 - 1.45) | 0.67 | -0.2 | 1 |
| WASF2 | 3.56 (3.2 - 3.89) | 3.41 (2.74 - 4.05) | 0.77 | -0.23 | 1 |
| WDR11 | 2.19 (1.74 - 2.36) | 1.48 (1.25 - 1.79) | <0.001 | 0.31 | <0.001 |
| WEE1 | 0.75 (0.55 - 0.93) | 0.93 (0.66 - 1.32) | <0.001 | -0.5 | <0.001 |
| WIPF1 | 1.71 (1.4 - 1.99) | 1.93 (1.57 - 2.21) | <0.001 | -0.4 | 0.45 |
| WNK1 | 1.54 (1.37 - 1.73) | 1.44 (1.24 - 1.76) | 0.01 | -0.3 | 1 |
| YBX1 | 5.61 (5.16 - 6.83) | 7.53 (6.07 - 8.77) | <0.001 | -0.6 | <0.001 |
| YBX3 | 6.57 (5.52 - 8.27) | 3.52 (2.85 - 3.92) | <0.001 | 0.55 | <0.001 |
| YPEL2 | 0.98 (0.82 - 1.11) | 0.57 (0.5 - 0.8) | <0.001 | 0.4 | <0.001 |
| YPEL3 | 1.87 (1.65 - 2.01) | 1.83 (1.64 - 2) | 0.03 | -0.31 | 1 |
| YTHDC2 | 1.44 (1.35 - 1.56) | 1.04 (1.01 - 1.1) | <0.001 | 0.24 | <0.001 |
| YWHAB | 2.83 (2.59 - 3.34) | 4.29 (3.86 - 4.66) | <0.001 | -0.59 | <0.001 |
| YWHAH | 1.49 (1.27 - 1.67) | 1.67 (1.44 - 2.19) | <0.001 | -0.48 | <0.001 |
| YWHAZ | 12.01 (10.42 - 14.87) | 13.11 (9.83 - 14.15) | <0.001 | -0.25 | <0.001 |
| ZBED6 | 1.08 (1.01 - 1.14) | 0.61 (0.56 - 0.66) | <0.001 | 0.26 | <0.001 |
| ZBTB20 | 1.67 (1.42 - 1.92) | 1.32 (1.13 - 1.58) | <0.001 | 0.36 | <0.001 |
| ZBTB24 | 1.31 (1.04 - 1.57) | 0.84 (0.57 - 1.16) | <0.001 | 0.42 | <0.001 |
| ZBTB38 | 1.07 (0.88 - 1.18) | 1.25 (1.12 - 1.57) | <0.001 | -0.42 | 1 |
| ZCCHC7 | 2.7 (2.51 - 3.1) | 1.49 (1.27 - 1.66) | <0.001 | 0.7 | <0.001 |
| ZEB2 | 2.37 (1.81 - 3.13) | 2.87 (1.95 - 4.2) | 0.14 | -0.46 | 1 |
| ZFAND5 | 1.12 (0.94 - 1.37) | 1.35 (1.03 - 1.61) | <0.001 | -0.36 | 0.23 |
| ZFAND6 | 0.79 (0.73 - 0.88) | 1.23 (1.09 - 1.35) | <0.001 | -0.44 | 0.02 |
| ZFC3H1 | 2.13 (1.96 - 2.3) | 1.69 (1.54 - 2.06) | <0.001 | 0.4 | <0.001 |
| ZFP36 | 7.23 (6.72 - 8.7) | 8.87 (7 - 10.68) | 0.01 | -0.3 | 1 |
| ZMYM2 | 1.73 (1.3 - 1.89) | 1.36 (1.15 - 2.09) | <0.001 | 0.3 | <0.001 |
| ZNF107 | 2.99 (2.54 - 3.82) | 1.73 (1.4 - 2.14) | <0.001 | 0.68 | <0.001 |
| ZNF141 | 1.08 (0.86 - 1.22) | 0.57 (0.44 - 0.74) | <0.001 | 0.3 | <0.001 |
| ZNF266 | 1.21 (0.97 - 1.56) | 0.67 (0.48 - 0.89) | <0.001 | 0.64 | <0.001 |
| ZNF292 | 2.84 (2.56 - 3.14) | 2.14 (1.98 - 2.38) | <0.001 | 0.2 | <0.001 |
| ZNF331 | 2.26 (1.11 - 4.21) | 2.09 (1.34 - 3.47) | <0.001 | 0.39 | <0.001 |
| ZNF394 | 1.24 (1.09 - 1.36) | 1.12 (0.99 - 1.35) | <0.001 | 0.22 | <0.001 |

| | | | | | |
|---|---|---|---|---|---|
| ZNF506 | 1.17 (0.95 - 1.4) | 0.73 (0.62 - 0.86) | <0.001 | 0.3 | <0.001 |
| ZNF638 | 1.82 (1.56 - 1.95) | 1.21 (1.08 - 1.41) | <0.001 | 0.39 | <0.001 |
| ZNF644 | 2.11 (1.96 - 2.26) | 1.74 (1.54 - 1.89) | <0.001 | 0.31 | <0.001 |
| ZNF721 | 1.28 (1.04 - 1.39) | 0.95 (0.87 - 1.01) | <0.001 | 0.21 | <0.001 |
| ZNF800 | 1.96 (1.47 - 2.17) | 1.71 (1.45 - 2.3) | <0.001 | 0.23 | <0.001 |
| ZNF92 | 1.31 (1.13 - 1.44) | 1 (0.87 - 1.13) | <0.001 | 0.33 | <0.001 |
| ZRANB2 | 2.24 (1.98 - 2.39) | 1.73 (1.56 - 2.01) | <0.001 | 0.3 | <0.001 |

**Table S2: Enriched GO-terms between CLL and NAC cells.** Enrichment was calculated with clusterProfilers enrichGO.

### A) NAC-CLL (low z-score) vs. CLL cells (high z-score)

| Term | Enrich | $log_{10}$(adjusted pval) | z-score | Mean $L_2FC$ |
|---|---|---|---|---|
| regulation of cell morphogenesis | 0.039 | -5.487 | -3.429 | -0.154 |
| B cell activation | 0.051 | -9.463 | -3.255 | -0.171 |
| Ras protein signal transduction | 0.042 | -5.171 | -3.175 | -0.163 |
| actin filament organization | 0.056 | -7.827 | -3.101 | -0.102 |
| dephosphorylation | 0.042 | -5.402 | -3 | -0.157 |
| myeloid cell differentiation | 0.049 | -6.515 | -2.782 | -0.126 |
| regulation of cellular component size | 0.047 | -6.84 | -2.722 | -0.125 |
| protein dephosphorylation | 0.035 | -5.881 | -2.655 | -0.151 |
| regulation of mRNA metabolic process | 0.054 | -12.801 | -2.54 | -0.072 |
| positive regulation of cellular catabolic process | 0.048 | -5.521 | -2.405 | -0.103 |
| positive regulation of cell adhesion | 0.061 | -8.654 | -2.39 | -0.076 |
| leukocyte cell-cell adhesion | 0.062 | -11.995 | -2.357 | -0.057 |
| antigen processing and presentation of exogenous peptide antigen | 0.012 | -5.621 | -2.138 | -0.147 |
| regulation of translation | 0.06 | -8.174 | -2.047 | -0.083 |
| regulation of actin filament-based process | 0.048 | -5.888 | -2.023 | -0.082 |
| RNA catabolic process | 0.042 | -7.15 | -2.021 | -0.054 |
| regulation of RNA splicing | 0.037 | -11.09 | -1.982 | -0.079 |
| positive regulation of cellular component biogenesis | 0.057 | -6.737 | -1.969 | -0.068 |
| lymphocyte differentiation | 0.061 | -10.893 | -1.912 | -0.057 |
| regulation of supramolecular fiber organization | 0.047 | -6.244 | -1.905 | -0.059 |
| mononuclear cell proliferation | 0.048 | -9.872 | -1.871 | -0.12 |
| nucleobase-containing compound catabolic process | 0.048 | -5.552 | -1.871 | -0.043 |
| regulation of hemopoiesis | 0.055 | -8.988 | -1.75 | -0.024 |
| regulation of protein modification by small protein conjugation or removal | 0.035 | -6.244 | -1.718 | -0.086 |
| nucleocytoplasmic transport | 0.043 | -6.737 | -1.697 | -0.053 |
| positive regulation of lymphocyte activation | 0.054 | -9.087 | -1.638 | -0.027 |
| regulation of intracellular transport | 0.048 | -8.527 | -1.483 | -0.084 |
| non-membrane-bounded organelle assembly | 0.048 | -6.883 | -1.336 | -0.041 |
| viral process | 0.048 | -5.77 | -1.336 | -0.045 |
| immune response-regulating cell surface receptor signaling pathway | 0.05 | -9.463 | -1.313 | -0.063 |
| activation of immune response | 0.052 | -8.02 | -1.291 | -0.072 |
| histone modification | 0.054 | -6.146 | -1.27 | -0.079 |
| alpha-beta T cell activation | 0.029 | -6.407 | -1.219 | -0.051 |
| regulation of protein-containing complex assembly | 0.048 | -6.101 | -1.069 | -0.056 |
| negative regulation of immune system process | 0.05 | -5.588 | -1.05 | -0.009 |
| leukocyte activation involved in immune response | 0.039 | -6.186 | -1.043 | -0.029 |
| positive regulation of endopeptidase activity | 0.027 | -5.501 | -0.898 | -0.043 |
| positive regulation of cytokine production | 0.056 | -6.756 | -0.868 | -0.003 |
| RNA splicing | 0.063 | -10.055 | -0.819 | -0.036 |
| intrinsic apoptotic signaling pathway | 0.042 | -7.602 | -0.714 | -0.08 |
| protein localization to nucleus | 0.04 | -6.078 | -0.59 | -0.034 |
| regulation of immune effector process | 0.048 | -7.153 | -0.267 | -0.003 |
| establishment of protein localization to organelle | 0.06 | -9.46 | 0.12 | -0.01 |
| maintenance of location in cell | 0.032 | -5.712 | 0.164 | 0.056 |
| regulation of protein stability | 0.049 | -9.721 | 0.397 | 0.009 |
| protein targeting | 0.043 | -6.88 | 0.849 | 0.003 |
| response to virus | 0.052 | -7.57 | 1.291 | 0.057 |
| ribonucleoprotein complex biogenesis | 0.058 | -7.354 | 2.321 | 0.061 |
| aerobic respiration | 0.032 | -7.278 | 3.138 | 0.151 |
| oxidative phosphorylation | 0.028 | -7.57 | 3.71 | 0.192 |

**Table S3: Cells included in BCR deep sequencing and number of sequences retrieved by NGS.**

| ID | year of PB sampling | sorted as | # of sorted cells[a] | # of reads | # of reads after quality control | # of collapsed reads | total IGHV rearrangement sequences | # CLL rearrangement sequences | CLL rearrangement per sequences with IGHV [%] |
|---|---|---|---|---|---|---|---|---|---|
| 03 | 2012 | CLL | 300,000 | 821,245 | 731,969 | 49,898 | 10,486 | 1,410 | 98.45 |
| 03 | 2012 | CLL | 300,000 | 1,160,887 | 1,040,004 | 66,278 | 12,821 | 1,692 | 98.32 |
| 03 | 2012 | NRB | 26,000 | 813,377 | 632,226 | 172,399 | 25,779 | 4,347 | 0.17 |
| 03 | 2012 | NRB | 30,000 | 619,396 | 482,353 | 153,693 | 27,737 | 3,422 | 1.19 |
| 03 | 2012 | CLL | 300,000 | 599,104 | 503,123 | 32,337 | 8,402 | 1,126 | 98.67 |
| 03 | 2012 | CLL | 300,000 | 603,819 | 479,822 | 38,296 | 8,718 | 1,209 | 97.82 |
| 03 | 2012 | NRB | 6,000 | 545,929 | 3,459 | 1,423 | 144 | 19 | 23.61 |
| 03 | 2012 | NRB | 16,000 | 755,783 | 330,958 | 97,836 | 15,648 | 1,837 | 0.28 |
| 03 | 2014 | CLL | 300,000 | 711,302 | 489,192 | 32,320 | 8,146 | 1,082 | 98.05 |
| 03 | 2014 | CLL | 300,000 | 591,773 | 488,375 | 28,641 | 7,794 | 1,053 | 98.43 |
| 03 | 2014 | NRB | 10,150 | 879,880 | 378,860 | 38,263 | 8,069 | 993 | 86.67 |
| 03 | 2014 | NRB | 12,000 | 532,273 | 279,069 | 26,161 | 6,224 | 791 | 90.30 |
| 03 | 2016 | CLL | 150,000 | 712,385 | 595,589 | 38,610 | 9,461 | 1,239 | 98.51 |
| 03 | 2016 | CLL | 150,000 | 691,335 | 558,075 | 45,872 | 9,933 | 1,281 | 97.82 |
| 03 | 2016 | NRB | 3750 | 699,548 | 35,756 | 8,895 | 1,227 | 60 | 29.58 |
| 03 | 2016 | NRB | 6000 | 985,839 | 138,752 | 32,855 | 4,280 | 174 | 30.61 |
| 03 | 2018 | CLL | 300,000 | 709,604 | 594,504 | 40,963 | 9,592 | 1,246 | 98.35 |
| 03 | 2018 | CLL | 300,000 | 558,205 | 467,936 | 30,533 | 8,160 | 1,064 | 98.20 |
| 03 | 2018 | NRB | 7,250 | 689,910 | 284,300 | 26,718 | 6,494 | 820 | 92.29 |
| 03 | 2018 | NRB | 1,250 | 787,038 | 27,161 | 5,685 | 0 | 0 | 0.00 |
| 03 | 2018 | CLL | 300,000 | 641,474 | 547,237 | 35,984 | 9,127 | 1,199 | 98.33 |
| 03 | 2018 | CLL | 300,000 | 569,242 | 495,649 | 35,814 | 8,595 | 1,121 | 98.70 |
| 03 | 2018 | NRB | 10,000 | 645,255 | 410,207 | 42,745 | 8,585 | 1,051 | 94.98 |
| 03 | 2018 | NRB | 12,750 | 863,291 | 573,429 | 47,530 | 10,036 | 1,304 | 96.17 |
| 04 | 2006 | CLL | 300,000 | 12,040 | 459 | 264 | 36 | 2 | 5.56 |
| 04 | 2006 | CLL | 300,000 | 1,045,988 | 763,416 | 41,021 | 5,078 | 4,883 | 96.16 |
| 04 | 2006 | NRB | 26,000 | 995,810 | 205,031 | 74,748 | 9,189 | 133 | 1.45 |
| 04 | 2006 | NRB | 15,000 | 1,164,506 | 131,824 | 44,991 | 5,087 | 42 | 0.83 |
| 04 | 2009 | CLL | 300,000 | 9,185 | 827 | 588 | 41 | 5 | 12.2 |
| 04 | 2009 | CLL | 300,000 | 1,193,630 | 751,490 | 52,185 | 7,755 | 7,388 | 95.27 |

| | | | | | | | | | |
|---|---|---|---|---|---|---|---|---|---|
| 04 | 2009 | NRB | 8,000 | 489,794 | 32,129 | 12,301 | 1,831 | 127 | 6.94 |
| 04 | 2009 | NRB | 12,000 | 988,115 | 89,294 | 30,708 | 3,840 | 338 | 8.8 |
| 04 | 2017 | CLL | 300,000 | 16,564 | 2,268 | 1,001 | 100 | 1 | 1 |
| 04 | 2017 | CLL | 300,000 | 31,965 | 8,288 | 3,313 | 472 | 117 | 24.79 |
| 04 | 2017 | NRB | 50,000 | 1,080,969 | 689,206 | 259,076 | 48,624 | 64 | 0.13 |
| 04 | 2017 | NRB | 65,000 | 1,138,010 | 798,005 | 352,900 | 62,883 | 81 | 0.13 |
| 04 | 2018 | CLL | 300,000 | 999,168 | 433,684 | 24,301 | 3,757 | 3,598 | 95.77 |
| 04 | 2018 | CLL | 300,000 | 1,420,756 | 782,408 | 42,663 | 6,445 | 6,226 | 96.6 |
| 04 | 2018 | NRB | 16,000 | 1,292,963 | 547,665 | 101,939 | 14,961 | 6,208 | 41.49 |
| 04 | 2018 | NRB | 17,000 | 902,447 | 604,118 | 107,795 | 16,672 | 6,158 | 36.94 |
| 09 | 2018 | CLL | 500,000 | 737,464 | 178,981 | 18,339 | 4,228 | 3,338 | 78.95 |
| 09 | 2018 | CLL | 500,000 | 1,123,530 | 594,240 | 40,189 | 8,952 | 6,348 | 70.91 |
| 09 | 2018 | NRB | 50,000 | 890,079 | 406,510 | 39,330 | 8,283 | 5,600 | 67.61 |
| 09 | 2018 | NRB | 50,000 | 864,744 | 187,912 | 27,015 | 4,806 | 3,717 | 77.34 |
| 09 | 2018 | CLL | 250,000 | 689,400 | 583,755 | 61,504 | 11,628 | 10,726 | 92.24 |
| 09 | 2018 | CLL | 250,000 | 747,219 | 656,884 | 53,174 | 10,757 | 10,490 | 97.52 |
| 09 | 2018 | NRB | 25,000 | 1,112,890 | 36,781 | 9,397 | 1,582 | 1,465 | 92.6 |
| 09 | 2018 | NRB | 25,000 | 503,036 | 228,749 | 27,890 | 5,242 | 4,470 | 85.27 |
| 10 | 2008 | CLL | 300,000 | 653,204 | 360,572 | 30,562 | 7,377 | 7,223 | 97.91 |
| 10 | 2008 | CLL | 300,000 | 641,906 | 354,130 | 29,326 | 7,269 | 7,126 | 98.03 |
| 10 | 2008 | NRB | 37,000 | 1,037,292 | 355,837 | 146,050 | 27,589 | 62 | 0.22 |
| 10 | 2008 | NRB | 25,000 | 914,510 | 333,637 | 104,747 | 17,758 | 118 | 0.66 |
| 10 | 2013 | CLL | 300,000 | 636,339 | 342,461 | 30,303 | 7,430 | 7,282 | 98.01 |
| 10 | 2013 | CLL | 300,000 | 581,727 | 320,581 | 27,149 | 6,842 | 6,698 | 97.9 |
| 10 | 2013 | NRB | 5,000 | 1,353,184 | 73,689 | 18,712 | 2,484 | 160 | 6.44 |
| 10 | 2013 | NRB | 15,000 | 1,201,706 | 353,067 | 70,426 | 9,495 | 753 | 7.93 |
| 11 | 2018 | CLL | 500,000 | 637,499 | 177,030 | 16,710 | 4,255 | 4,101 | 96.38 |
| 11 | 2018 | CLL | 500,000 | 781,471 | 317,293 | 24,038 | 7,062 | 6,923 | 98.03 |
| 11 | 2018 | NRB | 15,000 | 1,394,953 | 496,259 | 72,039 | 13,251 | 8,309 | 62.7 |
| 11 | 2018 | NRB | 10,000 | 716,995 | 259,262 | 43,603 | 8,898 | 5,703 | 64.09 |
| 11 | 2018 | CLL | 300,000 | 949,206 | 782,711 | 68,858 | 11,931 | 11,662 | 97.75 |
| 11 | 2018 | CLL | 300,000 | 756,055 | 646,577 | 53,784 | 10,193 | 9,991 | 98.02 |
| 11 | 2018 | NRB | 29,000 | 888,938 | 556,041 | 150,602 | 22,726 | 6,285 | 27.66 |
| 11 | 2018 | NRB | 23,000 | 1,107,784 | 469,298 | 129,247 | 19,290 | 5,858 | 30.37 |

| | | | | | | | | | |
|---|---|---|---|---|---|---|---|---|---|
| 12 | 2013 | CLL | 200,000 | 737,707 | 530,210 | 33,509 | 3,985 | 2,426 | 60.88 |
| 12 | 2013 | CLL | 110,000 | 677,509 | 582,893 | 28,114 | 4,457 | 2,243 | 50.32 |
| 12 | 2013 | NRB | 19,150 | 539,488 | 74,582 | 21,310 | 14,243 | 244 | 1.71 |
| 12 | 2013 | NRB | 10,150 | 796,122 | 62,855 | 14,348 | 9,659 | 285 | 2.95 |
| 12 | 2015 | CLL | 300,000 | 617,587 | 580,842 | 29,139 | 4,402 | 2,288 | 51.98 |
| 12 | 2015 | CLL | 300,000 | 806,781 | 711,238 | 35,895 | 4,258 | 2,647 | 62.17 |
| 12 | 2015 | NRB | 2,800 | 781,707 | 52,921 | 15,048 | 3,252 | 87 | 2.68 |
| 12 | 2015 | NRB | 3,000 | 695,428 | 78,466 | 21,930 | 1,863 | 128 | 6.87 |
| 12 | 2018 | CLL | 300,000 | 716,414 | 754,014 | 61,826 | 4,201 | 3,498 | 83.27 |
| 12 | 2018 | CLL | 300,000 | 587,166 | 665,246 | 35,101 | 4,841 | 2,926 | 60.44 |
| 12 | 2018 | NRB | 7,500 | 920,158 | 480,851 | 163,563 | 1,749 | 153 | 8.75 |
| 12 | 2018 | NRB | 7,500 | 933,616 | 405,483 | 135,108 | 2,481 | 74 | 2.98 |
| 12 | 2006 | CLL | 300,000 | 578,545 | 507,446 | 24,302 | 6,441 | 2,059 | 31.97 |
| 12 | 2006 | CLL | 300,000 | 96,243 | 21,799 | 3,900 | 5,330 | 214 | 4.02 |
| 12 | 2006 | NRB | 33,000 | 797,505 | 386,247 | 107,239 | 20,529 | 427 | 2.08 |
| 12 | 2006 | NRB | 23,000 | 508,179 | 247,663 | 57,373 | 17,434 | 300 | 1.72 |
| 13 | 2006 | NRB | 3,500 | 20,368 | 6,023 | 1,306 | 253 | 4 | 1.58 |
| 13 | 2006 | NRB | 13,500 | 745,369 | 650,866 | 38,041 | 8,818 | 8,733 | 99.04 |
| 13 | 2006 | CLL | 300,000 | 1,186,471 | 619,379 | 216,132 | 26,771 | 7,464 | 27.88 |
| 13 | 2006 | CLL | 200,000 | 1,059,955 | 392,342 | 139,809 | 20,100 | 4,620 | 22.99 |
| 13 | 2012 | NRB | 2,000 | 12,494 | 2,304 | 2,041 | 85 | 17 | 20 |
| 13 | 2012 | NRB | 2,000 | 18,439 | 1,279 | 791 | 61 | 29 | 47.54 |
| 13 | 2012 | CLL | 300,000 | 1,485,333 | 381,224 | 81,823 | 12,502 | 6,581 | 52.64 |
| 13 | 2012 | CLL | 300,000 | 1,077,078 | 168,938 | 41,067 | 6,529 | 3,835 | 58.74 |
| 13 | 2016 | NRB | 2,000 | 12,263 | 536 | 1 | 27 | 2 | 7.41 |
| 13 | 2016 | NRB | 2,000 | 4,700 | 211 | 165 | 10 | 3 | 30 |
| 13 | 2016 | CLL | 300,000 | 1,173,120 | 454,521 | 102,190 | 15,046 | 8,241 | 54.77 |
| 13 | 2016 | CLL | 300,000 | 885,765 | 285,218 | 73,158 | 10,729 | 5,077 | 47.32 |
| 14 | 2018 | CLL | 350,000 | 749,086 | 243,865 | 17,773 | 4,470 | 4,153 | 92.91 |
| 14 | 2018 | CLL | 350,000 | 590,483 | 345,584 | 24,041 | 6,239 | 5,785 | 92.72 |
| 14 | 2018 | NRB | 10,000 | 741,055 | 92,227 | 20,304 | 2,423 | 184 | 7.59 |
| 14 | 2018 | NRB | 10,000 | 751,472 | 160,424 | 18,406 | 3,779 | 216 | 5.72 |
| 15 | 2018 | CLL | 500,000 | 973,458 | 511,444 | 38,130 | 8,410 | 7,928 | 94.27 |
| 15 | 2018 | CLL | 500,000 | 1,073,185 | 677,560 | 43,531 | 10,023 | 9,312 | 92.91 |

| | | | | | | | | | |
|---|---|---|---|---|---|---|---|---|---|
| 15 | 2018 | NRB | 150,000 | 1,004,498 | 500,236 | 41,242 | 8,289 | 7,526 | 90.8 |
| 15 | 2018 | NRB | 150,000 | 942,559 | 477,081 | 37,744 | 7,402 | 6,793 | 91.77 |
| 16 | 2018 | CLL | 500,000 | 800,139 | 250,288 | 20,099 | 5,258 | 5,169 | 98.31 |
| 16 | 2018 | CLL | 500,000 | 881,043 | 576,114 | 31,112 | 8,617 | 8,413 | 97.63 |
| 16 | 2018 | NRB | 200,000 | 1,023,993 | 513,297 | 38,323 | 8,872 | 8,449 | 95.23 |
| 16 | 2018 | NRB | 200,000 | 1,650,412 | 1,021,723 | 76,834 | 14,599 | 14,025 | 96.07 |
| 17 | 2010 | CLL | 13,000 | 655,023 | 100,605 | 11,543 | 2,394 | 2,251 | 94.03 |
| 17 | 2010 | CLL | 13,000 | 179,144 | 42,696 | 5,224 | 1,022 | 977 | 95.6 |
| 17 | 2012 | CLL | 58,000 | 545,025 | 129,139 | 14,172 | 3,117 | 2,773 | 88.96 |
| 17 | 2012 | CLL | 58,000 | 428,305 | 145,398 | 14,218 | 3,114 | 2,873 | 92.26 |
| 17 | 2018 | CLL | 500,000 | 701,338 | 349,453 | 27,758 | 7,364 | 7,121 | 96.7 |
| 17 | 2018 | CLL | 500,000 | 1,139,250 | 528,211 | 45,213 | 11,152 | 10,902 | 97.76 |
| 17 | 2018 | NRB | 50,000 | 866,151 | 453,723 | 166,880 | 36,419 | 170 | 0.47 |
| 17 | 2018 | NRB | 50,000 | 772,766 | 355,190 | 125,376 | 28,242 | 123 | 0.44 |
| 18 | 2009 | CLL | 260,000 | 1,269,550 | 855,820 | 45,634 | 10,476 | 10,430 | 99.56 |
| 18 | 2009 | CLL | 260,000 | 1,418,697 | 930,099 | 51,169 | 11,137 | 11,088 | 99.56 |
| 18 | 2009 | NRB | 130,000 | 898,203 | 614,329 | 33,970 | 9,234 | 9,214 | 99.78 |
| 18 | 2009 | NRB | 130,000 | 1,062,523 | 721,914 | 31,642 | 8,974 | 8,938 | 99.6 |
| 18 | 2012 | CLL | 500,000 | 1,027,101 | 674,666 | 28,127 | 8,403 | 8,371 | 99.62 |
| 18 | 2012 | CLL | 500,000 | 634,488 | 429,098 | 23,154 | 7,011 | 6,965 | 99.34 |
| 19 | 2009 | CLL | 500,000 | 664,361 | 87,219 | 13,780 | 2,494 | 1,749 | 70.13 |
| 19 | 2009 | CLL | 500,000 | 806,969 | 120,890 | 14,360 | 2,666 | 1,598 | 59.94 |
| 19 | 2009 | NRB | 56,000 | 1,039,479 | 73,658 | 17,465 | 2,414 | 608 | 25.19 |
| 19 | 2009 | NRB | 56,000 | 854,462 | 10,110 | 2,386 | 552 | 70 | 12.68 |
| 19 | 2014 | CLL | 500,000 | 1,037,227 | 131,797 | 18,455 | 3,775 | 2,802 | 74.23 |
| 19 | 2014 | CLL | 500,000 | 1,049,570 | 129,588 | 22,198 | 4,168 | 3,661 | 87.84 |
| 20 | 2008 | CLL | 500,000 | 629,340 | 471,880 | 18,929 | 3,028 | 2,778 | 91.74 |
| 20 | 2008 | CLL | 500,000 | 1,243,605 | 949,423 | 30,755 | 4,169 | 3,890 | 93.31 |
| 20 | 2008 | NRB | 46,000 | 599,333 | 15,106 | 7,212 | 304 | 17 | 5.59 |
| 20 | 2008 | NRB | 46,000 | 741,182 | 109,797 | 24,574 | 3,453 | 374 | 10.83 |
| 20 | 2013 | CLL | 500,000 | 1,946,124 | 487,183 | 26,489 | 3,433 | 3,195 | 93.07 |
| 20 | 2013 | CLL | 500,000 | 788,998 | 224,632 | 12,653 | 2,265 | 2,127 | 93.91 |
| 20 | 2018 | CLL | 600,000 | 1,479,956 | 345,434 | 20,108 | 2,987 | 2,723 | 91.16 |
| 20 | 2018 | CLL | 600,000 | 728,864 | 171,433 | 11,431 | 2,040 | 1,909 | 93.58 |

| | | | | | | | | | |
|---|---|---|---|---|---|---|---|---|---|
| 20 | 2018 | NRB | 10,000 | 331,748 | 9,289 | 3,089 | 634 | 52 | 8.2 |
| 20 | 2018 | NRB | 10,000 | 1,006,334 | 34,891 | 11,544 | 1,399 | 320 | 22.87 |
| 21 | 2007 | CLL | 500,000 | 1,050,957 | 254,417 | 23,113 | 5,587 | 5,192 | 92.93 |
| 21 | 2007 | CLL | 500,000 | 843,918 | 199,715 | 19,820 | 4,816 | 4,511 | 93.67 |
| 21 | 2010 | CLL | 500,000 | 8,497 | 142 | 89 | 8 | 1 | 12.5 |
| 21 | 2010 | CLL | 500,000 | 942,431 | 149,640 | 12,927 | 3,135 | 2,929 | 93.43 |
| 21 | 2010 | NRB | 900 | 151,682 | 9,608 | 2,868 | 687 | 18 | 2.62 |
| 21 | 2010 | NRB | 3,000 | 1,766,539 | 90,623 | 21,099 | 3,399 | 98 | 2.88 |
| 22 | 2006 | CLL | 500,000 | 1,031,815 | 213,459 | 17,749 | 1,348 | 1,228 | 91.1 |
| 22 | 2006 | CLL | 500,000 | 1,116,711 | 220,034 | 20,027 | 1,409 | 1,264 | 89.71 |
| 22 | 2006 | NRB | 24,000 | 541,668 | 107,427 | 51,513 | 11,281 | 15 | 0.13 |
| 22 | 2006 | NRB | 24,000 | 854,341 | 187,069 | 76,355 | 18,212 | 13 | 0.07 |
| 22 | 2011 | CLL | 500,000 | 736,172 | 149,292 | 13,989 | 1,144 | 1,069 | 93.44 |
| 22 | 2011 | CLL | 500,000 | 849,802 | 168,262 | 17,160 | 1,214 | 1,122 | 92.42 |
| 23 | 2018 | CLL | 500,000 | 2,050,780 | 319,306 | 29,730 | 5,646 | 5,509 | 97.57 |
| 23 | 2018 | CLL | 500,000 | 965,348 | 414,168 | 29,275 | 6,301 | 6,070 | 96.33 |
| 23 | 2018 | NRB | 65,000 | 792,425 | 95,888 | 13,040 | 3,019 | 2,649 | 87.74 |
| 23 | 2018 | NRB | 65,000 | 1,394,228 | 210,259 | 24,001 | 5,059 | 4,318 | 85.35 |
| 24 | 2018 | CLL | 300,000 | 840,783 | 690,451 | 44,213 | 6,617 | 6,297 | 95.16 |
| 24 | 2018 | NRB | 9,000 | 924,608 | 604,473 | 224,921 | 36,465 | 7 | 0.02 |
| 24 | 2018 | NRB | 9,000 | 673,281 | 503,553 | 186,048 | 29,473 | 8 | 0.03 |
| 25 | 2004 | CLL | 300,000 | 547,693 | 464,802 | 99,460 | 17,492 | 2,159 | 81.31 |
| 25 | 2004 | CLL | 300,000 | 1,496,290 | 1,249,842 | 214,589 | 38,196 | 4,692 | 80.62 |
| 25 | 2004 | NRB | 2,500 | 221,845 | 1,660 | 1,118 | 109 | 10 | 33.03 |
| 25 | 2004 | NRB | 12,500 | 768,251 | 356,830 | 116,929 | 16,011 | 28 | 1.14 |
| 25 | 2006 | CLL | 300,000 | 952,168 | 744,446 | 106,186 | 21,003 | 2,591 | 84.14 |
| 25 | 2006 | CLL | 300,000 | 776,243 | 566,813 | 80,754 | 15,234 | 1,711 | 78.30 |
| 25 | 2006 | NRB | 4,500 | 626,848 | 37,370 | 12,344 | 1,787 | 106 | 41.80 |
| 25 | 2006 | NRB | 8,000 | 1,397,609 | 139,204 | 38,817 | 6,101 | 484 | 50.40 |
| 25 | 2015 | CLL | 300,000 | 1,170,290 | 896,171 | 84,341 | 15,792 | 1,847 | 80.76 |
| 25 | 2015 | CLL | 300,000 | 1,038,582 | 763,290 | 79,477 | 14,528 | 1,773 | 81.11 |
| 25 | 2015 | NRB | 18,000 | 799,450 | 492,768 | 181,351 | 23,277 | 31 | 0.76 |
| 25 | 2015 | NRB | 24,500 | 1,050,113 | 424,300 | 156,012 | 18,815 | 25 | 0.86 |
| 25 | 2017 | CLL | 300,000 | 1,715,138 | 1,236,811 | 101,314 | 17,570 | 2,225 | 85.89 |

| | | | | | | | | | |
|---|---|---|---|---|---|---|---|---|---|
| 25 | 2017 | CLL | 300,000 | 1,364,420 | 980,271 | 87,618 | 15,621 | 2,009 | 85.88 |
| 25 | 2017 | NRB | 4,000 | 932,190 | 63,663 | 17,964 | 2,630 | 203 | 52.70 |
| 25 | 2017 | NRB | 2,000 | 1,007,669 | 36,600 | 11,685 | 1,520 | 106 | 46.32 |
| 26 | 2018 | CLL | 500,000 | 361,005 | 118,753 | 19,024 | 3,825 | 2,117 | 55.35 |
| 26 | 2018 | CLL | 500,000 | 419,311 | 103,103 | 20,115 | 3,864 | 1,896 | 49.07 |
| 26 | 2018 | NRB | 25,000 | 832,351 | 338,168 | 114,961 | 15,139 | 66 | 0.44 |
| 26 | 2018 | NRB | 28,000 | 933,474 | 355,666 | 127,848 | 16,399 | 130 | 0.79 |
| 27 | 2018 | CLL | 300,000 | 638,891 | 573,155 | 36,901 | 9,300 | 6,519 | 70.1 |
| 27 | 2018 | CLL | 300,000 | 782,889 | 688,688 | 49,950 | 11,199 | 7,687 | 68.64 |
| 27 | 2018 | NRB | 26,000 | 839,911 | 663,599 | 100,417 | 19,891 | 7,913 | 39.78 |
| 27 | 2018 | NRB | 27,000 | 809,636 | 629,461 | 105,377 | 20,465 | 7,862 | 38.42 |

[a] Cells were distributed into two aliquots in each PB sampling.
